# Multi-response Mendelian randomization: Identification of shared and distinct exposures for multimorbidity and multiple related disease outcomes

**DOI:** 10.1101/2023.02.01.526689

**Authors:** Verena Zuber, Alex Lewin, Michael G. Levin, Alexander Haglund, Soumaya Ben-Aicha Gonzalez, Costanza Emanueli, Scott Damrauer, Stephen Burgess, Dipender Gill, Leonardo Bottolo

**Author notes:** Correspondence: Verena Zuber and Leonardo Bottolo.

## Abstract

The existing framework of Mendelian randomization (MR) infers the causal effect of one or multiple exposures on one single outcome. It is not designed to jointly model multiple outcomes, as would be necessary to detect causes of more than one outcome and would be relevant to model multimorbidity or other related disease outcomes. Here, we introduce Multi-response Mendelian randomization (MR^2^), a novel MR method specifically designed for multiple outcomes to identify exposures that cause more than one outcome or, conversely, exposures that exert their effect on distinct responses. MR^2^ uses a sparse Bayesian Gaussian copula regression framework to detect causal effects while estimating the residual correlation between summary-level outcomes, i.e., the correlation that cannot be explained by the exposures, and *viceversa*. We show both theoretically and in a comprehensive simulation study how unmeasured shared pleiotropy induces residual correlation. We also reveal how non-genetic factors that affect more than one outcome contribute to their correlation. We demonstrate that by accounting for residual correlation, MR^2^ has higher power to detect shared exposures causing more than one outcome. It also provides more accurate causal effect estimates than existing methods that ignore the dependence between related responses. Finally, we illustrate how MR^2^ detects shared and distinct causal exposures for five cardiovascular diseases in two applications considering cardiometabolic and lipidomic exposures and uncovers residual correlation between summary-level outcomes reflecting known relationships between cardiovascular diseases.

## Introduction

Researchers focus often on understanding, preventing and treating specific health conditions in isolation with a disease-centric approach. Yet, as life expectancy increases the incidence of diseases increases and a growing proportion of the adult population is affected by more than one chronic health condition [1, 2, 3]. Multimorbidity describes the simultaneous presence of two or more chronic conditions in one individual [4]. The Academy of Medical Science considers multimorbidity as a key priority for global health research [5] and the World Health Organization identifies people with multimorbidities at higher risk of patient safety issues [6]. To define effective prevention and intervention strategies, it is important to understand disease aetiology. Recent research into multimorbidity suggests the presence of disease clusters systematically co-occurring in subjects with specific genetic predispositions and exposed to certain exposures [7, 8]. Yet, to date, it is unclear if multimorbidity represents a random co-occurrence of seemingly unrelated individual health conditions without a common cause or if shared causal exposures are underpinning multiple health conditions [3, 9]. Our motivation is to develop principled causal inference methodology to detect shared or distinct causes of multiple related health outcomes using genetic evidence in a novel multitrait Mendelian randomization (MR) framework.

Observational studies may be biased by unmeasured confounding factors and cannot be used to infer causality. MR uses genetic variants as instrumental variables (IVs) [10] to infer the direct causal effect of an exposure on an outcome irrespective of unmeasured confounders [11, 12, 13]. MR has become an important analytical approach to gaining a deeper understanding of how modifiable exposures impact a single disease outcome.

Yet, while there are methods for multivariable MR that can deal with multiple exposures in one joint model [14, 15], to date there is no comprehensive MR methodology that can jointly model multiple outcomes, accounting for information shared between the outcomes, while simultaneously detecting common and distinct causes of disease. Consequently, existing MR methodology largely neglects links between related disease outcomes. For example, we have recently performed wide-angled MR investigations to look at genetic determinants of lipids and cardiovascular disease outcomes [16] and blood lipids and particle sizes as exposures for coronary and peripheral artery disease [17] where we have performed MR analysis for each outcome separately. While this strategy provides a first scan if similar causal exposures are significantly detected across different traits, there is no principled MR methodology available to test and define if an exposure affects more than one outcome. Moreover, existing models ignore information shared between outcomes since each trait is considered in isolation. Thus, novel methods are needed to make full use of the growing knowledge regarding clusters of diseases that may share the same causes. In addition, genome-wide association studies (GWAS) are generally sparsely controlled for confounders, and only a few covariates like age, sex and principal components to control for population stratification are included in the regression model to derive summary-level data for genetic associations. While this strategy reduces the risk of collider bias, there may be potential residual confounding in the GWAS themselves which can be addressed by jointly modelling multiple outcomes.

Here, we propose Multi-Response MR (MR^2^) to model multiple related health conditions in a joint multivariate (multiple outcomes) and multivariable (multiple exposures) MR model. Our motivations are the following. First, we seek to distinguish between exposures which are shared (affecting more than one outcome at the same time) or distinct (affecting only one outcome). Second, our multi-response model aims at increasing the power to detect exposures that affect more than one outcome while effectively reducing the number of false positives. Third, MR^2^ aims at combining information between outcomes to identify the effect of unmeasured pleiotropic pathways on the responses as well as the impact of non-genetic factors (independent of the exposures), such as social health determinants, on the correlation between disease outcomes.

MR^2^ is based on the seemingly unrelated regression model [18] which is defined by a series of multivariable MR models, one for each outcome that are connected by the correlation between the residuals of each MR model. The model is called “seemingly unrelated” because the multivariable MR models for each outcome seem unrelated (independent), but their error terms are correlated. Consequently, MR^2^ stands between an analysis that considers each outcome separately (current MR methodology) and a joint (graphical) model where each outcome is conditioned on the risk factors as well as on all the other outcomes. As an illustrative example of the advantages of the proposed model, we consider two outcomes *Y*_1_ and *Y*_2_ and two exposures *X*_1_ and *X*_2_, where the outcomes are correlated, and *X*_1_ → *Y*_1_ and *X*_2_ → *Y*_2_, respectively. Now, since *Y*_1_ and *Y*_2_ are correlated, separate MR models will likely identify both *X*_1_ and *X*_2_ as causes for both outcomes. However, by modelling the correlation between the residuals, that is the correlation between *Y*_1_ and *Y*_2_ after conditioning on *X*_1_ and *X*_2_, MR^2^ will detect the true causes, that is *X*_1_ → *Y*_1_ and *X*_2_ → *Y*_2_.

As the first motivating example, we want to identify which common cardiometabolic risk factors, including diabetes, dyslipidemia, hypertension, physical inactivity, obesity and smoking, are shared or distinct causes of five cardiovascular diseases, including atrial fibrillation, cardioembolic stroke, coronary artery disease, heart failure and peripheral artery disease. We include these outcomes because there is a priori epidemiological and clinical evidence that they are strongly connected due to shared risk factors and because one outcome causes another. For example, coronary artery disease can cause heart failure [19, 20] and atrial fibrillation [21]. In turn, atrial fibrillation can cause cardioembolic stroke [22]. Consequently, in clinal practice, these cardiovascular diseases are frequently present as multimorbidity [23]. Patients suffering from one disease are more likely to be affected by a second cardiovascular illness than a healthy individual becoming sick with one cardiovascular disease [24]. Epidemiological evidence [25, 26, 27, 28] also suggests that these diseases share a wide range of common exposures. Determining whether these exposures are universally causal or influenced by residual confouding/correlation is challenging to infer from traditional observational study designs. To date, no study has used genetic evidence in a joint multioutcome model to establish which exposures are shared or distinct. Here, we illustrate the advantage of using the proposed joint multivariable and multiresponse MR^2^ model to identify which cardiovascular exposures are shared or distinct for different cardiovascular conditions.

As a second motivating example, we follow up on the findings from the first example to define in more detail which lipid characteristics and lipoprotein related traits, as measured by high-throughput metabolomics, are likely causes of the selected cardiovascular diseases.

The manuscript is outlined as follows. After Material and Methods, where we introduce the Bayesian modelling framework of MR^2^, we present in Results an extensive simulation study. First, we illustrate how residual correlation is caused by unmeasured shared pleiotropy and, second, we compare MR^2^ with existing multivariable, single-outcome MR models and with other statistical learning algorithms for multi-response regression regarding their ability to detect important causal exposures, distinguish between shared and distinct exposure and accurately estimate causal effects. Then, we present the results from the two motivating application examples. In the real examples, we contrast the results obtained by MR^2^ with standard multivariable MR [14] and with MR-BMA, a recently proposed method for single-trait multivariable MR method [15] to highlight the gain of power and the reduction of false positives when multiple responses are jointly analysed. We also compare MR^2^ with multivariable MR-Egger [29] to demonstrate different effects of the unmeasured pleiotropy when dealing with multiple outcomes which cannot be detected by one outcome at-a-time methods. Finally, we conclude with a Discussion and directions for future research.

## Material and Methods

In this section, we illustrate the data input utilised in the proposed method as well as existing MR models including univariable (one exposure and one outcome) and multivariable MR (multiple exposures and one outcome). Then, we describe how multiple outcomes can be modelled jointly by considering the seemingly unrelated regression framework and how this can be generalised by the copula regression model. We show analytically, and demonstrate in the simulation study (see Results), how unmeasured shared pleiotropy affecting more than one outcome can be captured in a multi-response MR model which accounts for the residual correlation between outcomes. Finally, we conclude Material and Methods with an overview of MR^2^ which implements a sparse copula regression model and focuses on the selection of shared and distinct exposures for multiple health conditions. Technical details are presented in Appendix. The Markov chain Monte Carlo (MCMC) implementation of the proposed MR^2^ method is described in Supplemental Information.

### Mendelian Randomization data input

MR^2^ is formulated on summary-level data of genetic association with the exposures and outcomes from large-scale GWAS which are commonly available in the public domain.

According to the two-sample summary-level MR framework, we assume that the genetic associations with the exposure(s) and the genetic associations with the outcome(s) are taken from two distinct cohorts with nonoverlapping samples [30] and are thus *a priori* independent [31]. However, when considering multiple exposures, some of them may be derived from cohorts with full or partially overlapping samples. In this case, since multivariable MR models do account for measured pleiotropy between exposures [14, 32], overlapping samples can be analysed. Besides modelling multiple exposures, MR^2^ also explicitly considers the correlation between responses which, similarly, may be due to outcomes from cohorts with fully or partially overlapping samples.

In summary, the summary-level design facilitates the inclusion of different disease outcomes, as well as exposures. They do not necessarily need to be measured on different independent cohorts since MR^2^ allows for multiple correlated exposures and correlated responses.

### Overview of existing MR models

Standard MR models for summary-level data, both univariable (single exposure) and multivariable (multiple exposures), are formulated as weighted linear regression models where the genetic associations with exposure are regressed against the genetic associations with the outcome. Each genetic variant, used as IV, contributes one data point (or observation) to the regression model which we denote with the index *i*, *i* = 1,…, *n*. For each IV, we take the beta coefficient 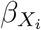 and standard error se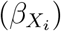 from a univariable regression in which the exposure *X* is regressed on the genetic variant *G_i_* in sample one and beta coefficient 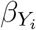 and standard error se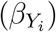 from a univariable regression in which the outcome *Y* is regressed on the genetic variant *G_i_* in sample two.

Then, univariable MR can be formulated as a weighted linear regression model in which the genetic associations with the outcome 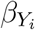 are regressed on the genetic associations with the exposure 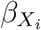 [33]

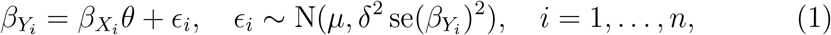

where *θ* is the effect estimate, *μ* is the intercept and *δ*^2^ is an overdispersion parameter, *δ*^2^ ≥ 1, that incorporates residual heterogeneity into the model [34, 35]. Standard MR models set *μ* = 0, while *μ* ≠ 0 captures unmeasured horizontal pleiotropy [36]. Weighting each genetic variant *i* by the first-order weights se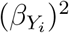 is equivalent to fitting an inverse variance weighting (IVW) MR model which gives genetic variants measured with higher precision larger weights [13]. Alternatively, the genetic associations may be standardized before the analysis by the weights 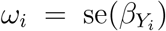, *i* = 1,…, *n*, that only depend on the standard errors of the genetic associations with the outcome.

Multivariable MR [14, 32] is an extension of univariable MR to consider not just one single exposure, but multiple exposures in one joint model. This joint model accounts for measured pleiotropy [14] by modelling explicitly pleiotropic pathways *via* any of the included exposures. Additionally, multivariable MR can be used to select the most likely causal exposures from a set of candidate exposures [15, 37].

In analogy with the univariable MR model in eq. (1), in multivariable MR the genetic associations with one outcome are regressed on the genetic associations with all the exposures [38]

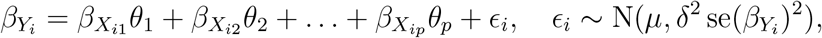

or, in vector notation,

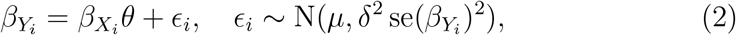

for each *i* = 1,…, *n*, where 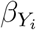 are the associations of the genetic variant *G_i_* with the outcome *Y*, 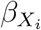 contains the associations of the genetic variant *G_i_* with the *p* exposures, *θ* = (*θ*_1_*,…, θ_p_*)^*T*^ is the vector of the effect estimates, *μ* is the intercept that models unmeasured horizontal pleiotropy and *δ*^2^ > 1 is the overdispersion parameter.

In multivariable MR, a genetic variant is a valid IV if the following criteria hold:

IV1 *Relevance*: The variant is associated with at least one of the exposures.

IV2 *Exchangeability*: The variant is independent of all confounders of each of the exposure-outcome associations.

IV3 *Exclusion restriction*: The variant is independent of the outcome conditional on the exposures and confounders.

Given these assumptions hold, we consider the effect estimates *θ* as the direct causal effect [38] of the exposure on the outcome after keeping all other exposures constant.

### Multi-response MR

We present here the extension of the multivariable MR model in eq. (2) for *p* exposures when *q* outcomes are jointly considered. Consequently, the observed summary-level input data is a matrix *β_Y_* of dimension *n × q* which contains the genetic associations of *n* genetic variants with the *q* outcomes and *β_X_* of dimension *n × p* which includes the genetic associations of the same *n* genetic variants with the *p* exposures.

In the following, we assume that the input data *β_Y_* and *β_X_* have been standardized according to IVW before the analysis and, for easy of exposition, we use the same notation after standardisation. Details on how to define IVW in a multiple outcomes framework are provided in Appendix.

The aim to model *q* related outcomes in one joint model can be achieved by using the seemingly unrelated regression (SUR) framework [18] which is formulated as a series of *q* multivariable regression equations, one for each of the *q* outcomes

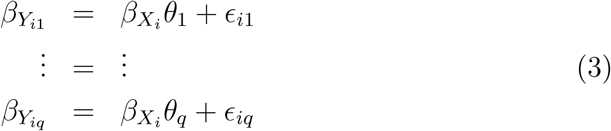

for each *i* = 1,…, *n*, where 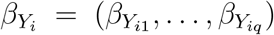 contains the observed genetic associations of the genetic variant *G_i_* with the *q* responses, 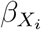 are the associations of the genetic variant *G_i_* with the *p* exposures, *θ*_1_ = (*θ*_11_*,…, θ*_1*p*_)^*T*^, …, *θ_q_* = (*θ_q_*_1_ …, *θ_qp_*)^*T*^ are the outcome-specific vectors of direct causal effects for each outcome and 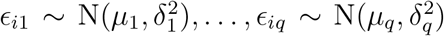 are the residuals with 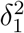*,…, 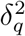* the response-specific intercepts and the overdispersion parameters.

The SUR model connects the *q* multivariable regressions in eq. (3) by allowing for correlation between the *q* residuals of the summary-level outcomes *ϵ_k_*, *k* = 1*,…, q*. More precisely, the SUR model estimates the (*q* × *q*)-dimensional covariance matrix between the vector of residuals *ϵ_i_* = (*ϵ_i_*_1_*,…, ϵ_iq_*)

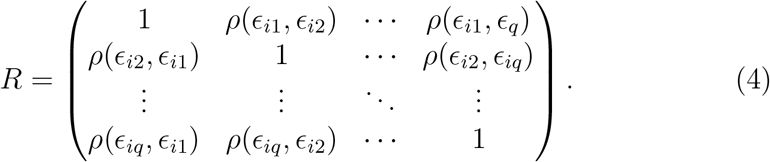

For instance, *ρ*(*ϵ_ik_, ϵ_ik_′*) is the correlation between the residuals of the multivariable MR model for summary-level outcome *k* and the residuals of the multivariable MR model for summary-level outcome *k*′, *k* ≠ *k*′ = 1*,…, q*.

Finally, the possibility to account for response-specific unmeasured horizontal pleiotropy can be turned off by setting the vector of intercepts *μ* = (*μ*_1_*,…, μ_q_*)^*T*^ to zero.

### Correlation between outcomes in multi-response MR

To understand what contributes to the residuals *ϵ_k_*, *k* = 1*,…, q*, of the summary-level MR model in eq. (3) and generates correlation between them in eq. (4), we focus on the following generating model for the *k*th outcome on individual-level data considering *N* subjects, as illustrated in Figure 1A,

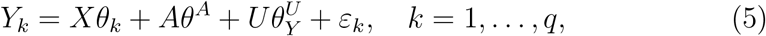

where *θ_k_* is the *p*-dimensional vector of effects of the exposures *X* on the *k*th outcome, *θ^A^* is the effect of the unmeasured pleiotropic pathway *A* on the responses, 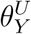 is the effect of the unmeasured confounder *U* on the outcomes and 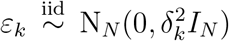 with 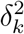 the response-specific residual variance and *N* the sample size. Moreover, *X*, *A* and *U* are random quantities that are descendants of the same set of IVs.

**Figure 1.**
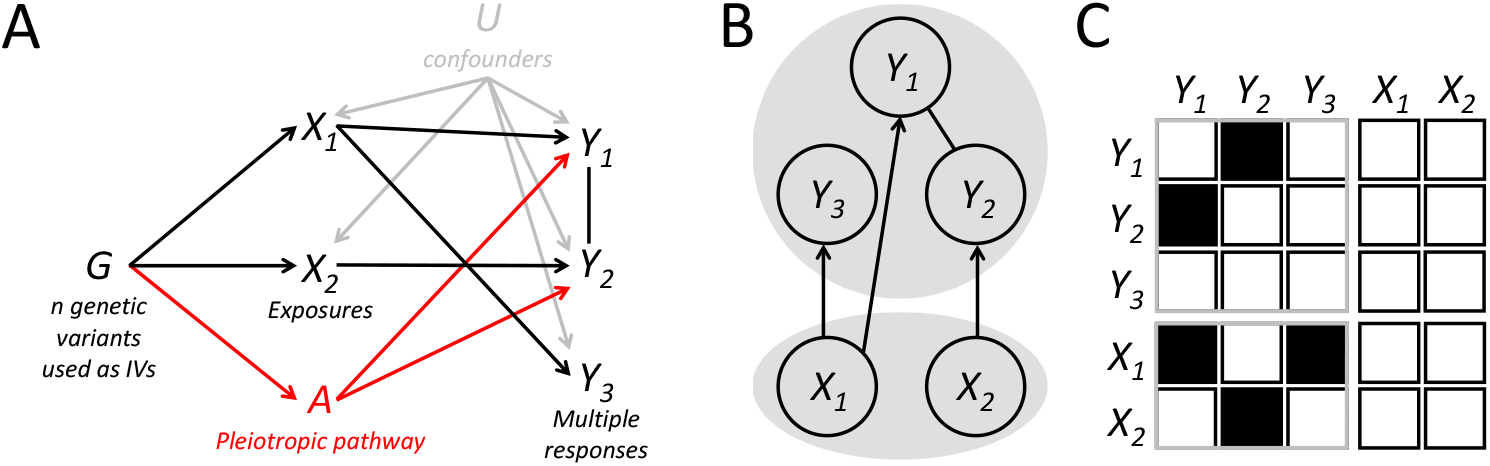
Multivariable and multi-response Mendelian randomisation with multiple exposures and multiple responses. (**A**) Directed acyclic graph (DAG) with *G*: Genetic variant(s), *X* = {*X*_1_, *X*_2_} : Exposures, *Y* = {*Y*_1_, *Y*_2_, *Y*_3_} : Responses, *U* : Unmeasured confounders, *A*: Unobserved pleiotropic pathway. (**B**) Schematic representation of the MR^2^ model which allows the exploration of the model space consisting of all possible subsets of exposures (bottom grey circle) directly associated with the responses (top grey circle) while estimating the residual correlation between the responses (top grey circle) and *viceversa*. Model depicted in (A) is reported in MR^2^ as follows: *X*_1_ is a shared exposure for outcomes *Y*_1_ and *Y*_3_ while *X*_2_ has a distinct direct causal effect on *Y*_2_ (directed edge). Residual dependence between *Y*_1_ and *Y*_2_ is still present after conditioning on the associated exposures (undirected edge) and it depends on the unmeasured pleiotropic pathway *A*. (**C**) MR^2^ model space exploration is equivalent to learning from the summary-level input data a partitioned non-symmetrical adjacency matrix. The model depicted in (A) is represented by an adjacency matrix describing the conditional dependence structure among the responses (top left symmetrical submatrix) and the direct causal association of the exposures with the outcomes (bottom left non-symmetrical submatrix). No reverse causation is allowed in the MR^2^ model (top right non-symmetrical submatrix) and the exposures can only have direct causal effects on the responses, *i.e*., no direct effects among the exposures are modelled (bottom right symmetrical submatrix).

Assuming a quantitative outcome *Y_k_* measured on *N* individuals, the *n*-dimensional vector of summary-level genetic associations with the *k*th response 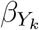 can be derived using the ordinary least squares (OLS) estimate

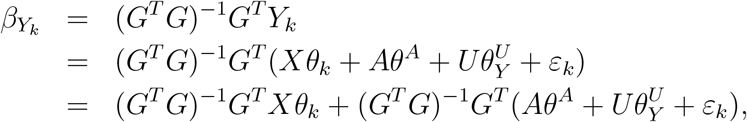

where the (*N × n*)-dimensional matrix *G* describes the *n* genetic variants associated with the exposures and measured on *N* individuals. Moreover, assuming that the genetic variants *G* selected as IVs are independent of each other (achieved by pruning) and independent of the confounder *U* (exchangeability assumption), the above equation simplifies to

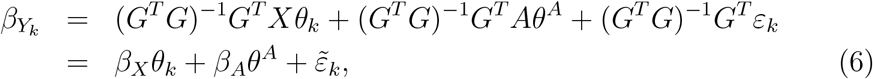

where *β_X_* = (*G^T^ G*)^−1^*G^T^ X* is the (*n × p*)-dimensional matrix of the summary-level genetic association with the measured exposures *X*, *β_A_* = (*G^T^ G*)^−1^*G^T^ A* is the *n*-dimensional vector of genetic associations with the unmeasured pleiotropic pathway *A* and 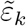 can be viewed as the OLS of the projection of *∊_k_* onto the linear space spanned by *G*. Note that 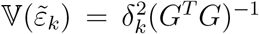 and, given the assumption of independence of the genetic variants, it simplifies to 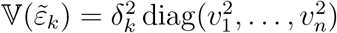, where diag(·) indicates a diagonal matrix. Thus, 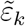 is a *n*-dimensional vector and 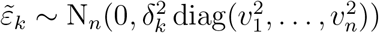.

Consequently, the residuals in eq. (3) for the *k*th summary-level response, *k* = 1,…, *q*, using eq. (6), can be decomposed into

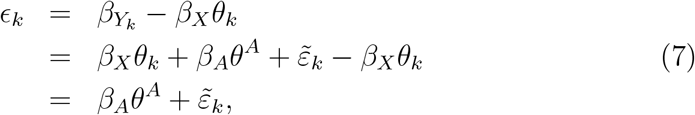

which shows that residuals in the summary-level MR model reflect unmeasured shared pleiotropy and residual variation.

Since in the designed framework the unmeasured pleiotropic pathway *A* depends on *G* and is random, *β_A_* in eq. (7) is also random and, thus, *β_A_θ^A^* cannot be treated as the fixed-effect response-specific intercept *μ_k_* [29]. Instead, it should be interpreted as a random intercept common to all summary-level responses. Similarly to random-effect models, the distribution of *ϵ_k_* follows a *n*-dimensional Gaussian distribution with 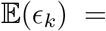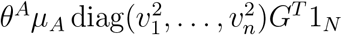 with 1_*N*_ a *N*-dimensional vector of ones, and 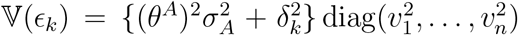, assuming *A* and *∊_k_* independent, E(*A*) = *μ_A_*1_*N*_ and the identity matrix of dimension *N*. Moreover, it reveals that the residual correlations in eq. (4) depend on the unmeasured shared pleiotropy, as illustrated below for two outcomes *k* and *k*′, *k ≠ k*′,

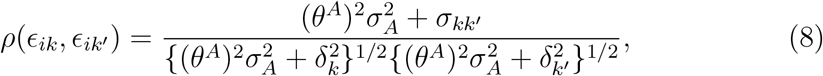

where *σ_kk_′* = Cov(*∊_k_, ∊_k_′*) is the covariance between individual-level responses’ errors in eq. (5) which is assumed constant across all *N* individuals.

Eq. (8) shows that, independently of the sign of *θ^A^* (the effect of the unmeasured pleiotropic pathway *A* on the responses) and the nature of the shared pleiotropy *A* (either “directed”, *i.e*., *A_l_* > 0, or “undirected”, *i.e*., *A_l_* ≷ 0, *l* = 1,…, *n*) the effect of unmeasured shared pleiotropy on the correlation between the residuals of the multivariable MR model is always positive and constant across all combinations of responses. Moreover, the residual correlation is different from zero even when Cov(*∊_k_, ∊_k_′*) = 0, *i.e*., there is no correlation between individual-level responses’ errors.

Alternative scenarios can be also considered. As illustrated in Figure 1A, some outcomes may not be influenced by the unmeasured pleiotropic pathway, so their summary-level residual correlation with other responses is nil. Likewise, there may exist several unmeasured pleiotropic pathways which are shared by subgroups of responses (not necessarily distinct). For instance, pathway *A*_1_ affects responses 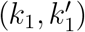, 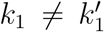, and *A*_2_ impacts responses 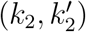, 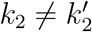. In these cases, the derivation of eq. (8) does not change, although *ρ*(*ϵ_ik_, ϵ_ik_′*) won’t be constant across all outcomes’ pairs. Full details of the derivation of the above quantities are presented in Supplemental Information.

Finally, we comment on *σ_kk_′* in eq. (8) which is the covariance between individual-level responses’ errors. In contrast to the correlation induced by unmeasured pleiotropic pathway *A*, which is genetically proxied by the selected IVs, *σ_kk_′*, *k* = *k*′ = 1*,…, q*, does not depend on *G*. Examples of non-genetic factors that contribute to the outcomes’ correlations are, for instance, social health determinants [39] such as personal features, socioeconomic status, culture, environment, health behaviors, access to care and government policy. Notably, these interconnected factors that determine an individual’s health status are not associated with the exposures and, therefore, are not confounders. This echoes the InSIDE (INstrument Strength Independent of Direct Effect) assumption [36], where the pleiotropic effect is independent of the genetic associations with the exposure albeit, here, it induces correlation between responses. Moreover, while further exposures can be included in the analysis with the hope to catch the effects of unmeasured shared pathways, the residual correlations generated by non-genetic factors cannot be “explained away” in the current MR framework. Thus, it should be considered as the baseline residual correlation between summary-level outcomes.

### Bayesian multi-response MR

MR^2^ is based on a recently proposed Bayesian method to select important predictors in regression models with multiple responses of any type [40]. Specifically, a sparse Gaussian copula regression (GCR) model [41] is used to account for the multivariate dependencies between the Gaussian responses *β_Y_* once their direct causal association with a set of important exposures *β_X_* is estimated. When only Gaussian responses are considered, the GCR is similar to the seemingly unrelated regression (SUR) model [18] (see Appendix). Figure 1B provides a schematic representation of the MR^2^ model which allows for the estimation of important exposures (bottom grey circle) directly associated with the responses (top grey circle) while estimating the residual correlation between the responses (top grey circle) and *viceversa*.

Regarding MR^2^ model specification, we use the hyper-inverse Wishart distribution as the prior density for the residual covariance matrix based on the theory of Gaussian graphical models [42]. This prior allows some of the off-diagonal elements of the inverse covariance matrix to be identical to zero and to estimate the residual correlation between the summary-level responses. For the exposures, we use a hierarchical non-conjugate model [43] to assign a prior distribution independently on each direct causal effect. A point mass at zero is specified on the regression coefficient of the null exposure whereas a Gaussian distribution is assigned to the non-zero effect.

From a computational point of view, we design a novel and efficient proposal distribution to update jointly the latent binary vectors for the selection of important exposures (selection step) and the corresponding non-zero effects (estimation step). For Gaussian responses, the designed proposal distribution allows the “implicit marginalisation” of the non-zero effects in the Metropolis-Hastings (M-H) acceptance probability [44], which makes our MCMC algorithm more efficient than existing sparse Bayesian SUR models [45, 46, 47] favoring the exploration of important combinations of exposures, *i.e*., the model space, in a very efficient manner (see Supplemental Information). Figure 1C shows that MR^2^ model exploration is equivalent to learning from the input data a non-symmetrical adjacency matrix partitioned into a symmetrical submatrix (top left) which describes the conditional dependence structure among the responses and a non-symmetrical submatrix (bottom left) representing the direct causal association of the exposures with the outcomes. Note that neither reverse causation (top right submatrix) nor direct causal effects between the responses (bottom right submatrix) are allowed in the MR^2^ model.

Two sets of parameters are deemed important in our analysis: The marginal posterior probability of inclusion (mPPI) which measures the strength of the direct causal association between each exposure-response combination, and the corresponding direct causal effect, and the edge posterior probability of inclusion (ePPI) which describes the strength of the residual dependence between each pair of summary-level responses, and the corresponding residual partial correlation. The posterior distribution of these quantities is usually summarised by their mean or any relevant quantile. For instance, the *α*/2% and (1 − *α*/2)% quantiles are used to build the (1 − *α*)% credible interval. We summarise all quantities of interest by their posterior mean (both mPPI and ePPI can be seen as a posterior mean, or frequency that an exposure-response combination or a pair of dependent responses are selected during the MCMC algorithm) and their 95% credible interval.

The selection of important exposures for each response and significantly correlated pairs of responses is based on mPPIs and ePPIs, respectively. Thresholding these quantities at 0.5 is usually suggested given that the optimal predictive model in linear regression is often the median probability model, which is defined as the model consisting of those predictors which have overall mPPI ≥ 0.5, the optimal predictive model under square loss, *i.e*., the optimal model that predicts not yet collected data [48]. Here, we follow a different approach based on in-sample selection. A non-parametric False Discovery Rate (FDR) strategy based on two-component mixture models [49, 50] which clusters low and high levels of mPPIs, and low and high levels of ePPIs, is applied to select important exposures for each response and significant dependence patterns among responses at a fixed FDR level.

Importantly, for the definition of exposures causing more than one outcome, the availability of the latent binary vectors for the selection of important exposures for each response recorded during the MCMC algorithm allows also the estimation of the joint posterior probability of inclusion (jPPI) defined as the number of times an exposure is selected to be associated with two or more responses at the same time during the MCMC. Thus, jPPI can be regarded as the joint probability of inclusion for any combination of responses. The detection of important direct causal effects of an exposure on a single or multiple responses is carried out by looking at significant mPPIs selected at a nominal FDR level. If an exposure is associated with more than one response, we declare the existence of a shared direct causal effect and calculate the jPPIs.

While the residual correlation between summary-level responses captures “global” unmeasured shared pleiotropy, which is calculated across all genetic variants, we additionally screen for individual genetic variants as potential outliers due to their “local” pleiotropic effect [51]. Building on Bayesian tools for outlier diagnostics, we propose the conditional predictive ordinate (CPO) [52] to detect individual genetic variants as outliers or high-leverage and influential observations in the MR^2^ model.

Full details of the MR^2^ model as well as post-processing of the MCMC output are presented in Appendix.

## Results

### Simulation study: Can the effect of shared pleiotropy be detected?

Here, we conduct a simulation study to illustrate the impact of unmeasured shared pleiotropy affecting more than one outcome. We consider one of the scenarios presented in the simulation study (Scenario III-Undirected pleiotropy) where the residual correlation between outcomes at the summary-level depends only on the unmeasured shared pleiotropy and *σ_kk_′* = 0 for all responses. We look at two quantities. First, the empirical correlations between the summary-level genetic associations of the outcomes measured on the IVs. They can be computed by simply calculating the pairwise correlation between the genetic associations with the outcomes on the summary-level. Second, the residual correlation between the summary-level genetic associations of the outcomes measured on the IVs after accounting for the exposures and estimated by MR^2^.

Figure 2 shows that the larger the pleiotropic effect *θ^A^* (ranging from 0.25 to 2), the bigger the empirical correlation. However, the empirical correlation (in grey) is not able to distinguish between the correlation due to the direct causal effects of the exposures on the outcomes and the unmeasured shared pleiotropy. MR^2^ can separate the source of correlation with a good agreement between the estimated values of the residual correlation (in red) and its theoretical value derived in eq. (8) (black dashed line).

**Figure 2.**
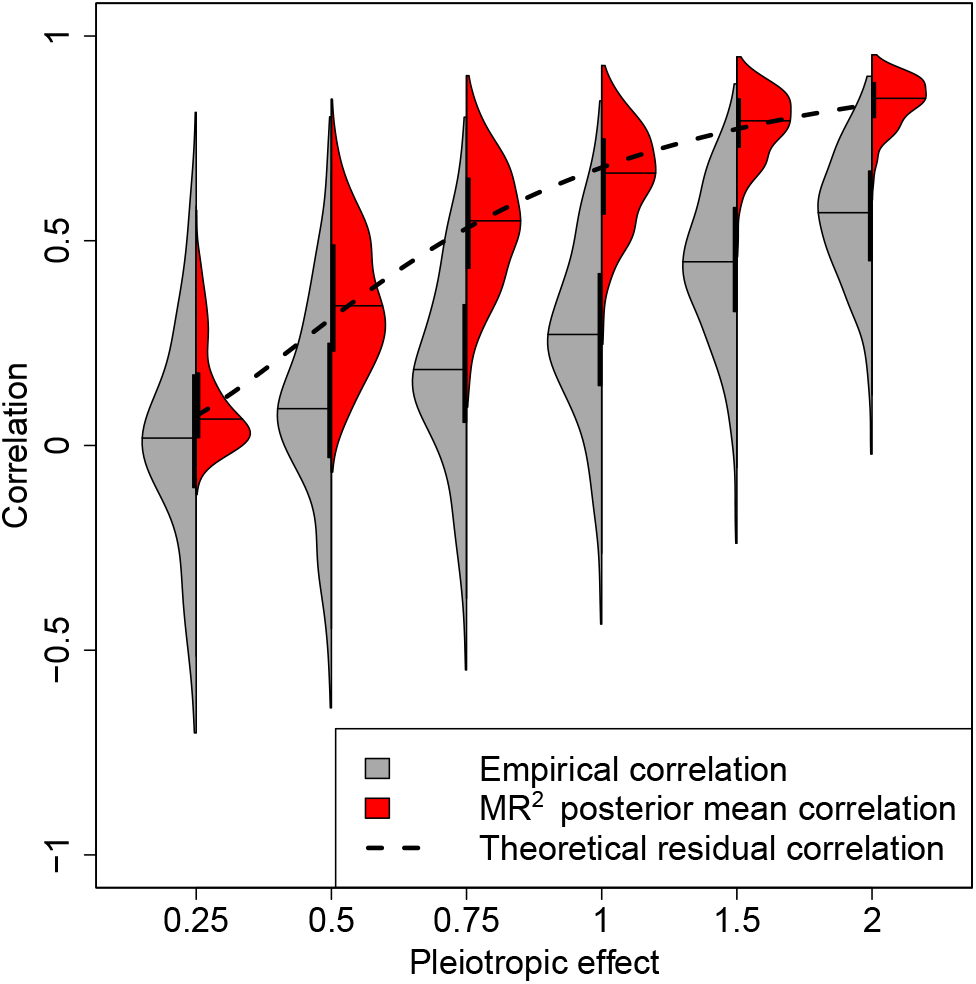
Correlations between the summary-level genetic associations with the outcomes induced by various levels of the pleiotropic effect. Each violin plot depicts the empirical correlations between the summary-level genetic associations of the outcomes (dark grey) and the posterior mean of the residual correlations estimated by MR^2^ (red) in simulation Scenario III-Undirected pleiotropy averaged over 50 replicates at different levels of the pleiotropy effect *θ^A^* = {0.25, 0.50, 0.75, 1.00, 1.50, 2.00}. Confounding effects on the exposures and outcomes are fixed at 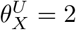 and 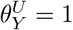1, respectively, without correlation between exposures and responses’ errors. For each side of the violin plot, the vertical black thick line displays the interquartile range, while the black line indicates the median. Black dashed line depicts the theoretical value of residual correlation between summary-level outcomes as a function of *θ^A^* as shown in eq. (8). For more details on the simulation setting, see Supplemental Table S.2.

### Simulation study: Comparison of methodologies

We perform a simulation study to demonstrate the increase in power, a better estimation of direct causal effects when accounting for multiple correlated outcomes and finally the ability to detect shared and distinct causal exposures when using the proposed MR^2^ method compared to existing multivariable MR models considering one outcome at-a-time and recently proposed multi-response multivariable methods. In total, we simulate *n* = 100 genetic variants used as IVs, *p* = 15 exposures and *q* = 5 outcomes. Out of the *p×q* exposure-outcome combinations, 30% have a non-zero direct causal effect. For all scenarios, we generate *N* = 100, 000 individuals of which half is used to generate the summary-level genetic associations for the exposure and half for the outcomes, respectively, providing data in a two-sample summary-level design. As alternative methods, we include standard multivariable MR (MV-MR) [38] and MR-BMA (Mendelian randomization with Bayesian model averaging) [15], a Bayesian variable selection approach for multivariable MR. We consider two multivariable and multivariate variable selection approaches which have to date not been applied to MR, the multiple responses Lasso (Multivariate Regression with Covariance Estimation, MRCE) [46] and the multiple responses Spike and Slab Lasso (mSSL) [47]. Both methods perform variable and covariance selection by inducing sparsity and setting the effect estimates of variables not included in the model to zero, as well as inducing sparsity in the residual covariance matrix. An overview of the alternative methods is provided in Supplemental Table S.1.

We simulate the following scenarios:

I - *Null* : There are no causal exposures for any of the outcomes and no confounder.
II - *Confounding* : There are 30% of exposures with a non-zero direct causal effect and there is a joint confounder for all outcomes.
III - *Undirected pleiotropy*: Residual correlation between outcomes is induced by a shared undirected pleiotropic pathway which can increase or decrease the level of the responses.
IV - *Directed pleiotropy* : Residual correlation between outcomes is induced by a shared directed pleiotropic pathway which only increases the level of the responses.
V - *Dependence*: Outcomes are simulated with correlated errors mimicking the effect of non-genetic factors that contribute to their correlation.

Full details regarding the simulation study setup are presented in Appendix. An overview of the simulations setting and the open parameters (θ and 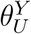 in eq. (5),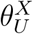, the effect of the unmeasured confounder *U* on the exposures, and *r_Y_*, the correlation between individual-level responses’ errors in eq. (5)) that vary across the simulation scenarios are shown in Supplemental Table S.2. In Supplementary Information, we also provide details regarding MR^2^ hyper-parameters setting, including the number of MCMC iterations after burn-in, as well as technical details of the alternative methods and their software implementations we used in the simulation study.

The performance in terms of exposure selection is evaluated using the receiver operating characteristic (ROC) curves where the true positive rate (TPR) is plotted against the false positive rate (FPR). As a baseline, all methods perform equally well in the case of no correlation between exposures as seen in Supplemental Figure S.1. In contrast, when the exposures are correlated, all variable selection approaches improve over the standard MV-MR as seen in Figure 3. This improvement depends on the correlation between exposures as shown in Supplemental Figure S.2.

**Figure 3.**
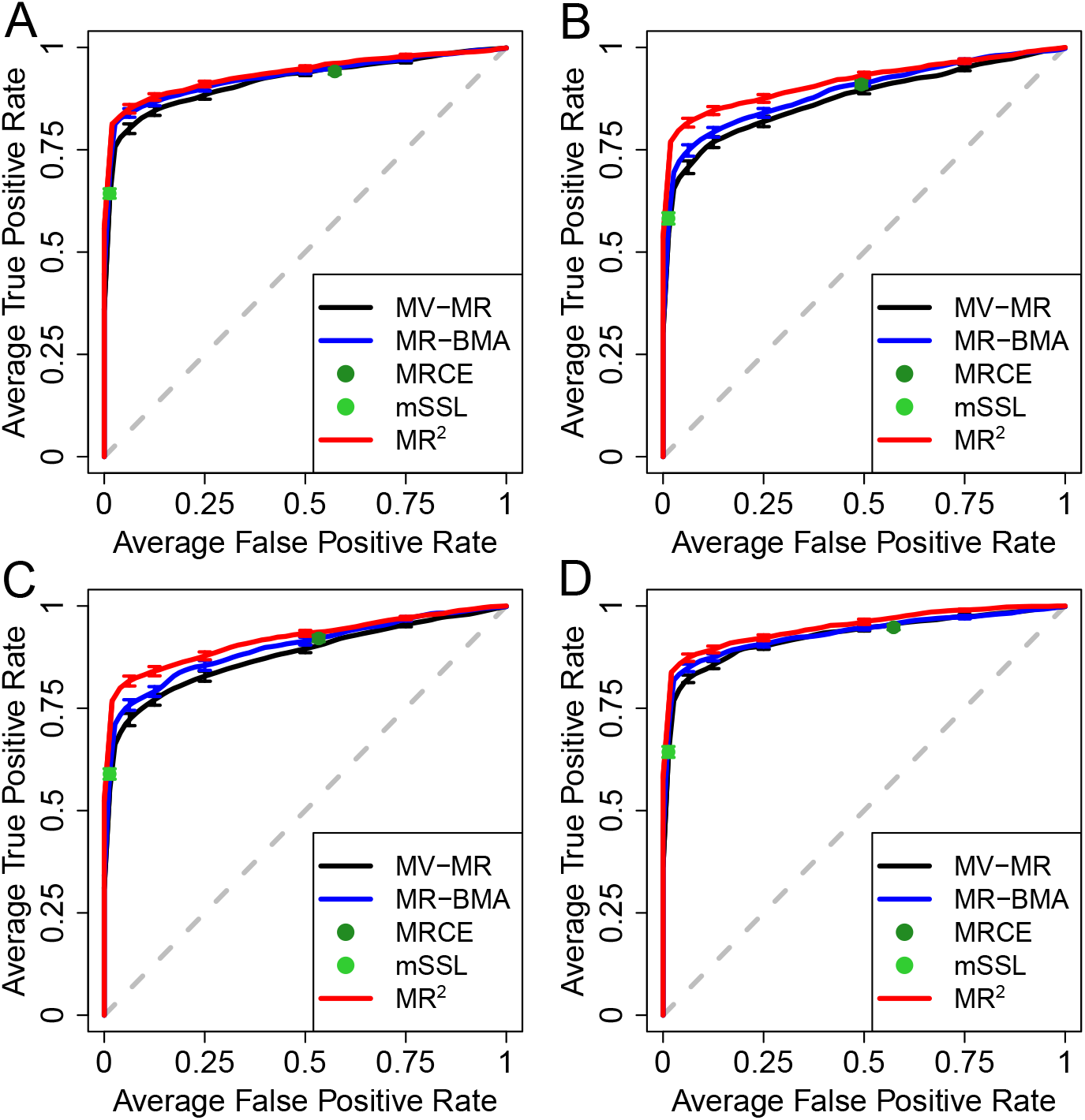
Receiver operating characteristic (ROC) curves in different simulated scenarios when the correlation between exposures is set at *r_X_* = 0.6, averaged over 50 replicates, illustrating the performance of different MR implementations and multi-response multivariable methods to distinguish between true and false causal exposures for five simulated outcomes by plotting the true (TPR) against the false positive rate (FPR). (**A**) depicts baseline Scenario II-Confounding where only the confounding effects on the exposures and outcomes, 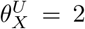 and 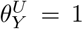, respectively, are used to simulate the data. Residual correlation induced by shared undirected pleiotropy (Scenario III-Undirected pleiotropy) and shared directed pleiotropy (Scenario IV-Directed pleiotropy) with pleiotropic effect set at *θ^A^* = 1 are shown in (**B**) and (**C**), respectively. (**D**) displays the performance of the different methods in Scenario IV-Dependence where the correlation between individual-level responses’ errors is fixed at *r_Y_* = 0.6. Vertical bars in each ROC curve, at specific FPR levels, indicate standard error across 50 replicates. For more details on the simulations setting, see Supplemental Table S.2.

In the following, we set the correlation between exposures at *r_X_* = 0.6 while the confounding effects on the exposures and outcomes are fixed at 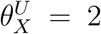 and 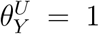, respectively. In Scenario III-Undirected pleiotropy and Scenario IV-Directed pleiotropy, the pleiotropic effect is set at *θ^A^* = 1. Finally, in Scenario V-Dependence, the correlation between individual-level responses’ errors in (5) is fixed at *r_Y_* = 0.6.

When residual correlation is induced by shared undirected (Figure 3B) and directed pleiotropy (Figure 3C), MR^2^ shows a better detection of true causal exposures than MR-BMA, which in turn improves over a basic MVMR model. As shown in Figure 4 for shared undirected pleiotropy and Supplemental Figure S.3 for shared directed pleiotropy, the degree of improvement depends on the strength of the pleiotropic pathway effect. In this scenario, the mSSL approach provides strong sparse solutions with many direct causal effects set to zero. In contrast, MRCE includes many more exposures in the model at the cost of a high false positive rate. Similar results are observed when the residual correlation between outcomes is induced directly through the correlation between individual-level responses’ errors (Figure 3D) with the degree of improvement depending on the level of correlation (Supplemental Figure S.4). For these scenarios, the corresponding area under the ROC curve (AUC) with standard error across 50 replicates, is provided in Table 1. The AUCs of all simulated scenarios across the full range of open parameters are reported in Supplemental Tables S.3-S.6.

**Figure 4.**
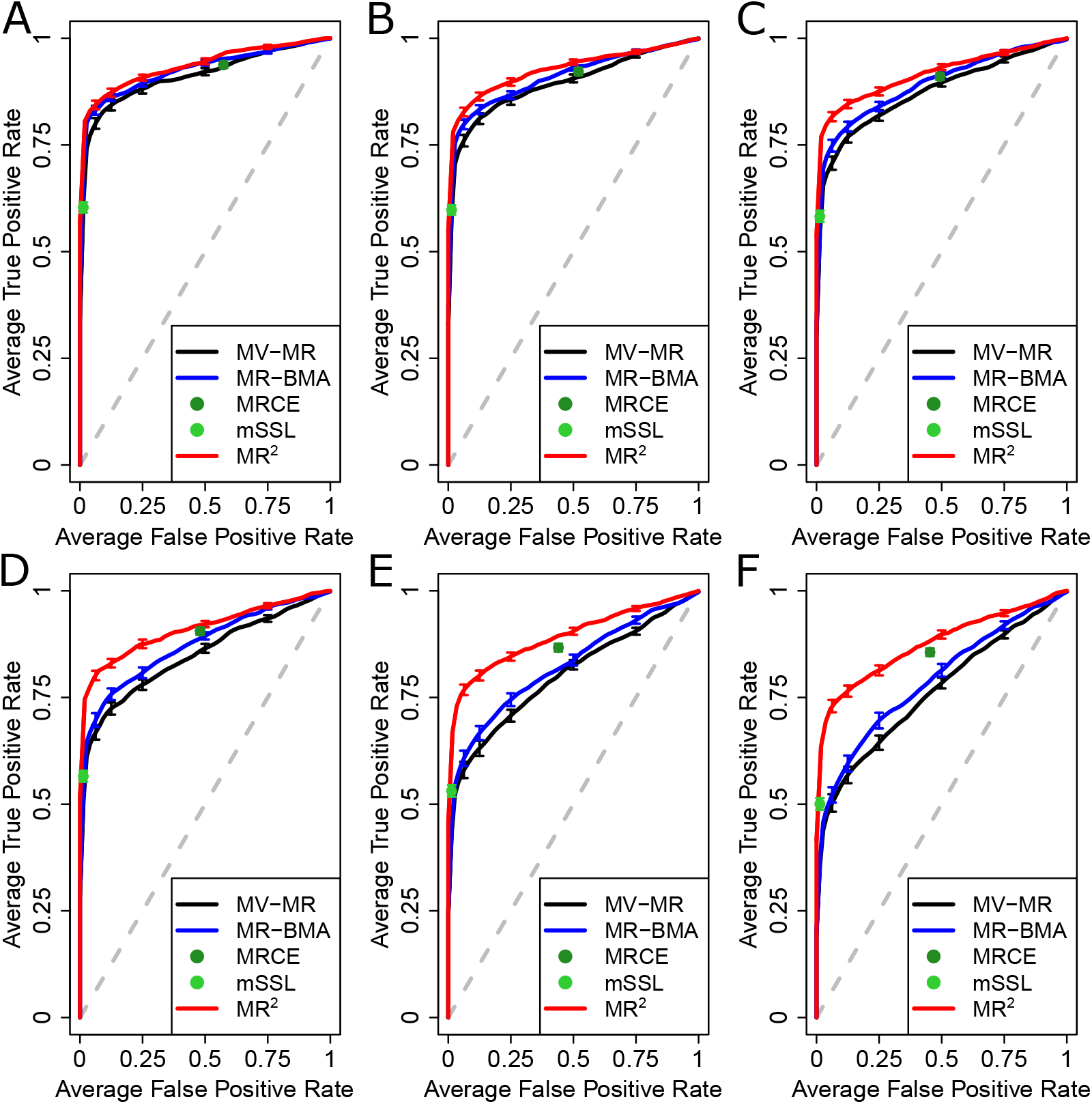
Receiver operating characteristic (ROC) curves for different levels of the pleiotropic pathway effect *θ^A^* and when the correlation between exposures is set at *r_X_* = 0.6, averaged over 50 replicates, illustrating the performance of different MR implementations and multi-response statistical methods to distinguish between true and false causal exposures for five simulated outcomes by plotting the true (TPR) against the false positive rate (FPR). Pleiotropic pathway effect varies from (**A**) to (**F**) with values *θ^A^* = {0.25, 0.5, 0.75, 1, 1.5, 2}. Confounding effects on the exposures and outcomes are fixed at 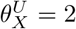 and 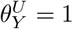, respectively. Vertical bars in each ROC curve, at specific FPR levels, indicate standard error across 50 replicates.

**Table 1.** Area under the curve (AUC) in different simulated scenarios when the correlation between exposures is set at *r_X_* = 0.6. The ability to distinguish between true and false causal exposures is evaluated by the area under the ROC curve averaged over 50 replicates with standard deviation in brackets. In baseline Scenario II-Confounding, only the confounding effects on the exposures and outcomes, 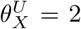 and 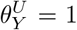, respectively, are used to induce residual correlation between the responses. Residual correlation induced by shared undirected pleiotropy and shared directed pleiotropy with pleiotropic effect *θ^A^* = 1 are presented in Scenario III-Undirected pleiotropy and Scenario IV-Directed pleiotropy, respectively. In Scenario IV-Dependence, outcomes are simulated with the correlation between individual-level responses’ errors fixed at *r_Y_* = 0.6. Best results are highlighted in bold. For more details on the simulations setting, see Supplemental Table S.2.

|  | Scenario II<br>Confounding | Scenario III<br>Undirected<br>pleiotropy | Scenario IV<br>Directed<br>pleiotropy | Scenario V<br>Dependence |
| --- | --- | --- | --- | --- |
| MV-MR | 0.922 (0.040) | 0.850 (0.059) | 0.857 (0.057) | 0.930 (0.037) |
| MR-BMA | 0.932 (0.034) | 0.875 (0.050) | 0.876 (0.058) | 0.936 (0.032) |
| MRCE | 0.926 (0.035) | 0.896 (0.046) | 0.904 (0.048) | 0.932 (0.037) |
| mSSL | 0.876 (0.044) | 0.834 (0.068) | 0.840 (0.055) | 0.869 (0.067) |
| MR<sup>2</sup> | **0.939** (0.033) | **0.912** (0.042) | **0.917** (0.043) | **0.952** (0.031) |

Next, we compare methods according to their performance in estimating the direct causal effect strength. This is evaluated by the sum of squared error (SSE) presented in Table 2 and Supplemental Tables S.7-S.11. As can be seen from Table 2, all methods show a comparable performance with negligible SSE in Scenario I-Null when there are no causal exposures. In the other scenarios, when there is correlation between the exposures, the largest SSE is consistently seen for the MV-MR approach. This is in keeping with other simulation studies [15] that have shown that MV-MR is unbiased but suffers from large variance. MR^2^ has almost everywhere the lowest SSE compared to alternative methods as well as the lowest standard error. The largest improvement with respect to MR-BMA is in Scenario III-Undirected pleiotropy and Scenario IV-Directed pleiotropy when the summary-level outcomes are correlated by a shared pleiotropic pathway. The multi-response implementations MRCE and mSSL perform roughly as MR-BMA in terms of SSE but suffer either from too little (mSSL) or too much (MRCE) sparsity which biases the direct causal effect estimates. Notably, MR-BMA performs better than both MRCE and mSSL in Scenario V-Dependence across all range of open parameters (Supplemental Table S.11) since the two multi-response multivariable methods are not able to identify the simulated correlation pattern between the responses’ errors, thus degrading the estimation of the direct causal effects. Instead, MR^2^ can detect it, resulting in the lowest SSE across the full range of open parameters.

**Table 2.** Sum of squared errors (SSE) and respective standard error in brackets in different simulated scenarios when the correlation between exposures is set at *r_X_* = 0.6. The quality of the direct causal effect estimation is evaluated by SSE, averaged over 50 replicates, with standard deviation in brackets. Scenario I-Null is simulated with the correlation between individual-level responses’ errors fixed at *r_Y_* = 0.6. In baseline Scenario II-Confounding, only the confounding effects on the exposures and outcomes, 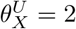 and 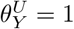, respectively, are used to induce residual correlation between the responses. Residual correlation induced by shared undirected pleiotropy and shared directed pleiotropy with pleiotropic effect *θ^A^* = 1 are presented in Scenario III-Undirected pleiotropy and Scenario IV-Directed pleiotropy, respectively. In Scenario IV-Dependence, outcomes are simulated with the correlation between individual-level responses’ errors fixed at *r_Y_* = 0.6. Best results are highlighted in bold. For more details on the simulations setting, see Supplemental Table S.2.

|  | Scenario I<br>Null | Scenario II<br>Confounding | Scenario III<br>Undirected<br>pleiotropy | Scenario IV<br>Directed<br>pleiotropy | Scenario V<br>Dependence |
| --- | --- | --- | --- | --- | --- |
| MV-MR | 0.001 (<0.001) | 1.246 (0.490) | 3.869 (1.502) | 3.613 (1.409) | 1.200 (0.464) |
| MR-BMA | **0.000** (<0.001) | 0.673 (0.302) | 1.714 (1.036) | 1.508 (0.663) | 0.700 (0.348) |
| MRCE | 0.001 (<0.001) | 0.904 (0.516) | 1.434 (0.877) | 1.172 (0.618) | 0.849 (0.423) |
| mSSL | **0.000** (<0.001) | 0.881 (0.374) | 1.104 (0.790) | 1.050 (0.700) | 0.891 (1.167) |
| MR<sup>2</sup> | **0.000** (<0.001) | **0.567** (0.269) | **0.858** (0.722) | **0.821** (0.681) | **0.521** (0.278) |

Finally, we assess the ability of MR^2^ to detect shared and distinct direct causal effects. To this aim, we calculate the proportion of significant causal effects associated with either single or multiple outcomes in the same scenarios presented in Tables 1 and 2, where the correlation between exposures is fixed at *r_X_* = 0.6. For a fair comparison, we fix the type I error to be the same in all the methods and, in particular, we set it at the level detected by mSSL, given its sparse Lasso solutions with a low false positive rate. Specifically, we selected the threshold of Benjamini-Hochberg False Discovery Rate (FDR) procedure [53] for MV-MR and mPPI for MR-BMA and MR^2^, respectively, to match the number of false positives detected by mSSL. This was not possible for MRCE given the fixed solution of the multiple responses Lasso. Table 3 shows the power of the methods considered. MR^2^ is the most powerful method to detect distinct direct causal effects, which is also the most likely simulated case (on average 37.3% of all exposure-response combinations), across all simulated scenarios. When the residual correlation between summary-level outcomes is simulated (Scenario III-Undirected pleiotropy, Scenario IV-Directed pleiotropy and Scenario V-Dependence) MR^2^ is also the most powerful method to detect shared direct causal effects. Similar results are also obtained for different levels of correlation between individual-level exposures (data not shown).

**Table 3.** Power to detect distinct and shared direct causal effects at a common false positives rate in different simulated scenarios when the correlation between exposures is set at *r_X_* = 0.6 over 50 replicates with standard deviation in brackets. Simulated exposure effects are either distinct, where an exposure has only a causal association with one outcome or shared, where the exposure exerts its effect on multiple (2,3,4) responses (Shared = 5 was not simulated). The average simulated proportion indicates the simulated proportion (over all exposure-response combinations) that an exposure is associated with one (distinct) or multiple (shared) responses. In baseline Scenario II-Confounding, only the confounding effects on the exposures and outcomes, 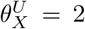 and 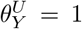, respectively, are used to induce residual correlation between the responses. Residual correlation induced by shared undirected pleiotropy and shared directed pleiotropy with pleiotropic effect *θ^A^* = 1 are presented in Scenario III-Undirected pleiotropy and Scenario IV-Directed pleiotropy, respectively. In Scenario IV-Dependence, outcomes are simulated with the correlation between individual-level responses’ errors fixed at *r_Y_* = 0.6. Best results are highlighted in bold. MRCE method is not considered since it is not possible to match its type I error with the other methods. For more details on the simulations setting, see Supplemental Table S.2.

| Simulated exposure effect | Average simulated proportion |  | Scenario II<br>Confounding | Scenario III<br>Undirected pleiotropy | Scenario IV<br>Directed pleiotropy | Scenario V<br>Dependence |
| --- | --- | --- | --- | --- | --- | --- |
| Distinct | 0.373 | MV-MR | 0.742 (0.437) | 0.516 (0.500) | 0.516 (0.500) | 0.680 (0.467) |
|  |  | MR-BMA | 0.779 (0.415) | 0.534 (0.499) | 0.567 (0.496) | 0.697 (0.460) |
|  |  | mSSL | 0.763 (0.426) | 0.693 (0.461) | 0.690 (0.463) | 0.697 (0.460) |
|  |  | MR<sup>2</sup> | **0.783** (0.413) | **0.697** (0.460) | **0.722** (0.448) | **0.743** (0.437) |
| Shared = 2 | 0.315 | MV-MR | **0.566** (0.496) | 0.224 (0.417) | 0.269 (0.444) | 0.438 (0.496) |
|  |  | MR-BMA | 0.551 (0.498) | 0.260 (0.439) | 0.309 (0.462) | 0.433 (0.496) |
|  |  | mSSL | 0.551 (0.498) | 0.422 (0.494) | 0.430 (0.495) | 0.453 (0.498) |
|  |  | MR<sup>2</sup> | 0.556 (0.496) | **0.435** (0.496) | **0.453** (0.498) | **0.463** (0.496) |
| Shared = 3 | 0.122 | MV-MR | **0.443** (0.497) | 0.173 (0.379) | 0.173 (0.379) | 0.265 (0.442) |
|  |  | MR-BMA | 0.381 (0.486) | 0.212 (0.409) | 0.240 (0.428) | 0.286 (0.452) |
|  |  | mSSL | 0.412 (0.493) | 0.298 (0.458) | 0.337 (0.473) | **0.306** (0.461) |
|  |  | MR<sup>2</sup> | 0.392 (0.489) | **0.308** (0.462) | **0.346** (0.476) | **0.306** (0.461) |
| Shared = 4 | 0.023 | MV-MR | **0.440** (0.498) | 0.133 (0.342) | 0.133 (0.342) | 0.231 (0.423) |
|  |  | MR-BMA | 0.400 (0.492) | 0.200 (0.403) | 0.133 (0.342) | 0.269 (0.445) |
|  |  | mSSL | **0.440** (0.498) | **0.267** (0.445) | **0.267** (0.445) | 0.346 (0.478) |
|  |  | MR<sup>2</sup> | **0.440** (0.498) | 0.233 (0.430) | **0.267** (0.445) | **0.385** (0.488) |

### Cardiometabolic risk factors for cardiovascular diseases

For the first real application example, as cardiovascular disease outcomes (CVDs), we consider atrial fibrillation (AF), cardioembolic stroke (CES), coronary artery disease (CAD), heart failure (HF) and peripheral artery disease (PAD). Common polygenic exposures were selected according to the National Health Service guidelines on causes for CVD Web page (https://www.nhs.uk/conditions/coronary-heart-disease/causes/ - last reviewed on the 10th March 2020). For high cholesterol, we include five major lipoprotein-related traits (apolipoprotein A1 (ApoA), apolipoprotein B (ApoB), highdensity lipoprotein (HDL), low-density lipoprotein (LDL) and triglycerides (TG). Obesity is measured by body mass index (BMI), exercising regularly (moderate-to-vigorous intensity exercises during leisure time) by physical activity (PA) defined by moderate-to-vigorous intensity exercises during leisure time and high blood pressure by systolic blood pressure (SBP). We also include a lifetime smoking index (SMOKING), a composite of smoking initiation, heaviness, duration and cessation, and type 2 diabetes (T2D) as exposures. Given the strong epidemiological, genetical and causal relationships between these exposures a multivariable MR design is necessary to account for potential horizontal pleiotropy and to facilitate selection of likely causal exposures. Genetic associations with exposures and outcomes are derived from publicly available summary-level data, see Supplemental Table S.12 for an overview of the data sources.

IVW is performed before the analysis for all outcomes and exposures using weights derived jointly from all responses as described in Appendix. Moreover, the summary-level genetic associations with the exposures are standardised before the analysis. This allows us to interpret and compare the estimated exposures effect size for each outcome and, more importantly, across outcomes. For a complete description of the pre-processing and instrumental variable selection steps, we refer to Supplemental Information. Briefly, we select *n* = 1, 540 independent genetic variants associated with any of the ten exposures as IVs after clumping. Results are obtained after removing outliers or high-leverage and influential observations using scaled CPO, and fitting the proposed model on the remaining *n* = 1, 533 IVs, see Supplemental Figure S.6.

MR^2^ identifies several exposures shared among CVDs as highlighted in Figure 5A and B which show the marginal posterior probability of inclusion (mPPI) and the posterior mean direct effect sizes (95% credible interval) for each exposure-outcome combination, respectively. Significant mPPIs, and corresponding direct effect estimates, are selected controlling False Discovery Rate (FDR) at 5% (see Appendix) which corresponds to mPPIs ≥ 0.77 (Supplemental Figure S.7A). For clarity of presentation, mPPIs and direct effect sizes for non-selected outcome-exposure pairs are plotted as zero.

**Figure 5.**
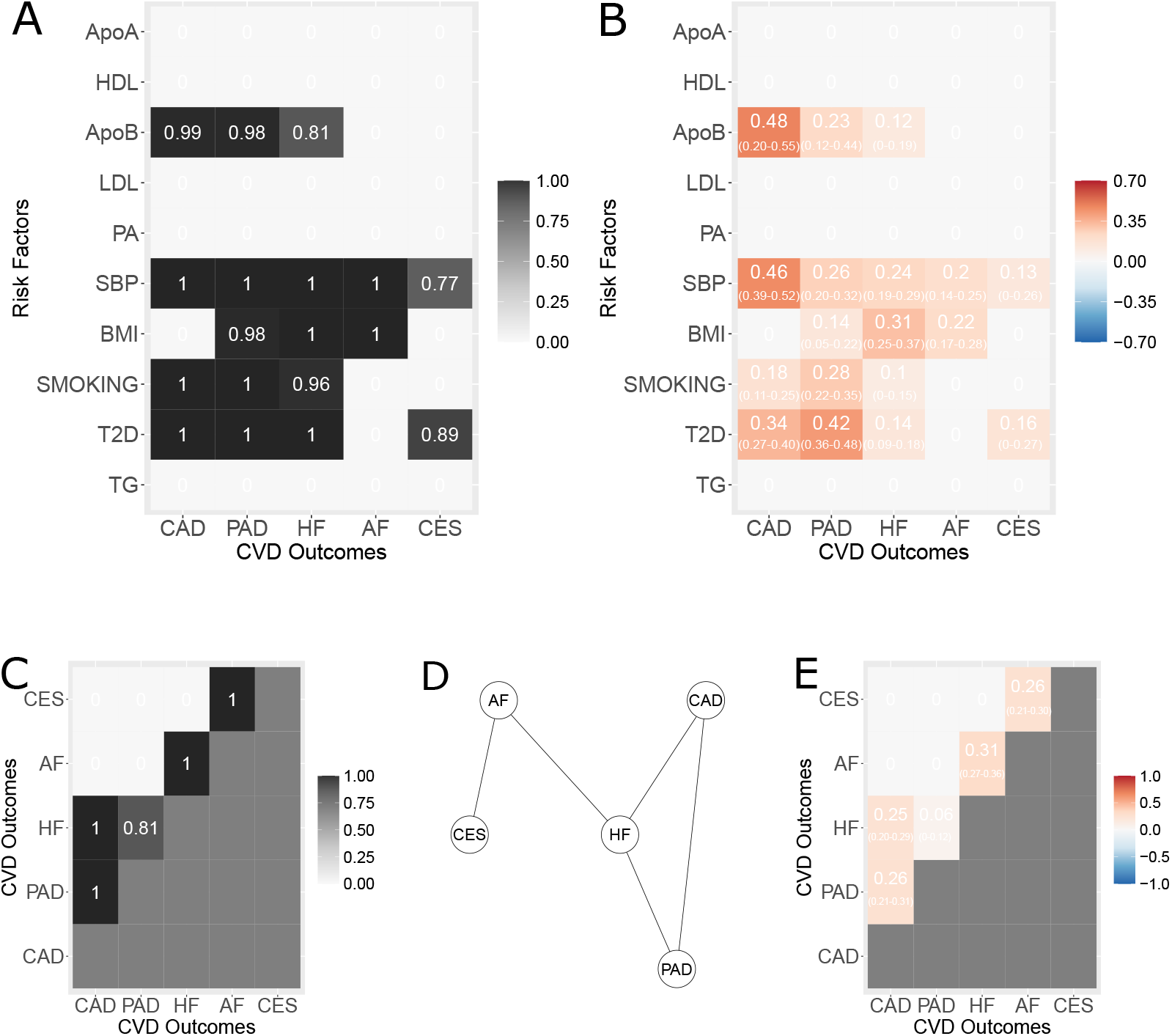
Results of the multivariable multi-response MR^2^ model in application example 1 on common exposures for cardiovascular disease outcomes (CVDs). (**A**) Marginal posterior probability of inclusion (mPPI) of each exposure (*y*axis) against each outcome (*x*-axis). Selected mPPIs for each exposure indicate whether an exposure is shared or distinct among multiple CVDs. (**B**) Posterior mean (95% credible interval) of the direct causal effect of each exposure (*y*-axis) against each outcome (*x*axis). For clarity of presentation, mPPIs and estimated direct effect sizes for non-selected outcome-exposure pairs (mPPI < 0.77 at 5% FDR) are plotted as zero. (**C**) Edge posterior probability of inclusion (ePPI) among outcomes. Only the upper triangular matrix is depicted. (**D**) Undirected graph estimated by using the selected ePPIs showing the residual dependence between outcomes not explained by the exposures. (**E**) Posterior mean (95% credible interval) of the partial correlations between outcomes. For clarity of presentation, ePPIs and partial correlations for non-selected outcomes pairs (ePPI < 0.78 at 5% FDR) are plotted as zero. Only the upper triangular matrix is depicted.

Results provided by MR^2^ can be read in two different ways. A traditional “vertical” way where, for each outcome, the significant exposures are highlighted. For instance, CAD has four significant exposures, *i.e*., ApoB, SBP, T2D and SMOKING in order of importance by looking at the posterior mean direct effect estimates. PAD and HF have the same ones, although in a different order. On the other hand, genetically-predicted levels of SBP and BMI are associated with AF and genetically-predicted levels of SBP and risk of T2D are associated with CES, respectively.

The main novelty of the proposed methodology is that it allows reading, interpreting and comparing the results also “horizontally” across outcomes. For instance, ApoB shows the strongest risk-increasing effect on CAD among all exposure-outcome combinations with a posterior mean direct causal effect of 0.48. Moreover, it is also selected for PAD with halved effect estimate (0.23) and around four times lower risk increase for HF (0.11). Similarly, the joint posterior probability of inclusion (jPPI) can be interpreted as the relevance of an exposure for a group of outcomes. For instance, ApoB has a strong jPPI of 0.78 to be jointly causal for CAD, PAD and HF (Supplemental Figure S.7C). Taken together, these findings extend the results of a previous study which found ApoB as a shared exposure for CAD and PAD [17] and for the first time implicates a likely causal role of ApoB also for HF.

MR^2^ provides also a more rigorous statistical control of the null hypothesis (no causal effects) since it takes into account both the conditional independence among exposures (multivariable) and across responses (multitrait). For example, conditional on ApoB, there is no evidence for any other major lipoprotein-related trait to have a likely causal role for any outcome and conditional on CAD, PAD and HF, there is no evidence for ApoB on any other outcome.

A second example regarding the ability of MR^2^ to disentangle complex causal relationships and advance cardiovascular domain knowledge is related to SBP. SBP is shown to increase the risk for all five CVDs considered, with a substantial jPPI of 0.77 for all five outcomes. However, SBP has the strongest posterior mean direct effect on CAD (0.46) and then it decreases steadily across the other responses, PAD (0.26), HF (0.24), AF (0.2) and CES (0.13), suggesting that SBP-lowering may not be a broad therapeutic target (as already shown in RCTs) or it would require potent SBP-lowering to meaningfully impact (in increasing order) HF, AF and CES at a population-level. Similar considerations can be extended to other exposures selected.

Almost all significant exposures have a large causal effect on CAD and PAD, but much lower on HF, AF and CES except BMI on HF and AF. Therefore, it seems plausible that these traditional cardiovascular exposures are not able to fully describe the disease aetiology of HF, and in particular, AF and CES. In turn, these results suggest that there may be other exposures not considered here as the main causes of these diseases. In contrast, other exposures included in this analysis such as PA that, although considered an important exposure in medical practice (NHS guidelines), conditionally on all the others, are not significant for any CVD. This suggests that beneficial effects of physical activity on these outcomes is likely mediated by the other traditional cardiometabolic risk factors.

MR^2^ also acknowledges residual correlation among summary-level responses which is not accounted for by the selected exposures as shown in Figures 5C-E, which depict the edge posterior probability of inclusion (ePPI), the indirect graph estimated by using the selected ePPIs and the posterior mean (95% credible interval) of the partial correlations between outcomes, respectively. Significant ePPIs and corresponding partial correlation are selected controlling False Discovery Rate (FDR) at 5% which corresponds to ePPIs ≥ 0.78 (see Supplemental Figure S.7B).

Significant residual dependence between the outcomes not explained by the exposures is identified between CAD and HF and between CAD and PAD with summary-level residual partial correlation 0.26 and 0.25, respectively, reflecting known vertical and horizontal pleiotropy of CAD being a likely cause of HF [19, 20] and horizontal pleiotropy between CAD and PAD. In contrast, PAD and HF show significant but four times lower level residual partial correlation (0.06). Other important residual partial correlations highlight the disease pathway HF-AF-CES as illustrated in Figure 5C. Compared to the empirical partial correlations between summary-level genetic associations with the responses without conditioning on the exposures (Supplemental Figure S.5B), the genetically predicted levels of exposures are able to explain around 32% and 21% of the summary-level partial correlation between CAD and PAD and between CAD and HF, respectively, and almost 62% of the residual correlation between PAD and HF. However, not all exposures contribute in the same way to this remarkable decrease. ApoB seems not as important (jPPI = 0.8) as the other associated exposures (jPPI > 0.96) to explain the dependence between PAD and HF, see Supplemental Figure S.7C. Finally, a little reduction is observed for the disease pathway HF-AF-CES, supporting the earlier hypothesis that other important shared exposures may be missing from the proposed MR model. However, as mentioned earlier and shown in eq. (8), we cannot rule out that, besides shared pleiotropy, some non-genetic factors may be responsible for the observed residual correlation.

We conclude this section comparing the results obtained by MR^2^ with existing multivariable MR methods (see Supplemental Table S.1), including multivariable MR-Egger [29] to confirm that, when dealing with multiple outcomes, there are different assumptions regarding the effect of the unmeasured pleiotropy and multivariable MR-Egger may not able to detect it.

MV-MR is not able to identify any lipid exposure for any outcome considered except for TG for AF at 5% Benjamini-Hochberg FDR (Supplemental Table S.13). Multivariable MR is not designed for the analysis of many exposures which are highly correlated [15]. Indeed, multi-collinearity reduces the precision of the estimated direct causal effects, which weakens the statistical power of the MV-MR model. Due to the strong correlation between genetic associations of lipid exposures (Supplemental Figure S.5C), MV-MR misses ApoB as likely causal exposure for CAD, PAD and HF. Similar results in terms of exposures selected and effect sizes are seen when multivariable MR-Egger is used, with a significant unmeasured horizontal pleiotropy identified only in CES (Supplemental Table S.14). Note that neither MV-MR nor multivariable MR-Egger can provide a clear picture of the effect size of SBP across disease outcomes, with much larger effect estimates and a narrow difference between the largest (CAD) and the smallest (CES).

MR-BMA is not able to draw a clear distinction between the exposures selected and excluded as shown in Supplemental Table S.15. This is exemplified in CAD where the exposures included depend on the multiple testing correction applied. When using a strict Bonferroni threshold, MR-BMA identifies in decreasing order SBP, SMOKING, T2D and ApoB with 0.66 as the smallest mPPI, while with a more lenient FDR threshold also HDL, LDL and BMI are additionally included with the smallest mPPI of 0.29. Regarding the other outcomes, the order of importance of the exposures is different from MR^2^ with remarkably larger model-averaged causal effect estimates (MACEs) for PAD and HF. MR-BMA suffers the same problems as MV-MR and multivariable MR-Egger regarding the magnitude of the direct causal effect estimates of SBP on the outcomes, demonstrating that this problem affects all single-trait methods regardless of their implementation. Interestingly, MR-BMA does not identify T2D as an exposure for CES, which is instead detected by MR^2^, MV-MR and multivariable MR-Egger, favouring in contrast BMI.

### Lipidomic risk factors for cardiovascular diseases

The first application example prioritizes ApoB as a shared exposure for three out of five CVD. Moreover, conditional on ApoB, no other major lipoprotein-related trait has a likely causal role for any outcome. Our next step is to better understand molecular determinants of ApoB by considering ten ApoB-containing lipoprotein subfractions of different sizes, ranging from small-large to extra-extra large very-large density lipoproteins, measured using nuclear magnetic resonance (NMR) spectroscopy [54]. Identification of specific sub-fractions to different CVD manifestations can help our understanding of the pathophysiology of the disease and provide insights into pathophysiology, molecular mechanisms, risk stratification and treatment [17]. However, performing MR using metabolites as exposures is a difficult task given the strong correlation and intricate dependence that exist between them, see Supplementary Figures S.8C and B.

For a description of the pre-processing and instrument selection steps, we refer to Supplemental Information. Briefly, given the prior hypothesis of ApoB as the leading exposure for CVDs, we select genetic variants associated with ApoB in UK Biobank at genome-wide significance, resulting in *n* = 148 IVs after clumping. Results are obtained after removing outliers or high-leverage and influential observations using scaled CPO, and fitting the proposed model on *n* = 141 IVs, see Supplemental Figure S.9.

When lipidomic risk factors are used for CVDs, results obtained by MR^2^ are very sparse with few significant direct causal effects at 5% FDR (mPPI < 0.29). Moreover, the separation between significant and non-significant causal associations is difficult, see Supplemental Figure S.10A. The proposed model identifies XS.VLDL.P as a shared exposure for both PAD and HF with mPPIs 0.32 and 0.31, respectively, and a jPPI of 0.14 (Supplemental Figure S.10C) with direct causal effects of 0.15 and 0.13 (Figure 6A and B). Distinct exposures are detected for CAD, where smaller particle sizes, IDL.P and L.LDL.P, are prioritized with mPPI 0.41 and 0.59 and direct causal effects of 0.4 and 0.63, respectively. No subfractions are identified for the other disease outcomes.

**Figure 6.**
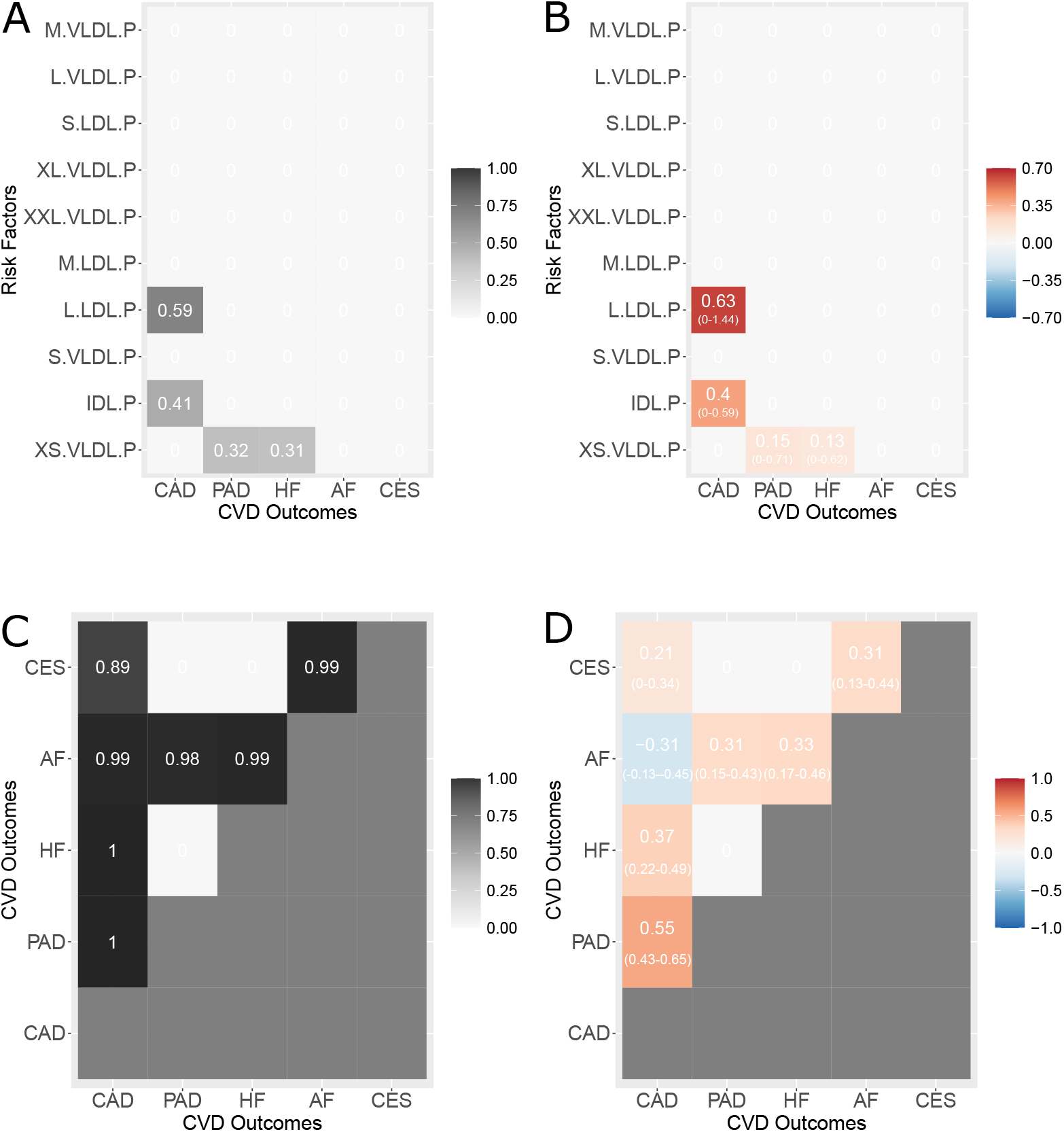
Results of the multi-response MR (MR^2^) model in application example 2 on molecular exposures for cardiovascular disease outcomes (CVDs). (**A**) Marginal posterior probability of inclusion (mPPI) of each exposure (*y*-axis) against each outcome (*x*-axis). Selected mPPIs for each exposure indicate whether an exposure is shared or distinct among multiple CVDs. (**B**) Posterior mean (95% credible interval) of the direct causal effect of each exposure (*y*-axis) against each outcome (*x*-axis). For clarity of presentation, mPPIs and estimated effect sizes for non-selected outcome-exposure pairs (mPPI < 0.29 at 5% FDR) are plotted as zero. (**C**) Edge posterior probability of inclusion (ePPI) among outcomes. Only the upper triangular matrix is depicted. (**D**) Posterior mean (95% credible interval) of the partial correlations between outcomes. For clarity of presentation, ePPIs and partial correlations for non-selected outcomes pairs (ePPI < 0.85 at 5% FDR) are plotted as zero. Only the upper triangular matrix is depicted.

In contrast to the previous example, when dealing with molecular exposures, credible intervals (CIs) are large, confirming the complexity of the analysis. For the strongest signal, L.LDL.P-CAD combination, the 95% CI ranges between 0 and 1.41. This is due to a combined effect of multicollinearity (although the designed independent prior in eq. (13) protects against it, see Appendix) and the small number of genetic variants associated with ApoB in UK Biobank. Thus, while L.LDL.P (or IDL.P) could be a likely cause for CAD, its large CI suggests caution.

As expected, there is a substantial residual partial correlation in almost all combinations of summary-level outcomes as depicted in Figure 6C and D, highlighting the existence of non-lipoprotein pleiotropic pathways between these traits not intercepted by the selected exposures (SBP, BMI, SMOKING, T2D pathways are missing by design) and, possibly, the effects of non-genetic factors on the responses.

When high levels of residual correlation are present and cannot be “explained away” or be “accounted for” by the exposures, the advantage of an MR multi-response model to reduce false positives is evident. For instance, MR-BMA (Supplemental Table S.18) identifies genetically-predicted levels of L.VLDL.P to be associated also with AF, possibly because AF and CAD are correlated at the summary level (Supplemental Figure S.8). Similarly, the causal effect of S.VLDL.P on AF is likely a false positive by looking at its small MACE. MV-MR is not able to identify any exposure for any outcome which is significant after multiple testing correction (Supplemental Table S.16) even with a less conservative Benjamini-Hochberg FDR procedure, demonstrating that standard methodology is not adapted to handle highly correlated exposure data. Similar conclusions can be extended to multivariable MR-Egger (Supplemental Table S.17) with no significant intercept identified for any outcome.

## Discussion

Here, we present MR^2^, a novel MR framework to analyze multiple related outcomes in a joint model and to define shared and distinct causes of related health outcomes. Based on the SUR model, information across outcomes is shared in MR^2^ by estimating the residual correlation of multivariable MR models for each outcome. MR^2^ is formulated on the summary level where the genetic variants used as IVs are considered observations. Residual correlation in the proposed model is consequently interpreted as the correlation between summary-level outcomes measured on the IVs not accounted for by the genetic associations with the exposures. We show, both theoretically and in a simulation example, how unmeasured shared pleiotropy induces residual correlation between summary-level genetic associations of the outcomes.

While residual diagnostics is an important strategy in summary-level univarible MR models to detect horizontal pleiotropy [55, 51] affecting one outcome, only multi-response MR models, like MR^2^, can detect how much cross-variation between outcomes is unexplained after accounting for the exposures of interest. We show in an extensive simulation study that MR^2^ has more power to detect true causal exposures, yields a better separation between causal and non-causal exposures and improves the accuracy of the effect estimation over existing multivariable MR methods that consider only one outcome at-a-time, such as MV-MR and MR-BMA. Moreover, MR^2^ demonstrates more power when an exposure is causal for more than one outcome.

Thanks to the formulation as a joint multi-response model MR^2^ can distinguish between shared and distinct exposures for the disease which is essential to define interventions that reduce the risk of more than one disease. We illustrate this in our application examples considering five CVDs. Multi-response models like MR^2^ are a necessary contribution to understanding better the causes of multi-morbidity. In particular, the discovery of shared and distinct causes of diseases may help define interventions with co-benefits, *i.e*., interventions which reduce the risk of more than one outcome. For instance, in our first application example, we have identified ApoB as likely causal exposure for CAD, PAD and HF even when accounting for other lipoprotein measures including LDL cholesterol.

While ApoB is an established exposure for CAD and PAD based on genetic evidence [56, 37, 17], we demonstrate that there is an independent effect of ApoB also on HF. This has the following two implications: for one, existing lipid-lowering therapies should be evaluated in terms of their impact on reducing ApoB and, second, future lipid-lowering therapies may be better tailored to reduce ApoB concentration. Moreover, this highlights the importance of including various disease endpoints in RCTs since the intervention may have benefits not just for the main disease of interest but also for other related diseases.

In our application examples, we discover residual correlation between CAD and PAD and between CAD and HF, as well as between the disease pathway HF-AF-CES, which was not accounted for by either common cardiometabolic disease exposures, including lipid traits, blood pressure and obesity, or by lipid characteristics measured using NMR spectroscopy. An important contributor to the residual correlation, when considering common cardiovascular disease exposures, are molecular pathways which are not accounted for when considering traits like ApoB or obesity. These pathways, such as inflammation or stress response, are highly polygenic with hundreds of independent regions in the genome associated with them. Another source of residual correlation between outcomes is the consequence of non-genetic factors that act exclusively on the responses. A final possible source of the observed residual correlation can be mediation, where genetic predisposition for one outcome may cause another, as may be the case for CAD and HF, as suggested for example by the Framingham Heart study [20].

There are also limitations to our work. First of all, weak instrument bias is towards the null in univariable MR [57], but can go towards any direction for multivariable MR depending on the correlation between exposures [15]. A necessary future extension of the approach is to make MR^2^ more robust concerning weak instruments [58]. Second, special care needs to be taken when selecting genetic variants as IVs. Importantly, the interpretation of any MR model is conditional on the IVs selected. For example, in our second data example, we have selected genetic variants based on their association with ApoB, which we identified in our primary analysis. Therefore, results need to be interpreted in light of this choice and may differ when re-selecting for example based on LDL-cholesterol. In practice, we recommend following the guidelines for reporting MR studies [59, 60] for further details.

The development of multi-response MR models on the individual level is another important future direction given the availability of large-scale biobanks with sufficient follow-up time allowing for the development of multiple disease outcomes. Such an analysis would let us study the presence of more than one disease in the same individual instead of analysing the shared genetic basis of the outcomes as in our current work.

While our study has been motivated to detect common causal exposures for multi-morbid health conditions, MR^2^ can be applied to any type of related outcomes. Potential application examples may include molecular biomarkers as outcomes, for example, a recent study has investigated the effect of sleep deprivation on the metabolome [61] or of morning cortisol levels on inflammatory cytokines [62]. Similarly, MR^2^ can be used to define exposures for heritable imaging phenotypes measured on the brain [63], heart [64] or body composition [65].

In conclusion, we present here MR^2^, the first summary-level MR method that can model multiple outcomes jointly and account for residual correlation between the outcomes. Moreover, MR^2^ can distinguish between shared or distinct causes of diseases, enhancing our understanding regarding which interventions can target more than one disease outcome.

## Supplemental information

Supplemental information includes details of the derivation of the residual correlation between summary-level outcomes, the MCMC implementation of the proposed model and Supplemental Tables and Figures to support the results of the simulation study and the two application examples.

## Web resources

The multivariable, multi-response Mendelian randomization R package MR2 is freely available on https://github.com/lb664/MR2/. It includes examples that explain how to generate the simulated data and run the algorithm. Post-processing routines are also included.

## Acknowledgements

The authors are thankful to Zhi Zhao and Manuela Zucknick for their insightful comments on an early draft of the manuscript.

The authors gratefully acknowledge the United Kingdom Research and Innovation Medical Research Council grants MR/W029790/1 (V.Z., L.B.) and MC UU 00002/7 (S.B.), The Alan Turing Institute under the Engineering and Physical Sciences Research Council grant EP/N510129/1 (L.B.), Marmaduke Sheild Fund (L.B.), the British Heart Foundation Centre of Research Excellence at Imperial College London grant RE/18/4/34215 (D.G.), Wellcome Trust award 225790/Z/22/Z (S.B.), the Institute for Translational Medicine and Therapeutics of the Perelman School of Medicine at the University of Pennsylvania (M.G.L.), the NIH/NHLBI National Research Service Award postdoctoral fellowship (T32HL007843) (M.G.L.), the Measey Foundation (M.G.L.) and US Department of Veterans Affairs award IK2-CX001780 (S.M.D.). This publication does not represent the views of the Department of Veterans Affairs or the United States Government.

## Author contributions

Conceptualization: V.Z., L.B., D.G.; Methodology: V.Z., L.B.; Formal Analysis: L.B., V.Z.; Resources: (all collaborators); Data curation: V.Z., D.G.; Writing – original draft: V.Z., L.B.; Writing – review & editing: L.B., V.Z., all collaborators; Visualization: L.B., V.Z.; Supervision: L.B.

## Appendix

### Inverse variance weighting for multiple responses

When considering one outcome at-a-time, the univariable MR model in eq. (1) and multivariable MR model in eq. (2) weight a genetic variant *i* = 1,…, *n*, used as IV, by the standard error of the genetic association with the outcome 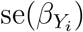. These weights are also known as the “first-order weights” [66].

In order to generalise the IVW for multiple outcomes, we note that the standard error of the genetic association with a binary trait *Y* of a genetic variant *i*, with minor allele frequency MAF_*i*_, is [67]

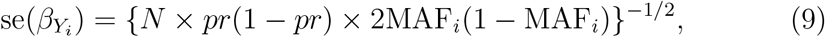

where *pr* is the percentage of cases and *N* is the total sample size. In eq. (9), the only quantity that depends on the genetic variant *i* is the minor allele frequency MAF*_i_* which may be considered as comparable when taking summary-level data from cohorts with similar ethnicity. All other parameters are specific for trait *Y* and do not depend on the genetic variant *i* and, consequently, are not relevant when weighting individually the genetic variants.

When considering *q* outcomes, we propose to use as weight *ω_i_*, *i* = 1,…, *n*, the mean of the *q* standard errors of the genetic associations with each outcome, *i.e*., 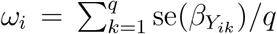, which essentially, under the hypothesis of cohorts with similar ethnicity, will result in weighting by the average of the inverse of the genetic variant standard deviations defined by {2MAF*_i_*(1 − MAF*_i_*)}^1/2^.

### Bayesian multi-response MR

An alternative way to the SUR model in eq. (3) to model jointly the *q* related outcomes is to describe their dependence through a copula function. A *q*-variate function *C*(*u*_1_*,…, u_q_*), where *C* : [0, 1]*^q^* → [0, 1], is called a copula if it is a continuous distribution function and each marginal is a uniform distribution function on [0, 1]. If *F_Y_*1 (·)*,…, F_Y_q* (·) are the marginal cumulative density functions (cdfs) of the random variables *Y*_1_*,…, Y_q_*, their joint cdf can be described through a specific copula function *C* as

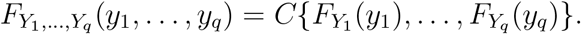

The Gaussian copula *C* is described through the function

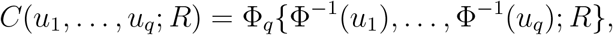

where Φ*_q_*(; *R*) is the cdf of an *q*-variate Gaussian distribution with zero mean vector and correlation matrix *R* and Φ^−1^(·) is the inverse of the univariate standard Gaussian cdf. If all the marginal distributions are continuous, the matrix *R* can be interpreted as the correlation matrix of the elements of *Y* and zeros in its inverse imply the conditional independence between the corresponding elements of the responses.

The Gaussian copula regression (GCR) model is described by the transformation

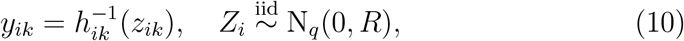

where *z_i_*_1_*,…, z_iq_* are realizations from *Z_i_* and 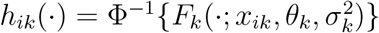 for each *i* = 1,…, *n* and *k* = 1*,…, q*, *x_ik_* is the *p*-dimensional vector of predictors for the *i*th sample and the *k*th response and *θ_k_* = (*θ_k_*_1_*,…, θ_kp_*)*^T^* is a *p*-dimensional vector of regression coefficients.

A detailed discussion of the Bayesian formulation of the GCR model in eq. (10) with multiple responses of any type is presented in [40]. In the following, we summarise the main aspects of the proposed multi-response multivariable MR model when all margins are univariate Gaussian. Details of the MCMC algorithm are presented in Supplemental Information.

The SUR model is a special case of eq. (10) when all margins are univariate Gaussian with mean *x_ik_θ_k_* and variance 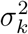 and *R* is the correlation matrix. Thus, eq. (3) and (4) can be written in terms of a Gaussian copula regression model as

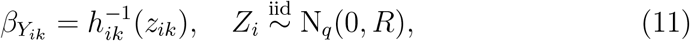

with or, equivalently,

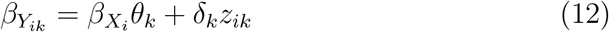

for each *i* = 1,…, *n* and *k* = 1*,…, q* and with *R* as in eq. (4).

### Priors specification

The selection of important exposures is achieved by utilizing a binary latent vector *γ_k_* = (*γ_k_*_1_*,…, γ_kp_*), *k* = 1*,…, q*, where *γ_kj_* is 1 if, for the *j*th exposure and the *k*th response, the effect estimate is different from zero and 0 otherwise. Specifically, we assume that for each *k*, the effects in eq. (12) are distributed as

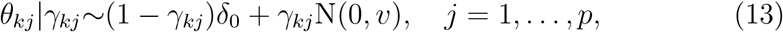

where *δ*_0_ denotes a point mass at zero and *v* is a fixed value. Sparsity for the selection of important exposures for the *k*th response is enforced by specifying the hierarchical structure

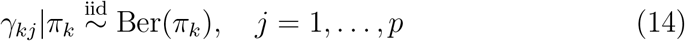

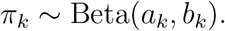

Integrating out *π_k_* in eq. (14), it is readily shown that marginally

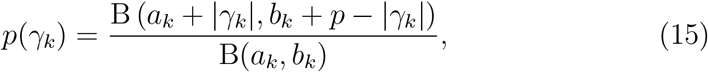

where B(·, ·) is the beta function and 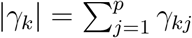. The hyperparameters *a_k_* and *b_k_* can be chosen using prior information about the average number of important exposures associated with the *k*th response and its variance.

Since the Gaussian copula regression model in eq. (11) is defined through the correlation matrix *R*, we need first to expand *R* into a covariance matrix. We define the transformation *W* = *Z*Δ, where *Z* is the (*n × q*)-dimensional matrix of the Gaussian latent variables obtained by inverting eq. (12)

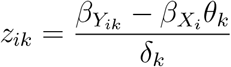

and Δ is an *q* × *q* diagonal matrix with elements *δ_k_*, *k* = 1*,…, q*. Then,

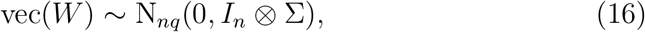

where Σ = Δ*R*Δ and *I_n_* is a diagonal matrix of dimension *n* that encodes the independence assumption among the genetic variants *G* (achieved by pruning). In this framework, the correlation matrix can be obtained by the inverse transformation *R* = Δ^−1^ΣΔ^−1^. We utilize the theory of decomposable graphical models [42] to perform a conjugate analysis of the covariance structure of the model since the hyper-inverse Wishart distribution is a conjugate prior distribution for the covariance matrix Σ with respect to the symmetric adjacency matrix G of a decomposable graph 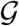.

Finally, we assign the following prior structure on the symmetric adjacency matrix G and Δ. Let *g_ℓ_*, *ℓ* = 1*,…, q*(*q* 1)/2, be the binary indicator for the presence of the *ℓ*th off-diagonal edge in the lower triangular part of the symmetric adjacency matrix G of the decomposable graph 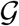. To select important elements of the inverse covariance matrix, we assume the sparsity prior

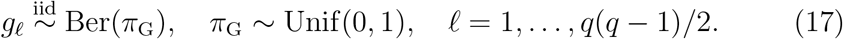

We denote by *p*(G) the induced marginal prior on the symmetric adjacency matrix and define the joint distribution of Δ, *R* and G as *p*(Δ, *R*, G) = *p*(Δ|*R*)*p*(*R*|G)*p*(G), where

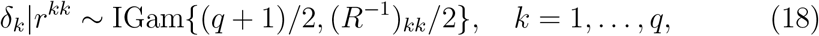

with (*R*^−1^)*_kk_* the *k*th diagonal element of *R*^−1^ [68].

### Outliers, high-leverage and influential observations

Detection of outliers is performed by using the conditional predictive ordinate (CPO) [52] to detect genetic variants *G_i_*, *i* = 1,…, *n*, as outliers in the multiresponse MR model. The CPO is also known as the leave-one-out crossvalidation predictive density [69] which, for the multi-response MR model in eq. (11), is defined as

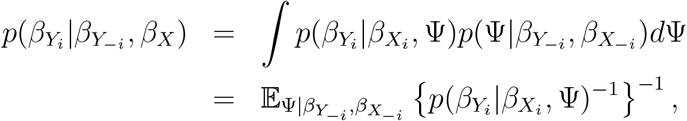

where 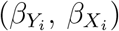 and 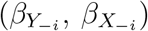 are the IVW-standardised summary-level associations with the genetic variant *G_i_* and all the remaining genetic variants, respectively, Ψ is the whole parameter space Ψ = Γ, Θ, G, Δ, *R* with Γ = (*γ*_1_*,…, γ_q_*)*^T^* and Θ = (*θ*_1_*,…, θ_q_*)*^T^*.

The CPO describes the posterior probability of observing the *q*-dimensional vector of values of *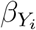* when the model is fitted to all data except *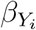*, with a larger value implying a better fit of the model for *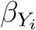* and a very low CPO value suggests that the summary-level associations with the genetic variant *G_i_* are either an outlier or a high-leverage and influential observation.

The CPO can be calculated by using the output of the MCMC algorithm. Let *T* be the number of recorded MCMC iterations. By considering the inverse likelihood across *T* iterations, the estimated CPO for each genetic variant *G_i_*, *i* = 1,…, *n*, is

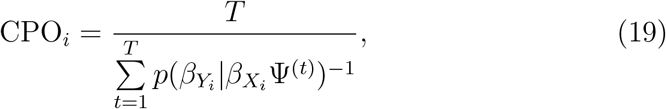

where Ψ^(*t*)^ is *t*th posterior sample of Ψ obtained from the MCMC algorithm. Thus, the Monte Carlo estimate of the CPO is obtained without actually omitting 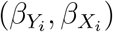 from the estimation of the posterior distribution of Ψ and is provided by the harmonic mean of the likelihood [70].

Regarding the threshold for the detection of outliers or high-leverage and influential observations, log-inverse-CPOs larger than 40 can be considered as possible outliers and higher than 70 as extreme values [71]. [72] recommends scaling CPOs by dividing each one by its individual maximum (likelihood) recorded across the MCMC iterations and considering observations with scaled CPOs under 0.01 to be outliers or high-leverage and influential observations. If few CPOs are less than 0.01, the model is considered to fit adequately.

### False Discovery Rate

The marginal posterior probability of inclusion (mPPI) which measures the strength of direct causal association between each exposure-response combination and the edge posterior probability of inclusion (ePPI) which describes the strength of the residual conditional correlation between each responses’ pair are utilised to select significant exposure-response combinations and important pairs of dependent responses.

Let

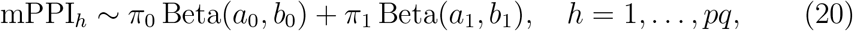

be the two-component mixture model that classifies the mPPI for each exposure-response combination into the null (*H*_0_) or the alternative distribution (*H*_1_) parameterized as beta densities with parameters (*a*_0_, *b*_0_) and (*a*_1_, *b*_1_), respectively, with weights *π*_0_, *π*_1_ ≥ 0 and *π*_0_ + *π*_1_ = 1.

In the real application examples, the parameters of the mixture model in eq. (20) are estimated using the EM algorithm [73]. The posterior probability of allocation to the alternative hypothesis of each mPPI is

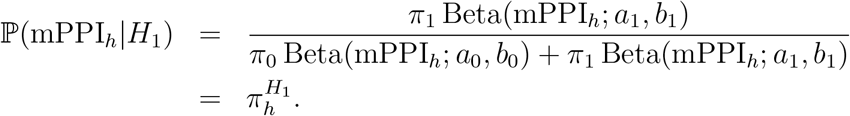

Finally, let 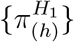, *h* = 1*,…, pq*, the sequence in decreasing order of the posterior probabilities of allocation to the alternative hypothesis of the mPPIs for each exposure-response combination. Significant mPPIs are chosen such that

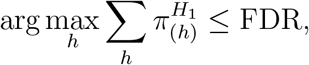

where FDR is the designed level.

### Simulation study

Our simulation study is formulated in a two-sample summary-level MR design, where *N* = 100, 000 independent individuals are simulated, of which half are used to compute the genetic associations with the exposures and half to compute the genetic associations with the outcomes. In the following, we indicate with the subscripts “*X*” and “*Y*” the relevant quantities that are associated with the exposures and the responses, respectively. For instance, *N_X_* and *N_Y_* indicate the sample size used in the genetic associations with the exposures and the outcomes, respectively.

In all simulation scenarios, we consider *p* = 15 exposures, *q* = 5 outcomes and *n* = 100 independent genetic variants as IVs. Genotypes for the *i*th genetic variant and each individual *l* are simulated independently according to a binomial distribution with minor allele frequency (MAF) equal to 0.05, *i.e*., 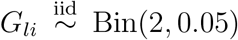, *l* = 1,…, *n* and *i* = 1,…, *n*. Without loss of generality, the resulting matrix of genotypes *G* is split into two equally sized groups, *G_X_* and *G_Y_*, of dimension *N_X_ × n* and *N_Y_ × n* with *N_X_* = *N_Y_* = *N*/2, respectively. All genetic variants are considered with equal weights, thus no IVW is needed given that the same MAF at 5% is used to simulate the genotypes.

Overall, the data generation process consists of two steps. In the first step, the raw data for the exposures *X* and the responses *Y* are simulated. Then, second step, two-sample summary-level data are obtained as the regression coefficients 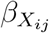, *i* = 1,…, *n*, *j* = 1*,…, p*, from a univariable regression in which the exposure *X_j_* is regressed on the genetic variant *G_i_* in sample one and the regression coefficients 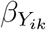, *i* = 1,…, *n*, *k* = 1*,…, q* from a univariable regression in which the outcome *Y_k_* is regressed on the genetic variant *G_i_* in sample two. In the following, we detail each step and how we simulate the quantities involved.

- For the first stage of the simulation study, the *j*th exposure, *j* = 1*,…, p*, is generated by

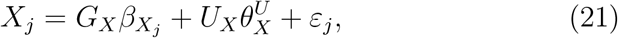

where *G_X_* and *U_X_* are the matrix of genotypes of the *n* IVs and the confounder *U* measured on the same *N_X_* = 50, 000 individuals, respectively, and 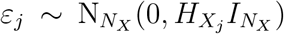, where *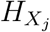* is the *j*th diagonal element of the (*p* × *p*)-dimensional matrix 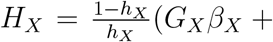 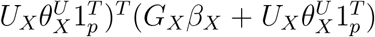 with 1*_p_* a *p*-dimensional vector of ones, 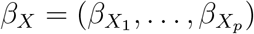 and *h_X_* the desired level of the proportion of variance explained, fixed at 10% in all simulated scenarios. The confounder *U* is drawn from a multivariate standard Gaussian distribution, *i.e*., *U* ~ N*_N_* (0, *I_N_*) and, then, split into two equally sized vectors *U_X_* and *U_Y_* with effect 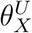 impacting all the exposures *X* and 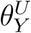 effecting all the outcomes *Y*. Their value is fixed at 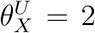 and 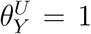 in main Results. Instrument strength depends on the degree of confounding, where the average *F* -statistic is on average around 45 and 35 for the weakest degrees of confounding 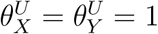 and for the strongest confounding effects 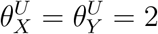, respectively. The effects 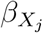 of the genetic variants on the *j*th exposure, *j* = 1*,…, p*, across *n* IVs are drawn independently from a multivariate Gaussian distribution, *i.e*.,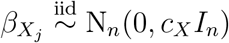, where *c_X_* is a scaling factor related to the sought range of the simulated *β_X_* and defined as the difference between the maximum and minimum desired value of the effect size divided by four. The range of the simulated *β_X_* is chosen between 2 and 2 with 95% of the simulated values within this interval. It also implies a unit variance for the summary-level genetic associations with the exposures. We prepare different scenarios in which the exposures are either independent or correlated by inducing a dependence between exposures by multiplying the (*n* × *p*)-dimensional matrix *β_X_* with the (*p* × *p*)-dimensional matrix *D_X_*, where *D_X_* is the Cholesky decomposition of the Toeplitz matrix *R_X_* with 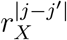 for the (*j, j*′) element of *R_X_*, *j* = *j*′ = 1*,…, p*. The matrix *R_X_* implies a tridiagonal sparse inverse correlation matrix 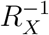. We use different levels of correlation between the exposures, ranging from independence to strong correlation, *i.e*., *r_X_* = 0, 0.2, 0.4, 0.6, 0.8, where *r_X_* = 0.6 represents a medium correlation strength.
- For the second stage of the simulation study, according to eq. (5), an outcome *k*, *k* = 1*,…, q*, is generated on another independent set of *N_Y_* = 50, 000 individuals

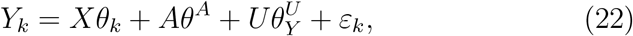

where *θ_k_* is *p*-dimensional vector which contains the direct causal effects of the exposures on the *k*th outcome, *X* is the (*N_X_ × p*)-dimensional matrix of exposures simulated using eq. (21), *θ^A^* and 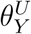 are the effects of the unmeasured pleiotropic pathway *A* and the unmeasured confounder *U* on the same outcome, respectively, and 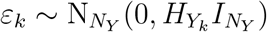, where 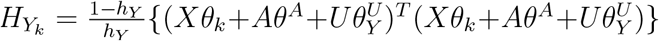 with *h_Y_* the desired level of the proportion of variance explained, fixed at 25% for all outcomes in all simulated scenarios, except for Scenario I-Null setting. We also simulate different second-level data with various values of the pleiotropic effect on the outcomes *θ^A^* = {0.25, 0.50, 0.75, 1.00, 1.50, 2.00}. In the Simulation V-Dependence scenario, we induce the correlation between the outcomes by controlling directly the responses’ error correlation. Specifically, we simulate the errors *∊* = (*∊*_1_*,…, ∊_q_*) for all *k* in eq. (22) from a multivariate Gaussian distribution with mean vector 0 and correlation structure based on the Toeplitz matrix *R_Y_* with 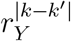 for the (*k, k*′) element of *R_Y_*, *k* = *k*′ = 1*,…, q* and the level of *r_Y_* chosen in the set *r_Y_* = { 0, 0.2, 0.4, 0.6, 0.8 } such that it induces correlation between the responses’ errors

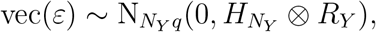

where 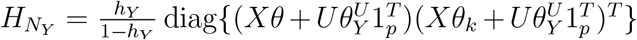 is a *N_Y_* × *N_Y_* diagonal matrix with *θ* = (*θ*_1_*,…, θ_p_*)*^T^* the vector of the direct effect estimates. To evaluate the impact in eq. (22) of the unmeasured shared pleiotropy on the outcomes, we generate the *N_Y_* -dimensional vector *A* as follows

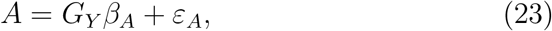

where *β_A_* is the *n*-dimensional vector of genetic associations with the unmeasured pleiotropic pathway *A* drawn from a uniform distribution defined on a range between −2 and 2, *i.e*.,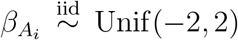, 1,…, *n*, mimicking shared “undirected” pleiotropy, i.e., *A_l_* ≷ 0, *l* = 1,…, *n*. Similarly to the simulation of the exposures, the error term in eq. (23) is simulated controlling the level of variance explained 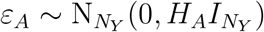, where 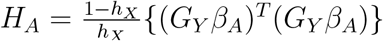. Additionally, we consider shared “directed” pleiotropy, where the effect direction on the pleiotropic pathway *A* is drawn from a uniform distribution defined only on a positive range between 0 and 2, *i.e*., 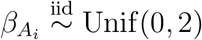. Finally, the direct causal effects *θ_k_*, *k* = 1*,…, q*, are drawn independently from a multivariate Gaussian distribution, 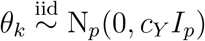, where *c_Y_* is a scaling factor related to the sought range of the simulated *θ_k_* and defined as the difference between the maximum and minimum desired value of the direct causal effects divided by four. For the direct causal effects, the desired interval lies between 2 and 2, implying unit variance for the direct causal effects.

In the simulation study, we include one “null” setting with no direct causal effects. All other settings considered a (*q × p*)-dimensional sparse matrix of direct causal effects Θ = (*θ*_1_*,…, θ_q_*)*^T^*, where 30% cells of the matrix are non-zero and where several exposures are either shared or distinct for the outcomes. Specifically, we select at random the same proportion of cells in the matrix Θ and assign them the simulated values, while the other cells are set to zero. On average, the most likely configuration is with a distinct exposure (37.3%) followed by a shared exposure between two outcomes (31.5%), no direct causal effects (16.6%) and an exposure shared by more than two outcomes (14.5%) (see Table 3 for an overview). On the other hand, the most likely number of associated exposures for each outcome is four (24.4%), five (22.5%), three (18%), six (14.5%), two (8.5%) and seven (6.6%).

After creating data on the individual level, we compute the corresponding summary-level statistics from the two independent groups of individuals. The input data for the simulation are the summary-level statistics *β_X_*, a (*n* × *p*)-dimensional matrix, and *β_Y_*, a (*n* × *q*)-dimensional matrix, derived from a univariable linear regression model where each genetic variant *G_i_*, *i* = 1,…, *n*, is regressed against exposure *X_j_*, *j* = 1*,…, p*, or outcome *Y_k_*, *k* = 1*,…, q*, at-a-time. Additionally, we monitored the average *F* -statistic in the first stage to control for instrument strength. All genetic variants are considered with equal weights, thus no IVW is needed given that the same MAF at 5%.

All simulation runs are repeated 50 times each of which is initialised with a different random seed.

## Acronyms

AUC: Area under the curve
CPO: Conditional predictive ordinate
CVD: Cardiovascular disease
DAG: Directed acyclic graph
EM: Expectation-Maximisation
FDR: False Discovery Rate
GWAS: Genome-wide association study
ePPI: Edge posterior probability inclusion
FPR: False positive rate
IV: Instrument variable
IVW: Inverse variance weighing
MAF: Minor allele frequency
M-H: Metropolis-Hastings
MACE: Model-averaged causal effect estimate
MCMC: Monte carlo Markov Chain
mPPI: Marginal posterior probability of inclusion
MR: Mendelian randomization
MR^2^: Multi-response Mendelian randomization
MR-BMA: Mendelian randomization Bayesian model averaging
MRCE: Multivariate Regression with Covariance Estimation
mSSL: Multiple responses Spike-and-Slab Lasso
MV-MR: Multivariable Mendelian randomisation
OLS: Ordinary least squares
RCT: Randomised clinical trial
ROC: Receiver operating characteristic
SSE: Sum squared error
SUR: Seemingly unrelated regression
TPR: True positive rate
AF: Atrial fibrilation
CAD: Coronary artery disease
CES: Cardioembolic stroke
HF: Heart failure
PAD: Peripheral artery disease
ApoA: Apolipoprotein A
ApoB: Apolipoprotein B
BMI: Body mass index
HDL: High-density lipoprotein
LDL: Low-density lipoprotein
PA: Physical activity
SBP: Systolic blood pressure
SMOKING: Smoking composite index
TG: Triglycerides
T2D: Type 2 diabetes
S.LDL.P: Small large-density lipoprotein particles
M.LDL.P: Medium large-density lipoprotein particles
L.LDL.P: Large-density lipoprotein particles
IDL.P: Intermediate-density lipoprotein particles
XS.VLDL.P: Extra small very-large density lipoprotein particles
S.VLDL.P: Small very-large density lipoprotein particles
M.VLDL.P: Medium very-large density lipoprotein particles
L.VLDL.P: Large very-large density lipoprotein particles
XL.VLDL.P: Extra large very-large density lipoprotein particles
XXL.VLDL.P: Extra-extra large very-large density lipoprotein particles

