## Supplemental Information for "Multi-response Mendelian randomization: Identification of shared and distinct exposures for multimorbidity and multiple related disease outcomes"

[1]Department of Epidemiology and Biostatistics, School of Public Health, Imperial College London, London, UK
[2]MRC Centre for Environment and Health, School of Public Health, Imperial College London, London, UK
[3]UK Dementia Research Institute, Imperial College London, London, UK
[4]Department of Medical Statistics, London School of Hygiene and Tropical Medicine, London, UK
[5]Division of Cardiovascular Medicine, Perelman School of Medicine, University of Pennsylvania, Philadelphia, USA
[6]Department of Medicine, Perelman School of Medicine, University of Pennsylvania, Philadelphia, USA
[7]Corporal Michael J. Crescenz VA Medical Center, Philadelphia, USA
[8]Department of Brain Sciences, Faculty of Medicine, Imperial College London, London, UK
[9]National Heart and Lung Institute, Imperial College London, London, UK
[10]Department of Surgery, Perelman School of Medicine, University of Pennsylvania, Philadelphia, USA
[11]Department of Genetics, Perelman School of Medicine, University of Pennsylvania, Philadelphia, USA
[12]MRC Biostatistics Unit, School of Clinical Medicine, University of Cambridge, Cambridge, UK
[13]Cardiovascular Epidemiology Unit, School of Clinical Medicine, University of Cambridge, Cambridge, UK
[14]Chief Scientific Advisor Office, Research and Early Development, Novo Nordisk, Copenhagen, Denmark
[15]Department of Medical Genetics, School of Clinical Medicine, University of Cambridge, Cambridge, UK
[16]Alan Turing Institute, London, UK

### Contents

**S.1 Multi-response MR residual correlation** **S.4**

**S.2 Details of the MCMC implementation** **S.5**

**S.3 Simulation study** **S.8**
S.3.1 Comparison of methods setup . . . . . . . . . . . . . . . . S.8
S.3.2 Supplemental Tables . . . . . . . . . . . . . . . . . . . . . . S.11
S.3.3 Supplemental Figures . . . . . . . . . . . . . . . . . . . . . S.22

**S.4 Application examples** **S.26**
S.4.1 Data pre-processing . . . . . . . . . . . . . . . . . . . . . . S.26
S.4.2 Analysis plan and comparisons with alternative methods S.27
S.4.3 Supplemental Tables . . . . . . . . . . . . . . . . . . . . . . S.29
S.4.4 Supplemental Figures . . . . . . . . . . . . . . . . . . . . . S.37

### List of Tables

S.1 Overview of MR models that consider one outcome at-a-time and multi-response multivariable methods compared in the simulation study . . . S.11
S.2 Overview of different simulations setting . . . S.12
S.3 Scenario II-Confounding: Area under the curve (AUC) of the simulated scenario . . . S.13
S.4 Scenario III-Undirected pleiotropy: Area under the receiver operating characteristic (AUC) of the simulated scenario . S.14
S.5 Scenario IV-Directed pleiotropy: Area under the receiver operating characteristic (AUC) of the simulated scenario . . . S.15
S.6 Simulation V-Dependence: Area under the receiver operating characteristic (AUC) of the simulated scenario . . . S.16
S.7 Scenario I-Null: Sum of squared errors (SSE) of the simulated scenario . . . S.17
S.8 Scenario II-Confounding: Sum of squared errors (SSE) of the simulated scenario . . . S.18
S.9 Scenario III-Undirected pleiotropy: Sum of squared errors (SSE) of the simulated scenario . . . S.19
S.10 Simulation IV-Directed pleiotropy: Sum of squared errors (SSE) of the simulated scenario . . . S.20
S.11 Simulation V-Dependence: Sum of squared errors (SSE) of the simulated scenario . . . S.21
S.12 Overview of summary-level data in the two application examples . . . S.30
S.13 MV-MR: Application example 1 on common risk factors for cardiovascular disease outcomes . . . S.31
S.14 MR-Egger: Application example 1 on common risk factors for cardiovascular disease outcomes . . . S.32
S.15 MR-BMA: Application example 1 on common risk factors for cardiovascular disease outcomes . . . S.33
S.16 MV-MR: Application example 2 on molecular risk factors for cardiovascular disease outcomes . . . S.34
S.17 MV-Egger: Application example 2 on molecular risk factors for cardiovascular disease outcomes . . . S.35
S.18 MR-BMA: Application example 2 on molecular risk factors for cardiovascular disease outcomes . . . S.36

### List of Figures


S.1 Receiver operating characteristic (ROC) curves in different simulated scenarios when there is no correlation between exposures . . . S.22
S.2 Simulation II-Confounding: Receiver operating characteristic (ROC) curves for different levels of correlation between exposures . . . S.23
S.3 Simulation IV-Directed pleiotropy: Receiver operating characteristic (ROC) curves for different levels of the pleiotropic pathway effect . . . S.24
S.4 Simulation V-Dependence: Receiver operating characteristic (ROC) curves for different levels of the correlation between individual-level responses' errors . . . S.25
S.5 Correlation and partial correlation between summary-level outcomes and between summary-level exposures in application example 1 on common risk factors for cardiovascular disease outcomes . . . S.38
S.6 $MR^2$: Outliers, high-leverage and influential observations detection in application example 1 on common risk factors for cardiovascular disease outcomes . . . S.39
S.7 $MR^2$: Posterior inference of direct causal effects in application example 1 on common risk factors for cardiovascular disease outcomes . . . S.40
S.8 Correlation and partial correlation between exposures and among risk factors in application example 1 on common risk factors for cardiovascular disease outcomes . . . S.41
S.9 $MR^2$: Outliers, high-leverage and influential observations detection in application example 2 on molecular risk factors for cardiovascular disease outcomes . . . S.42
S.10 $MR^2$: Posterior inference of direct causal effects in application example 2 on molecular risk factors for cardiovascular disease outcomes . . . S.43

### S.1 Multi-response MR residual correlation

In eq. (6) of the main paper, $\tilde{\varepsilon}_k$, $k = 1, \ldots, q$, is normally distributed with mean $\mathbb{E}(\tilde{\varepsilon}_k) = 0$ and variance

$$\begin{aligned}
\mathbb{V}(\tilde{\varepsilon}_k) &= (G^T G)^{-1} G^T V(\varepsilon_k) G (G G^T)^{-1} \\
&= (G^T G)^{-1} G^T \delta_k^2 I_N G (G^T G)^{-1} \\
&= \delta_k^2 (G^T G)^{-1}.
\end{aligned} \tag{S.1}$$

In eq. (7), the summary-level error $\epsilon_k$, $k = 1, \ldots, q$, is normally distributed with mean

$$\begin{aligned}
\mathbb{E}(\beta_A \theta^A + \tilde{\varepsilon}_k) &= \mathbb{E}(\beta_A \theta^A) + \mathbb{E}(\tilde{\varepsilon}_k) \\
&= \theta^A (G^T G)^{-1} G^T \, \mathbb{E}(A) + (G^T G)^{-1} G^T \, \mathbb{E}(\varepsilon_k) \\
&= \theta^A \mu_A (G^T G)^{-1} G^T 1_N,
\end{aligned}$$

where $\mathbb{E}(A) = \mu_A 1_N$ with $1_N$ a $N$-dimensional vector of ones.
Assuming $A$ and $\varepsilon_k$ independent in eq. (5) and using eq. (S.1), then

$$\begin{aligned}
\mathbb{V}(\beta_A \theta^A + \tilde{\varepsilon}_k) &= \mathbb{V}(\beta_A \theta^A) + \mathbb{V}(\tilde{\varepsilon}_k) \\
&= (\theta^A)^2 \, \mathbb{V}(\beta_A) + \delta_k^2 (G^T G)^{-1} \\
&= (\theta^A)^2 \sigma_A^2 (G^T G)^{-1} + \delta_k^2 (G^T G)^{-1} \\
&= \{(\theta^A)^2 \sigma_A^2 + \delta_k^2\} (G^T G)^{-1}
\end{aligned} \tag{S.2}$$

with $\mathbb{V}(\beta_A) = (G^T G)^{-1} G^T \sigma_A^2 I_N G (G^T G)^{-1} = \sigma_A^2 (G^T G)^{-1}$.
The correlation between the residuals of the summary-level outcomes $k$ and $k'$, $k \neq k'$, is

$$\begin{aligned}
\rho(\epsilon_k, \epsilon_{k'}) &= \mathbb{V}(\beta_A \theta^A + \tilde{\varepsilon}_k)^{-1/2} \, \mathrm{Cov}(\beta_A \theta^A + \tilde{\varepsilon}_k, \beta_A \theta^A + \tilde{\varepsilon}_{k'}) \times \\
&\quad \mathbb{V}(\beta_A \theta^A + \tilde{\varepsilon}_{k'})^{-1/2} \\
&= \{(\theta^A)^2 \sigma_A^2 + \delta_k^2\}^{-1/2} (G^T G)^{1/2} \times \\
&\quad \mathbb{V}(\beta_A \theta^A) + \mathrm{Cov}(\tilde{\varepsilon}_k, \tilde{\varepsilon}_{k'})] \times \\
&\quad \{(\theta^A)^2 \sigma_A^2 + \delta_{k'}^2\}^{-1/2} (G^T G)^{1/2} \\
&= \{(\theta^A)^2 \sigma_A^2 + \delta_k^2\}^{-1/2} \times \\
&\quad (G^T G)^{1/2} [(\theta^A)^2 \sigma_A^2 (G^T G)^{-1} + \sigma_{kk'} (G^T G)^{-1}] \times \\
&\quad \{(\theta^A)^2 \sigma_A^2 + \delta_{k'}^2\}^{-1/2} (G^T G)^{1/2} \\
&= \frac{(\theta^A)^2 \sigma_A^2 + \sigma_{kk'}}{\{(\theta^A)^2 \sigma_A^2 + \delta_k^2\}^{1/2} \{(\theta^A)^2 \sigma_A^2 + \delta_{k'}^2\}^{1/2}} I_n
\end{aligned} \tag{S.3}$$

using (S.2) and the fact that

$$\begin{aligned}\mathrm{Cov}(\tilde{\varepsilon}_k, \tilde{\varepsilon}_{k'}) &= (G^T G)^{-1} G^T \sigma_{kk'} I_N G (G G^T)^{-1} \\ &= \sigma_{kk'} (G^T G)^{-1} G^T G (G G^T)^{-1} \\ &= \sigma_{kk'} (G^T G)^{-1},\end{aligned}$$

where $\sigma_{kk'} = \mathrm{Cov}(\varepsilon_k, \varepsilon_{k'})$ is the covariance between individual-level responses' errors which is assumed constant across all individuals.

Considering all outcomes and all IVs, the summary-level covariance matrix can be written in compact form as $\rho(\epsilon) = I_n \otimes R$ with $\epsilon = (\epsilon_1, \ldots, \epsilon_q)$ and $R_{kk'}$ as in eq. (S.3) for any of the $n$ IVs.

### S.2 Details of the MCMC implementation

The MCMC algorithm for the selection of important predictors in regression models with multiple responses of any type is presented in [1]. Here, we describe the main steps of the MCMC algorithm for the proposed multi-response multivariable Mendelian randomization $\mathrm{MR}^2$ model as well as the main advantages of the designed proposal distribution for the non-zero direct causal effects $\theta_k^*$ over standard implementations of Bayesian inference for the SUR model. In the following, the symbol "*" indicates the proposed new value used in the Metropolis-Hastings (M-H) acceptance ratio.

Given a candidate value of the latent binary vector $\gamma_k^*$, the proposal density for $\theta_k^*$ is

$$q(\theta_k^* | \gamma_k^*, \beta_{Y_k}, Z_k, \Delta, R) \propto \mathcal{N}_{|\gamma_k^*|}(\theta_k^*; m_{\gamma_k^*}, V_{\gamma_k^*}), \tag{S.4}$$

where

- $V_{\gamma_k^*}^{-1} = \delta_{k|-k}^{-2} (\beta_X^T)_{\gamma_k^*} (\beta_X)_{\gamma_k^*} + v^{-1} I_{|\gamma_k^*|}$ is the $(|\gamma_k^*| \times |\gamma_k^*|)$-dimensional proposal precision matrix,
- $m_{\gamma_k^*} = V_{\gamma_k^*} (\beta_X^T)_{\gamma_k^*} (\beta_{Y_k} - \delta_k \mu_{k|-k}) / \delta_{k|-k}^2$ is the $|\gamma_k^*|$-dimensional proposal mean,
- $\delta_{k|-k}^2$ is the conditional variance of $\beta_{Y_k} | Z_{-k}$, *i.e.*, $\delta_{k|-k}^2 = \delta_k^2 / (R^{-1})_{kk}$,
- $\mu_{k|-k}$ is the $n$-dimensional vector of conditional means of $Z_k | Z_{-k}$, *i.e.*, $\mu_{k|-k} = Z_{-k} R_{-k,-k}^{-1} R_{-k,k}$

and where the subscript "$-k$" implies that the corresponding matrix (or vector) consists of all the elements except those related to the $k$th response.

The interpretation of the above quantities sheds light on the benefits of the proposed model. $V_{\gamma_k^*}^{-1}$ resembles the standard Gibbs sampling update of the regression coefficients precision matrix in the linear model. However, in $V^{-1}_{\gamma^*_k}$, we use the conditional variance $\delta^2_{k|-k}$ of $\beta_{Y_k}|Z_{-k}$ instead of the marginal variance $\delta^2_k$. In $m_{\gamma^*_k}$, since $\mathbb{E}(\beta_{Y_k}) = (\beta_X)_{\gamma^*_k}(\theta_k)_{\gamma^*_k} + \delta_k Z_{-k} R^{-1}_{-k,-k} R_{-k,k} = (\beta_X)_{\gamma^*_k}(\theta_k)_{\gamma^*_k} + \delta_k \mu_{k|-k}$, then $\mathbb{E}(\beta_{Y_k} - \delta_k \mu_{k|-k}) = (\beta_X)_{\gamma^*_k}(\theta_k)_{\gamma^*_k}$. Thus, we recenter the genetic associations with the $k$th outcome such that, once conditioned on $\gamma^*_k$, they depend only on $(\beta_X)_{\gamma^*_k}(\theta_k)_{\gamma^*_k}$ as in eq. (12). The other terms in $m_{\gamma^*_k}$ mirror the mean of the full conditional for the regression coefficients of the linear model. In summary, the proposal density $q(\theta^*_k|\gamma^*_k, \beta_{Y_k}, Z_{_k}, \Delta, R)$ is inspired by the Gibbs updates of the regression coefficients in the linear model with suitable modifications to account for the conditional dependence between the $k$th outcome and the remaining responses at the summary level.

The designed proposal distribution allows also the "implicit marginalisation" of the direct causal effects when the joint update of $(\theta_k, \gamma_k)$, $k = 1, \ldots, q$, in the M-H step is performed. For the $k$th response, the acceptance probability $\alpha$ of the joint proposal is

$$\alpha = 1 \wedge \frac{\left|V_{\gamma^*_k}\right|^{1/2} v^{|\gamma_k|/2} \exp\left(\frac{1}{2} m^T_{\gamma^*_k} V^{-1}_{\gamma^*_k} m_{\gamma^*_k}\right) p(\gamma^*_k) q(\gamma_k|\gamma^*_k)}{\left|V_{\gamma_k}\right|^{1/2} v^{|\gamma^*_k|/2} \exp\left(\frac{1}{2} m^T_{\gamma_k} V^{-1}_{\gamma_k} m_{\gamma_k}\right) p(\gamma_k) q(\gamma^*_k|\gamma_k)}, \tag{S.5}$$

where $p(\gamma_k)$ is defined in eq. (15) and $q(\gamma^*_k|\gamma_k)$ is the proposal distribution for the latent binary vector $\gamma^*_k$ conditional to the current value $\gamma_k$. Similarly to [2], in eq. (S.5), the current and proposed value of the direct causal effects $(\theta_k, \theta^*_k)$ do not appear and the acceptance probability resembles the Bayes Factor of the standard linear regression model used in the update of the latent binary vector $\gamma_k$.

The advantages of the "implicit marginalisation" of the regression coefficients in eq. (S.5) for the SUR model have not been recognised until recently [1]. This is a key point that distinguishes our prosed methodology from other SUR models [3, 4]. Similarly to MR$^2$, a sparse Bayesian SUR model [5] proposes two computational algorithms, an MCMC algorithm using Gibbs updates [6] for variable selection called "undirected" and a "directed" algorithm involving a Monte Carlo numerical approximation of the marginal likelihood to integrate out the regression coefficients combined with an M-H algorithm similar to eq. (S.5) for posterior models' exploration. However, Monte Carlo numerical approximation is unstable and very time-consuming, making [5] unfeasible even for small data sets.

We conclude with the description regarding the MCMC steps required to obtain samples from $p(\Sigma|W, \mathrm{G})$ and transform them using the inverse transformation $R = \Delta^{-1}\Sigma\Delta^{-1}$, where $\Delta$ is sampled from its prior distribution in eq. (18). To sample the symmetric adjacency matrix G from

$p(\mathrm{G}|\beta_Y, Z, \Delta, R)$, we target the distribution with density

$$p(\mathrm{G}|W) = \frac{p(W|\mathrm{G})p(\mathrm{G})}{\sum_{\mathrm{G}} p(W|\mathrm{G})p(\mathrm{G})}, \tag{S.6}$$

where the summation in the denominator is over all the decomposable graphs G, $p(\mathrm{G})$ is defined in (17), $W$ is defined in Appendix and $p(W|\mathrm{G})$ can be computed analytically due to the tractability of the hyper-inverse Wishart distribution [7]. Specifically, we sample from eq. (S.6) by using an M-H step in which, conditionally on the current symmetric adjacency matrix G, a new graph is proposed by adding or deleting an edge between two vertices whose index has been chosen randomly between the vertices that belong to a decomposable graph. The proposed graph is then accepted or rejected using the accept/reject mechanism of the M-H step which targets the density in eq. (S.6). Finally, conditionally on $W$ and the updated symmetric adjacency matrix G, we sample $\Sigma$ from its conditional hyper-inverse Wishart distribution, *i.e.*, $\mathrm{HIW}_{\mathcal{G}}(2+n, I_q + W^T W)$.

#### S.3 Simulation study

##### S.3.1 Comparison of methods setup

###### $MR^2$ parameters setting

The $MR^2$ algorithm is run for $7,500$ sweeps of which $2,500$ as burn-in and results saved every 5 sweeps, resulting in $1,000$ posterior samples for all the unknowns. The hyper-parameters $a_k$ and $b_k$ in eq. (15) are chosen by specifying $\mathbb{E}(\gamma_k) = 2$ and $\mathbb{V}(\gamma_k) = 2$ which implies *a priori* a range between zero and eight for the number of significant direct causal effects for each outcome, see [1] for details. Another hyper-parameter to be specified is the prior variance $v$ in eq. (13). We set $v = 1$ since the summary-level genetic associations with the risk factors in eq. (21) have been simulated with unit variance and no IVW is required given the same MAF for all simulated genotypes.

In case of different MAF across genotypes, we recommend setting $v = 1$ and standardise the summary-level genetic associations with the risk factors after IVW in order to place appropriate prior mass on reasonable values of the non-zero direct causal effects [8].

###### Alternative methods and parameters setting

As alternative methods, we include standard multivariable MR (MV-MR) which considers multiple exposures in one joint model but does not perform variable selection [9]. Next, we consider MR-BMA [10], a Bayesian variable selection approach for multivariable MR, with a prior probability of inclusion set at $2/15 = 0.13$ reflecting two associated causal risk factors for each outcome. In this way, we match the prior specification of the expected number of risk factors associated with the $k$th response used in $MR^2$. Both MV-MR and MR-BMA only consider one outcome at-a-time and are performed on each outcome separately.

Additionally, we include two multivariate and multivariable variable selection approaches which have to date not been applied to MR, the multiple responses Lasso (Multivariate Regression With Covariance Estimation, MRCE) [3] and the multiple responses Spike and Slab Lasso (mSSL) [4].

Regarding the MRCE algorithm, we select the cross-validation procedure option for the sparse Lasso solution which performs $k$-fold cross-validation using candidate tuning parameters specified in the vectors $\lambda_1$ and $\lambda_2$, the two penalisation parameters for the sparse regression and sparse covariance estimation, respectively. We run the algorithm with the default recommended values including 5-fold cross-validation and 40 equally spaced candidate tuning parameters $\lambda_1$ and $\lambda_2$ between 0 and 1. Results (not shown) do not change if a finer grid of points for $\lambda_1$ and $\lambda_2$ are chosen.

We use the default parametrization for the instance of the mSSL algorithm that performs "dynamic posterior exploration" (mSSL_dpe) which is the slowest, but the most accurate version of the algorithm. As recommended, we also run the "dynamic conditional posterior exploration" (mSSL_dcpe) and "separate variable and covariance selection" (sep_sslg) versions of the algorithm, but with inferior results (data not shown).

A unit variance of the predictors is required by both MRCE and mSSL algorithms. This condition is automatically satisfied since, in all simulated scenarios, the summary-level genetic associations with the risk factors in eq. (21) have unit variance. The simulation setup also simplifies the choice of prior variance of the non-zero regression coefficients used in MR-BMA. We use the same specification ($v = 1$) employed in MR$^2$. The output of MRCE and mSSL algorithms consists of the sparse Lasso solution of the direct causal effects $\Theta = (\theta_1, \ldots, \theta_q)^T$ and residual covariance $\Sigma$, or its inverse $\Omega$, including the intercepts for each outcome. Thus, both MRCE and mSSL can be seen as the multi-response versions, with exposure selection, of multivariable MR-Egger [11], see Supplemental Table S.1. However, in the designed simulation setup, the unmeasured pleiotropic pathway is shared across responses and it induces residual correlation at the summary-level data as shown in eq. (5). Thus, its effect is detected by the estimated residual covariance $\Sigma$ with the estimated intercepts close to zero (data not shown). Similar considerations hold when the residual correlation is induced by directly modelling the correlation between the individual-level responses' errors.

##### Performance measures

There are two important aspects when comparing methods for multivariable MR. First, the ability of the algorithms to detect the true causal risk factors and, second, the accuracy of the direct causal effect estimates. Receiver operating characteristic (ROC) curves plot the true positive rate (TPR) against the false positive rate (FPR) and illustrate how well the methods can distinguish between true and false causal exposures when the discrimination threshold is varied. As discrimination threshold, in MV-MR we rank according to $p$-values of the direct causal effect estimates and in MR-BMA and MR$^2$ according to mPPI. The two multi-response implementations of Lasso rank exposures by their causal effect estimates of which the unimportant ones are forced to zero and thus excluded from the model. Consequently, these two methods do not provide a full ranking of all exposures, but rather a single threshold. Thus, they are presented in the ROC curves as a single point instead of a continuous line.

Second, to contrast the quality of the direct causal effect estimation, we calculate the sum of squared errors (SSE), defined as the squared difference between the estimated and the true simulated direct causal effect. The SSE captures both the squared bias and the variance, thus penalizing methods with a wide spread and high variability.

##### S.3.2 Supplemental Tables

| | Designed for MR | Bayesian method | Multivariate outcomes | Selection risk factors | Estimation residual correlation | Reference |
|---|---|---|---|---|---|---|
| MV-MR | ✓ | ✗ | ✗ | ✗ | ✗ | Burgess *et al.* 2015 [9] |
| MR-BMA | ✓ | ✓ | ✗ | ✓ | ✗ | Zuber *et al.* 2020 [10] |
| MRCE | ✗ | ✓ | ✓ | ✓ | ✓ | Rothman *et al.* 2010 [3] |
| mSSL | ✗ | ✓ | ✓ | ✓ | ✓ | Deshpande *et al.* 2019 [4] |
| $MR^2$ | ✓ | ✓ | ✓ | ✓ | ✓ | |

**Table S.1. Overview of competing multivariable MR models that consider one outcome at-a-time and multi-response multivariable methods used in the simulation study.** For each method, we indicate if: It has been designed specifically for MR, it is a Bayesian method, it can analyse multiple outcomes jointly, it can provide selection of important risk factors associated with the outcomes and it can estimate residual correlation between summary-level responses.

| Scenario | Direct causal effect | Correlation among residuals | Further notes | Open parameters |
|---|---|---|---|---|
| I-Null | $\theta_k = 0$ | No | Null direct causal effect of all exposures, only the confounders have an effect | Impact of confounder: $\theta_X^U = 2$, $\theta_Y^U = 1$ |
| II-Confounding | $\theta_k \neq 0$ | No | Direct causal effect on randomly selected exposure-confounder pairs | Impact of confounder: $\theta_X^U = 2$, $\theta_Y^U = 1$ |
| III-Undirected pleiotropy | $\theta_k \neq 0$ | Yes, induced by pleiotropic pathway | Additional effect of shared pleiotropic pathway $A$ ("undirected": $A_l \gtrless 0$) on the level of outcomes | Pleiotropic effect: $\theta^A = \{0.25, 0.5, 0.75, 1, 1.5, 2\}$ |
| IV-Directed pleiotropy | $\theta_k \neq 0$ | Yes, induced by pleiotropic pathway | Additional effect of shared pleiotropic pathway $A$ ("directed": $A_l > 0$) on the level of outcomes | Pleiotropic effect: $\theta^A = \{0.25, 0.5, 0.75, 1, 1.5, 2\}$ |
| V-Dependence | $\theta_k \neq 0$ | Yes, modelled direclty | Correlation between individual-level responses' errors in eq. (5) | Correlation level: $r_Y = \{0, 0.2, 0.4, 0.6, 0.8\}$ |

**Table S.2. Overview of different simulations setting.** Additional open parameter is the correlation between exposures $r_X = \{0, 0.2, 0.4, 0.6, 0.8\}$. Fixed parameters are the sample size of independent subjects ($N = 50,000$) for the first (risk factors) and second (responses) level of the simulations, the number of genetic variants used as instrumental variables ($n = 100$), the range of sought values for the exposure effects ($\beta_{X_j}$, $j = 1, \ldots, p$, should be in the range between $-2$ and 2), the percentage of simulated non-zero direct causal effects among the exposure-outcome combinations (30%), the range of sought values for the direct causal effects ($\theta_k$, $k = 1, \ldots, q$, should be in the range between $-2$ and 2) and the proportion of variance explained to generate $p = 15$ exposures ($h_X = 10\%$) and each of $q = 5$ outcomes ($h_Y = 25\%$).

| | $\theta_X^U$ | | $\theta_Y^U$ | |
|---|---|---|---|---|
| | | | 1 | 2 |
| $r_X$ = 0 | 1 | MV-MR | 0.958 (0.026) | 0.958 (0.026) |
| | | MR-BMA | 0.960 (0.025) | 0.961 (0.025) |
| | | MRCE | 0.959 (0.025) | 0.959 (0.026) |
| | | mSSL | 0.901 (0.072) | 0.909 (0.048) |
| | | $MR^2$ | **0.961** (0.025) | **0.962** (0.025) |
| | 2 | MV-MR | 0.951 (0.030) | 0.952 (0.030) |
| | | MR-BMA | 0.956 (0.028) | 0.956 (0.029) |
| | | MRCE | 0.952 (0.036) | 0.953 (0.037) |
| | | mSSL | 0.879 (0.067) | 0.892 (0.044) |
| | | $MR^2$ | **0.961** (0.025) | **0.960** (0.025) |
| 0.2 | 1 | MV-MR | 0.958 (0.024) | 0.959 (0.024) |
| | | MR-BMA | 0.962 (0.025) | 0.962 (0.025) |
| | | MRCE | 0.957 (0.029) | 0.958 (0.026) |
| | | mSSL | 0.906 (0.048) | 0.901 (0.050) |
| | | $MR^2$ | **0.964** (0.023) | **0.964** (0.025) |
| | 2 | MV-MR | 0.951 (0.027) | 0.951 (0.027) |
| | | MR-BMA | **0.957** (0.027) | 0.956 (0.027) |
| | | MRCE | 0.952 (0.027) | 0.950 (0.029) |
| | | mSSL | 0.886 (0.071) | 0.888 (0.039) |
| | | $MR^2$ | **0.957** (0.026) | **0.958** (0.024) |
| 0.4 | 1 | MV-MR | 0.946 (0.041) | 0.946 (0.041) |
| | | MR-BMA | 0.955 (0.034) | 0.956 (0.034) |
| | | MRCE | 0.948 (0.038) | 0.947 (0.039) |
| | | mSSL | 0.899 (0.050) | 0.897 (0.043) |
| | | $MR^2$ | **0.962** (0.028) | **0.960** (0.032) |
| | 2 | MV-MR | 0.940 (0.042) | 0.937 (0.042) |
| | | MR-BMA | 0.950 (0.038) | 0.949 (0.037) |
| | | MRCE | 0.943 (0.040) | 0.942 (0.039) |
| | | mSSL | 0.870 (0.060) | 0.880 (0.051) |
| | | $MR^2$ | **0.953** (0.037) | **0.953** (0.035) |
| 0.6 | 1 | MV-MR | 0.928 (0.038) | 0.930 (0.038) |
| | | MR-BMA | 0.940 (0.031) | 0.941 (0.031) |
| | | MRCE | 0.936 (0.034) | 0.937 (0.032) |
| | | mSSL | 0.884 (0.047) | 0.893 (0.048) |
| | | $MR^2$ | **0.948** (0.030) | **0.946** (0.029) |
| | 2 | MV-MR | 0.922 (0.040) | 0.924 (0.038) |
| | | MR-BMA | 0.932 (0.034) | 0.934 (0.035) |
| | | MRCE | 0.926 (0.035) | 0.927 (0.037) |
| | | mSSL | 0.876 (0.044) | 0.880 (0.046) |
| | | $MR^2$ | **0.939** (0.033) | **0.944** (0.029) |
| 0.8 | 1 | MV-MR | 0.909 (0.034) | 0.909 (0.033) |
| | | MR-BMA | 0.924 (0.040) | 0.925 (0.040) |
| | | MRCE | 0.914 (0.039) | 0.916 (0.039) |
| | | mSSL | 0.867 (0.058) | 0.870 (0.048) |
| | | $MR^2$ | **0.932** (0.035) | **0.937** (0.036) |
| | 2 | MV-MR | 0.896 (0.037) | 0.899 (0.038) |
| | | MR-BMA | 0.916 (0.041) | 0.919 (0.041) |
| | | MRCE | 0.900 (0.048) | 0.904 (0.044) |
| | | mSSL | 0.846 (0.058) | 0.851 (0.052) |
| | | $MR^2$ | **0.927** (0.038) | **0.932** (0.037) |

**Table S.3. Scenario II-Confounding: Area under the receiver operating characteristic (AUC) of the simulated scenario**, averaged over 50 replicates, with standard deviation in brackets, illustrating the performance of different MR multivariable implementations and multi-response multivariable methods. In simulated Scenario II-Confounding, 30% of all exposure-confounder combinations are generated to have a direct causal effect. Open parameters are the correlation between exposures $r_X = \{0, 0.2, 0.4, 0.6, 0.8\}$ and the confounding effects on the exposures $\theta_X^U = \{1, 2\}$ and outcomes $\theta_Y^U = \{1, 2\}$. Best results are highlighted in bold.

| | | $\theta^A$ | | | | | |
|---|---|---|---|---|---|---|---|
| | | 0.25 | 0.5 | 0.75 | 1 | 1.5 | 2 |
| $r_X$ 0 | MV-MR | 0.943 (0.035) | 0.931 (0.038) | 0.917 (0.040) | 0.900 (0.043) | 0.857 (0.051) | 0.818 (0.056) |
| | MR-BMA | 0.947 (0.033) | 0.937 (0.034) | 0.924 (0.039) | 0.907 (0.040) | 0.863 (0.050) | 0.826 (0.055) |
| | MRCE | 0.946 (0.038) | 0.943 (0.034) | 0.934 (0.038) | 0.928 (0.040) | **0.918** (0.037) | 0.903 (0.044) |
| | mSSL | 0.895 (0.052) | 0.880 (0.057) | 0.876 (0.053) | 0.869 (0.062) | 0.846 (0.055) | 0.827 (0.063) |
| | $MR^2$ | **0.952** (0.035) | **0.946** (0.038) | **0.938** (0.033) | **0.933** (0.033) | 0.915 (0.040) | **0.904** (0.045) |
| 0.2 | MV-MR | 0.947 (0.034) | 0.931 (0.039) | 0.912 (0.048) | 0.889 (0.054) | 0.850 (0.061) | 0.804 (0.076) |
| | MR-BMA | 0.948 (0.034) | 0.933 (0.037) | 0.916 (0.043) | 0.893 (0.051) | 0.853 (0.064) | 0.813 (0.072) |
| | MRCE | 0.949 (0.035) | 0.944 (0.033) | 0.942 (0.034) | 0.934 (0.031) | **0.923** (0.037) | **0.913** (0.042) |
| | mSSL | 0.889 (0.050) | 0.865 (0.069) | 0.865 (0.060) | 0.860 (0.059) | 0.843 (0.057) | 0.823 (0.067) |
| | $MR^2$ | **0.950** (0.033) | **0.948** (0.029) | **0.940** (0.035) | **0.940** (0.029) | 0.922 (0.039) | 0.911 (0.039) |
| 0.4 | MV-MR | 0.941 (0.036) | 0.923 (0.042) | 0.900 (0.050) | 0.877 (0.054) | 0.832 (0.056) | 0.783 (0.065) |
| | MR-BMA | 0.947 (0.034) | 0.934 (0.036) | 0.916 (0.043) | 0.895 (0.052) | 0.852 (0.057) | 0.806 (0.063) |
| | MRCE | 0.943 (0.036) | 0.936 (0.036) | 0.929 (0.039) | 0.923 (0.044) | 0.912 (0.048) | **0.892** (0.051) |
| | mSSL | 0.889 (0.058) | 0.874 (0.058) | 0.859 (0.060) | 0.851 (0.054) | 0.832 (0.060) | 0.811 (0.062) |
| | $MR^2$ | **0.948** (0.035) | **0.946** (0.036) | **0.934** (0.042) | **0.926** (0.044) | **0.914** (0.048) | 0.889 (0.056) |
| 0.6 | MV-MR | 0.919 (0.044) | 0.903 (0.047) | 0.881 (0.052) | 0.857 (0.057) | 0.804 (0.070) | 0.756 (0.078) |
| | MR-BMA | 0.933 (0.037) | 0.919 (0.044) | 0.901 (0.050) | 0.876 (0.058) | 0.828 (0.062) | 0.791 (0.071) |
| | MRCE | 0.923 (0.041) | 0.917 (0.041) | 0.906 (0.047) | 0.904 (0.048) | 0.887 (0.048) | 0.874 (0.051) |
| | mSSL | 0.868 (0.056) | 0.859 (0.053) | 0.850 (0.060) | 0.840 (0.055) | 0.820 (0.058) | 0.796 (0.072) |
| | $MR^2$ | **0.940** (0.033) | **0.929** (0.039) | **0.920** (0.039) | **0.917** (0.043) | **0.895** (0.043) | **0.883** (0.050) |
| 0.8 | MV-MR | 0.885 (0.053) | 0.852 (0.059) | 0.818 (0.069) | 0.793 (0.076) | 0.738 (0.081) | 0.703 (0.084) |
| | MR-BMA | 0.895 (0.042) | 0.874 (0.046) | 0.850 (0.050) | 0.826 (0.055) | 0.780 (0.067) | 0.747 (0.072) |
| | MRCE | 0.887 (0.047) | 0.880 (0.050) | 0.874 (0.052) | 0.867 (0.052) | 0.852 (0.060) | 0.834 (0.066) |
| | mSSL | 0.842 (0.061) | 0.827 (0.071) | 0.807 (0.068) | 0.797 (0.057) | 0.766 (0.083) | 0.748 (0.075) |
| | $MR^2$ | **0.912** (0.043) | **0.905** (0.038) | **0.893** (0.044) | **0.881** (0.048) | **0.855** (0.054) | **0.843** (0.048) |

**Table S.5. Simulation IV-Directed pleiotropy: Area under the receiver operating characteristic (AUC) of the simulated scenario**, averaged over 50 replicates, with standard deviation in brackets, illustrating the performance of different MR multivariable implementations and multi-response multivariable methods. In simulated Scenario IV-Directed pleiotropy, 30% of all exposure-confounder combinations are generated to have a direct causal effect. Open parameters are the correlation between exposures $r_X = \{0, 0.2, 0.4, 0.6, 0.8\}$ and the pleiotropic effect $\theta^A = \{0.25, 0.5, 0.75, 1, 1.5, 2\}$. Confounding effects on the exposures and outcomes are fixed at $\theta_X^U = 2$ and $\theta_Y^U = 1$, respectively. Best results are highlighted in bold.

| | | $r_Y$ | | | | |
|---|---|---|---|---|---|---|
| | | 0 | 0.2 | 0.4 | 0.6 | 0.8 |
| $r_X$ | | | | | | |
| 0 | MV-MR | 0.951 (0.030) | 0.951 (0.028) | 0.953 (0.026) | 0.950 (0.026) | 0.954 (0.023) |
| | MR-BMA | 0.956 (0.028) | 0.954 (0.029) | 0.956 (0.026) | 0.955 (0.026) | 0.956 (0.023) |
| | MRCE | 0.951 (0.037) | 0.951 (0.033) | 0.951 (0.033) | 0.953 (0.028) | 0.955 (0.024) |
| | mSSL | 0.886 (0.068) | 0.893 (0.069) | 0.886 (0.063) | 0.884 (0.069) | 0.896 (0.069) |
| | MR$^2$ | **0.958** (0.024) | **0.962** (0.024) | **0.961** (0.027) | **0.962** (0.024) | **0.967** (0.020) |
| 0.2 | MV-MR | 0.951 (0.027) | 0.949 (0.028) | 0.951 (0.030) | 0.951 (0.032) | 0.950 (0.032) |
| | MR-BMA | **0.957** (0.027) | **0.957** (0.028) | **0.958** (0.029) | 0.957 (0.029) | 0.955 (0.030) |
| | MRCE | 0.953 (0.028) | 0.953 (0.027) | 0.954 (0.029) | 0.952 (0.030) | 0.956 (0.031) |
| | mSSL | 0.890 (0.065) | 0.900 (0.067) | 0.888 (0.071) | 0.894 (0.082) | 0.887 (0.085) |
| | MR$^2$ | **0.957** (0.026) | 0.956 (0.025) | 0.955 (0.029) | **0.960** (0.027) | **0.968** (0.027) |
| 0.4 | MV-MR | 0.940 (0.042) | 0.949 (0.030) | 0.944 (0.034) | 0.946 (0.032) | 0.948 (0.029) |
| | MR-BMA | 0.950 (0.038) | 0.953 (0.029) | 0.952 (0.033) | 0.953 (0.032) | 0.956 (0.031) |
| | MRCE | 0.942 (0.041) | 0.948 (0.032) | 0.943 (0.038) | 0.945 (0.036) | 0.949 (0.035) |
| | mSSL | 0.880 (0.050) | 0.885 (0.051) | 0.877 (0.064) | 0.885 (0.070) | 0.889 (0.075) |
| | MR$^2$ | **0.953** (0.037) | **0.958** (0.026) | **0.956** (0.029) | **0.957** (0.031) | **0.961** (0.029) |
| 0.6 | MV-MR | 0.922 (0.040) | 0.928 (0.037) | 0.925 (0.040) | 0.930 (0.037) | 0.930 (0.044) |
| | MR-BMA | 0.932 (0.034) | 0.935 (0.040) | 0.934 (0.037) | 0.936 (0.032) | 0.935 (0.033) |
| | MRCE | 0.924 (0.036) | 0.933 (0.040) | 0.927 (0.042) | 0.932 (0.037) | 0.930 (0.046) |
| | mSSL | 0.878 (0.066) | 0.870 (0.054) | 0.875 (0.053) | 0.869 (0.067) | 0.878 (0.079) |
| | MR$^2$ | **0.939** (0.033) | **0.943** (0.033) | **0.943** (0.036) | **0.952** (0.031) | **0.951** (0.029) |
| 0.8 | MV-MR | 0.896 (0.037) | 0.900 (0.039) | 0.901 (0.041) | 0.902 (0.039) | 0.901 (0.037) |
| | MR-BMA | 0.916 (0.041) | 0.909 (0.039) | 0.912 (0.037) | 0.911 (0.039) | 0.910 (0.038) |
| | MRCE | 0.904 (0.043) | 0.899 (0.049) | 0.901 (0.042) | 0.903 (0.041) | 0.909 (0.042) |
| | mSSL | 0.849 (0.059) | 0.821 (0.082) | 0.843 (0.073) | 0.855 (0.062) | 0.835 (0.088) |
| | MR$^2$ | **0.927** (0.038) | **0.922** (0.032) | **0.926** (0.035) | **0.934** (0.030) | **0.936** (0.035) |

**Table S.6. Simulation V-Dependence: Area under the receiver operating characteristic (AUC) of the simulated scenario**, averaged over 50 replicates, with standard deviation in brackets, illustrating the performance of different MR multivariable implementations and multi-response multivariable methods. In simulated Scenario V-Dependence, 30% of all exposure-confounder combinations are generated to have a direct causal effect. Open parameters are the correlation between exposures $r_X = \{0, 0.2, 0.4, 0.6, 0.8\}$ and the correlation between individual-level responses' errors $r_Y = \{0, 0.2, 0.4, 0.6, 0.8\}$. Confounding effects on the exposures and outcomes are fixed at $\theta_X^U = 2$ and $\theta_Y^U = 1$, respectively. Best results are highlighted in bold.

| | | | $r_Y$ | | | | |
|---|---|---|---|---|---|---|---|
| | | | 0 | 0.2 | 0.4 | 0.6 | 0.8 |
| $r_X$ | 0 | MV-MR | 0.001 (<0.001) | 0.001 (<0.001) | 0.001 (<0.001) | 0.001 (<0.001) | 0.001 (<0.001) |
| | | MR-BMA | **0.000** (<0.001) | **0.000** (<0.001) | **0.000** (<0.001) | **0.000** (<0.001) | **0.000** (<0.001) |
| | | MRCE | **0.000** (<0.001) | **0.000** (<0.001) | **0.000** (<0.001) | **0.000** (<0.001) | **0.000** (<0.001) |
| | | mSSL | **0.000** (<0.001) | **0.000** (<0.001) | **0.000** (<0.001) | **0.000** (<0.001) | **0.000** (<0.001) |
| | | $MR^2$ | **0.000** (<0.001) | **0.000** (<0.001) | **0.000** (<0.001) | **0.000** (<0.001) | **0.000** (<0.001) |
| | 0.2 | MV-MR | 0.001 (<0.001) | 0.001 (<0.001) | 0.001 (<0.001) | 0.001 (<0.001) | 0.001 (<0.001) |
| | | MR-BMA | **0.000** (<0.001) | **0.000** (<0.001) | **0.000** (<0.001) | **0.000** (<0.001) | **0.000** (<0.001) |
| | | MRCE | **0.000** (<0.001) | **0.000** (<0.001) | **0.000** (<0.001) | **0.000** (<0.001) | **0.000** (<0.001) |
| | | mSSL | **0.000** (<0.001) | **0.000** (<0.001) | **0.000** (<0.001) | **0.000** (<0.001) | **0.000** (<0.001) |
| | | $MR^2$ | **0.000** (<0.001) | **0.000** (<0.001) | **0.000** (<0.001) | **0.000** (<0.001) | **0.000** (<0.001) |
| | 0.4 | MV-MR | 0.001 (<0.001) | 0.001 (<0.001) | 0.001 (<0.001) | 0.001 (<0.001) | 0.001 (<0.001) |
| | | MR-BMA | **0.000** (<0.001) | **0.000** (<0.001) | **0.000** (<0.001) | **0.000** (<0.001) | **0.000** (<0.001) |
| | | MRCE | 0.001 (<0.001) | 0.001 (<0.001) | 0.001 (<0.001) | 0.001 (<0.001) | 0.001 (<0.001) |
| | | mSSL | **0.000** (<0.001) | **0.000** (<0.001) | **0.000** (<0.001) | **0.000** (<0.001) | **0.000** (<0.001) |
| | | $MR^2$ | **0.000** (<0.001) | **0.000** (<0.001) | **0.000** (<0.001) | **0.000** (<0.001) | **0.000** (<0.001) |
| | 0.6 | MV-MR | 0.001 (<0.001) | 0.001 (<0.001) | 0.001 (<0.001) | 0.001 (<0.001) | 0.001 (<0.001) |
| | | MR-BMA | **0.000** (<0.001) | **0.000** (<0.001) | **0.000** (<0.001) | **0.000** (<0.001) | **0.000** (<0.001) |
| | | MRCE | 0.001 (<0.001) | 0.001 (<0.001) | 0.001 (<0.001) | 0.001 (<0.001) | 0.001 (<0.001) |
| | | mSSL | **0.000** (<0.001) | **0.000** (<0.001) | **0.000** (<0.001) | **0.000** (<0.001) | **0.000** (<0.001) |
| | | $MR^2$ | **0.000** (<0.001) | **0.000** (<0.001) | **0.000** (<0.001) | **0.000** (<0.001) | **0.000** (<0.001) |
| | 0.8 | MV-MR | 0.003 (<0.001) | 0.003 (0.001) | 0.003 (0.001) | 0.003 (0.001) | 0.003 (0.001) |
| | | MR-BMA | **0.000** (<0.001) | **0.000** (<0.001) | **0.000** (<0.001) | **0.000** (<0.001) | **0.000** (<0.001) |
| | | MRCE | 0.001 (<0.001) | 0.001 (<0.001) | 0.001 (<0.001) | 0.001 (<0.001) | 0.001 (<0.001) |
| | | mSSL | **0.000** (<0.001) | **0.000** (<0.001) | **0.000** (<0.001) | **0.000** (<0.001) | **0.000** (<0.001) |
| | | $MR^2$ | **0.000** (<0.001) | **0.000** (<0.001) | **0.000** (<0.001) | **0.000** (<0.001) | **0.000** (<0.001) |

**Table S.7. Scenario I-Null: Sum of squared errors (SSE) of the simulated scenario**, averaged over 50 replicates, with standard deviation in brackets, illustrating the performance of different MR multivariable implementations and multi-response multivariable methods to estimate the true simulated effect size. In simulated Scenario I-Null, no direct causal effects are simulated and the outcomes are generated based only on the effect of the confounders. Open parameters are the correlation between exposures $r_X = \{0, 0.2, 0.4, 0.6, 0.8\}$ and the correlation between individual-level responses' errors $r_Y = \{0, 0.2, 0.4, 0.6, 0.8\}$. Best results are highlighted in bold.

| $r_X$ | $\theta_X^U$ | | $\theta_Y^U$ = 1 | $\theta_Y^U$ = 2 |
|---|---|---|---|---|
| 0 | 1 | MV-MR | 0.494 (0.200) | 0.491 (0.203) |
| | | MR-BMA | 0.294 (0.141) | 0.287 (0.140) |
| | | MRCE | 0.368 (0.161) | 0.364 (0.163) |
| | | mSSL | 0.417 (0.644) | 0.333 (0.179) |
| | | $MR^2$ | **0.261** (0.126) | **0.252** (0.120) |
| | 2 | MV-MR | 0.611 (0.246) | 0.616 (0.247) |
| | | MR-BMA | 0.347 (0.163) | 0.341 (0.159) |
| | | MRCE | 0.449 (0.196) | 0.449 (0.199) |
| | | mSSL | 0.577 (0.831) | 0.432 (0.208) |
| | | $MR^2$ | **0.316** (0.143) | **0.311** (0.140) |
| 0.2 | 1 | MV-MR | 0.577 (0.227) | 0.598 (0.261) |
| | | MR-BMA | 0.338 (0.165) | 0.354 (0.200) |
| | | MRCE | 0.416 (0.182) | 0.432 (0.221) |
| | | mSSL | 0.377 (0.181) | 0.398 (0.216) |
| | | $MR^2$ | **0.297** (0.148) | **0.309** (0.177) |
| | 2 | MV-MR | 0.730 (0.322) | 0.743 (0.322) |
| | | MR-BMA | 0.417 (0.220) | 0.425 (0.221) |
| | | MRCE | 0.530 (0.268) | 0.539 (0.270) |
| | | mSSL | 0.646 (0.966) | 0.493 (0.223) |
| | | $MR^2$ | **0.378** (0.199) | **0.386** (0.200) |
| 0.4 | 1 | MV-MR | 0.711 (0.325) | 0.863 (0.325) |
| | | MR-BMA | 0.410 (0.209) | 0.478 (0.211) |
| | | MRCE | 0.522 (0.257) | 0.633 (0.263) |
| | | mSSL | 0.462 (0.252) | 0.589 (0.251) |
| | | $MR^2$ | **0.355** (0.185) | **0.430** (0.189) |
| | 2 | MV-MR | 0.704 (0.385) | 0.875 (0.493) |
| | | MR-BMA | 0.409 (0.235) | 0.478 (0.299) |
| | | MRCE | 0.523 (0.321) | 0.638 (0.402) |
| | | mSSL | 0.461 (0.340) | 0.552 (0.372) |
| | | $MR^2$ | **0.353** (0.216) | **0.432** (0.265) |
| 0.6 | 1 | MV-MR | 0.994 (0.383) | 1.005 (0.597) |
| | | MR-BMA | 0.555 (0.239) | 0.553 (0.373) |
| | | MRCE | 0.730 (0.331) | 0.729 (0.516) |
| | | mSSL | 0.569 (0.277) | 0.564 (1.117) |
| | | $MR^2$ | **0.440** (0.217) | **0.444** (0.340) |
| | 2 | MV-MR | 1.246 (0.490) | 1.261 (0.605) |
| | | MR-BMA | 0.673 (0.302) | 0.677 (0.514) |
| | | MRCE | 0.904 (0.405) | 0.925 (0.367) |
| | | mSSL | 0.881 (0.374) | 0.718 (0.414) |
| | | $MR^2$ | **0.567** (0.269) | **0.577** (0.331) |
| 0.8 | 1 | MV-MR | 2.015 (1.013) | 1.990 (0.984) |
| | | MR-BMA | 1.223 (0.724) | 1.183 (0.664) |
| | | MRCE | 1.330 (0.684) | 1.301 (0.643) |
| | | mSSL | 1.178 (1.093) | 1.022 (0.647) |
| | | $MR^2$ | **0.793** (0.533) | **0.761** (0.455) |
| | 2 | MV-MR | 2.496 (1.192) | 2.519 (1.191) |
| | | MR-BMA | 1.441 (0.807) | 1.469 (0.808) |
| | | MRCE | 1.667 (0.806) | 1.689 (0.788) |
| | | mSSL | 1.475 (1.113) | 1.392 (0.810) |
| | | $MR^2$ | **1.057** (0.629) | **1.077** (0.614) |

**Table S.8. Scenario II-Confounding: Sum of squared errors (SSE) of the simulated scenario**, averaged over 50 replicates, with standard deviation in brackets, illustrating the performance of different MR multivariable implementations and multi-response multivariable methods to estimate the true effect size. In simulated Scenario II-Confounding, 30% of all exposure-confounder combinations are generated to have a direct causal effect. Open parameters are the correlation between exposures $r_X = \{0, 0.2, 0.4, 0.6, 0.8\}$ and confounding effects on the exposures $\theta_X^U = \{1, 2\}$ and outcomes $\theta_Y^U = \{1, 2\}$. Best results are highlighted in bold.

| | | $\theta^A$ | | | | | |
|---|---|---|---|---|---|---|---|
| | | 0.25 | 0.5 | 0.75 | 1 | 1.5 | 2 |
| $r_X$ | 0 | | | | | | |
| | MV-MR | 0.683 (0.254) | 0.944 (0.329) | 1.320 (0.417) | 1.874 (0.565) | 3.559 (1.094) | 5.801 (1.859) |
| | MR-BMA | 0.407 (0.179) | 0.502 (0.210) | 0.648 (0.266) | 0.859 (0.343) | 1.556 (0.596) | 2.662 (1.021) |
| | MRCE | 0.521 (0.209) | 0.607 (0.239) | 0.670 (0.274) | 0.736 (0.297) | 1.003 (0.601) | 1.340 (0.887) |
| | mSSL | 0.465 (0.249) | 0.530 (0.257) | 0.560 (0.277) | 0.605 (0.294) | 0.721 (0.393) | 1.017 (0.788) |
| | $MR^2$ | **0.360** (0.160) | **0.406** (0.159) | **0.431** (0.168) | **0.462** (0.162) | **0.610** (0.499) | **0.652** (0.337) |
| | 0.2 | | | | | | |
| | MV-MR | 0.783 (0.379) | 1.054 (0.442) | 1.514 (0.553) | 2.179 (0.686) | 4.015 (1.321) | 6.445 (2.103) |
| | MR-BMA | 0.428 (0.209) | 0.563 (0.242) | 0.774 (0.288) | 1.065 (0.375) | 1.756 (0.571) | 2.731 (0.841) |
| | MRCE | 0.567 (0.274) | 0.666 (0.316) | 0.786 (0.361) | 0.893 (0.431) | 1.141 (0.649) | 1.425 (0.876) |
| | mSSL | 0.561 (0.482) | 0.630 (0.443) | 0.724 (0.461) | 0.747 (0.462) | 0.895 (0.521) | **1.081** (0.516) |
| | $MR^2$ | **0.370** (0.212) | **0.425** (0.231) | **0.498** (0.253) | **0.540** (0.267) | **0.735** (0.484) | 1.092 (0.965) |
| | 0.4 | | | | | | |
| | MV-MR | 0.936 (0.376) | 1.261 (0.450) | 1.822 (0.578) | 2.673 (0.903) | 5.035 (1.847) | 8.244 (3.092) |
| | MR-BMA | 0.517 (0.240) | 0.648 (0.273) | 0.869 (0.335) | 1.201 (0.450) | 2.204 (0.906) | 3.542 (1.403) |
| | MRCE | 0.689 (0.302) | 0.778 (0.321) | 0.882 (0.353) | 1.032 (0.498) | 1.325 (0.917) | 1.629 (1.073) |
| | mSSL | 0.590 (0.289) | 0.682 (0.302) | 0.702 (0.295) | 0.778 (0.354) | 1.089 (0.872) | 1.349 (0.952) |
| | $MR^2$ | **0.436** (0.237) | **0.491** (0.241) | **0.520** (0.242) | **0.647** (0.621) | **0.844** (0.899) | **1.088** (1.137) |
| | 0.6 | | | | | | |
| | MV-MR | 1.332 (0.642) | 1.826 (0.763) | 2.699 (1.051) | 3.869 (1.502) | 7.017 (2.846) | 11.518 (5.051) |
| | MR-BMA | 0.774 (0.571) | 0.971 (0.719) | 1.330 (0.883) | 1.714 (1.036) | 2.965 (1.483) | 4.487 (1.944) |
| | MRCE | 0.947 (0.551) | 1.094 (0.613) | 1.251 (0.668) | 1.434 (0.877) | 1.806 (1.128) | 2.191 (1.176) |
| | mSSL | 0.877 (0.663) | 0.933 (0.678) | 1.013 (0.720) | 1.104 (0.790) | 1.410 (0.943) | 1.795 (1.149) |
| | $MR^2$ | **0.624** (0.523) | **0.693** (0.614) | **0.759** (0.658) | **0.858** (0.722) | **1.125** (0.987) | **1.565** (1.284) |
| | 0.8 | | | | | | |
| | MV-MR | 2.716 (1.263) | 3.693 (1.544) | 5.272 (2.085) | 7.728 (2.944) | 13.611 (5.260) | 22.417 (8.816) |
| | MR-BMA | 1.617 (0.722) | 1.922 (0.858) | 2.432 (1.045) | 3.065 (1.284) | 4.913 (1.875) | 7.036 (2.735) |
| | MRCE | 1.891 (0.764) | 2.118 (0.858) | 2.380 (1.091) | 2.667 (1.275) | 3.004 (1.203) | 3.456 (1.478) |
| | mSSL | 1.538 (0.795) | 1.929 (1.204) | 2.102 (1.180) | 2.325 (1.316) | 3.309 (2.217) | 3.817 (2.051) |
| | $MR^2$ | **1.198** (0.594) | **1.391** (0.644) | **1.560** (0.759) | **1.731** (0.815) | **2.251** (1.153) | **2.707** (1.310) |

**Table S.9. Simulation III-Undirected pleiotropy: Sum of squared errors (SSE) of the simulated scenario**, averaged over 50 replicates, with standard deviation in brackets, illustrating the performance of different MR multivariable implementations and multi-response multivariable methods to estimate the true effect size. In simulated Scenario III-Undirected pleiotropy, 30% of all exposure-confounder combinations are generated to have a direct causal effect. Open parameters are the correlation between exposures $r_X = \{0, 0.2, 0.4, 0.6, 0.8\}$ and the pleiotropic effect $\theta^A = \{0.25, 0.5, 0.75, 1, 1.5, 2\}$. Confounding effects on the exposures and outcomes are fixed at $\theta_X^U = 2$ and $\theta_Y^U = 1$, respectively. Best results are highlighted in bold.

| | | $\theta^A$ | | | | | |
|---|---|---|---|---|---|---|---|
| | | 0.25 | 0.5 | 0.75 | 1 | 1.5 | 2 |
| $r_X$ 0 | MV-MR | 0.679 (0.255) | 0.909 (0.332) | 1.290 (0.456) | 1.829 (0.687) | 3.383 (1.331) | 5.621 (2.265) |
| | MR-BMA | 0.410 (0.183) | 0.510 (0.214) | 0.658 (0.274) | 0.895 (0.358) | 1.659 (0.603) | 2.642 (0.885) |
| | MRCE | 0.491 (0.193) | 0.534 (0.210) | 0.566 (0.219) | 0.616 (0.23) | 0.756 (0.298) | 0.960 (0.422) |
| | mSSL | 0.442 (0.244) | 0.491 (0.296) | 0.526 (0.271) | 0.564 (0.266) | 0.681 (0.367) | **0.858** (0.420) |
| | MR$^2$ | **0.366** (0.165) | **0.412** (0.166) | **0.453** (0.187) | **0.507** (0.205) | **0.588** (0.252) | 0.938 (1.021) |
| 0.2 | MV-MR | 0.768 (0.363) | 1.021 (0.438) | 1.465 (0.607) | 2.080 (0.857) | 3.767 (1.538) | 6.349 (2.649) |
| | MR-BMA | 0.430 (0.199) | 0.560 (0.236) | 0.786 (0.340) | 1.088 (0.500) | 1.818 (0.799) | 2.849 (1.118) |
| | MRCE | 0.540 (0.272) | 0.584 (0.278) | 0.659 (0.306) | 0.722 (0.326) | 0.893 (0.374) | 1.062 (0.437) |
| | mSSL | 0.484 (0.326) | 0.591 (0.485) | 0.658 (0.440) | 0.696 (0.451) | 0.810 (0.415) | **0.989** (0.513) |
| | MR$^2$ | **0.371** (0.195) | **0.424** (0.221) | **0.496** (0.227) | **0.546** (0.249) | **0.673** (0.359) | 1.033 (0.996) |
| 0.4 | MV-MR | 0.924 (0.393) | 1.245 (0.500) | 1.780 (0.695) | 2.541 (0.989) | 4.782 (1.885) | 8.228 (3.153) |
| | MR-BMA | 0.508 (0.244) | 0.624 (0.260) | 0.819 (0.348) | 1.091 (0.495) | 1.970 (0.899) | 3.301 (1.250) |
| | MRCE | 0.645 (0.296) | 0.699 (0.305) | 0.760 (0.317) | 0.850 (0.359) | 1.050 (0.500) | 1.260 (0.640) |
| | mSSL | 0.546 (0.279) | 0.608 (0.286) | 0.678 (0.285) | 0.709 (0.296) | 0.853 (0.347) | 1.167 (0.695) |
| | MR$^2$ | **0.430** (0.235) | **0.480** (0.243) | **0.526** (0.258) | **0.593** (0.275) | **0.827** (0.765) | **0.994** (0.696) |
| 0.6 | MV-MR | 1.294 (0.618) | 1.764 (0.755) | 2.548 (1.036) | 3.613 (1.409) | 6.802 (2.678) | 11.567 (4.625) |
| | MR-BMA | 0.732 (0.451) | 0.876 (0.481) | 1.156 (0.570) | 1.508 (0.663) | 2.581 (1.055) | 4.122 (1.611) |
| | MRCE | 0.903 (0.516) | 0.975 (0.549) | 1.072 (0.589) | 1.172 (0.618) | 1.406 (0.719) | 1.683 (0.702) |
| | mSSL | 0.835 (0.669) | 0.882 (0.658) | 0.946 (0.676) | 1.050 (0.700) | 1.261 (0.788) | 1.637 (0.883) |
| | MR$^2$ | **0.593** (0.432) | **0.677** (0.518) | **0.748** (0.605) | **0.821** (0.681) | **1.041** (0.855) | **1.420** (1.064) |
| 0.8 | MV-MR | 2.661 (1.249) | 3.515 (1.528) | 5.054 (2.103) | 7.234 (2.824) | 13.513 (5.142) | 22.485 (8.544) |
| | MR-BMA | 1.606 (0.691) | 1.892 (0.819) | 2.381 (0.975) | 3.102 (1.272) | 5.314 (2.453) | 7.718 (3.558) |
| | MRCE | 1.791 (0.732) | 1.892 (0.770) | 2.100 (0.867) | 2.257 (0.940) | 2.685 (1.216) | 3.140 (1.419) |
| | mSSL | 1.393 (0.757) | 1.651 (0.982) | 1.894 (1.155) | 2.103 (1.246) | 2.839 (1.726) | 3.259 (1.913) |
| | MR$^2$ | **1.205** (0.588) | **1.360** (0.604) | **1.514** (0.716) | **1.703** (0.766) | **2.156** (1.014) | **2.689** (1.387) |

**Table S.10. Simulation IV-Directed pleiotropy: Sum of squared errors (SSE) of the simulated scenario**, averaged over 50 replicates, with standard deviation in brackets, illustrating the performance of different MR multivariable implementations and multi-response multivariable methods to estimate the true effect size. In simulated Scenario IV-Directed pleiotropy, 30% of all exposure-confounder combinations are generated to have a direct causal effect. Open parameters are the correlation between exposures $r_X = \{0, 0.2, 0.4, 0.6, 0.8\}$ and the pleiotropic effect $\theta^A = \{0.25, 0.5, 0.75, 1, 1.5, 2\}$. Confounding effects on the exposures and outcomes are fixed at $\theta^U_X = 2$ and $\theta^U_Y = 1$, respectively. Best results are highlighted in bold.

| | | $r_Y$ | | | | |
|---|---|---|---|---|---|---|
| | | 0 | 0.2 | 0.4 | 0.6 | 0.8 |
| $r_X$ | 0 | | | | | |
| | MV-MR | 0.611 (0.246) | 0.663 (0.315) | 0.658 (0.296) | 0.667 (0.305) | 0.653 (0.308) |
| | MR-BMA | 0.347 (0.163) | 0.363 (0.216) | 0.361 (0.207) | 0.372 (0.203) | 0.357 (0.193) |
| | MRCE | 0.449 (0.196) | 0.475 (0.254) | 0.462 (0.227) | 0.463 (0.235) | 0.427 (0.228) |
| | mSSL | 0.577 (0.831) | 0.582 (0.858) | 0.694 (1.053) | 0.664 (1.062) | 0.623 (1.036) |
| | $MR^2$ | **0.315** (0.143) | **0.329** (0.207) | **0.325** (0.216) | **0.305** (0.189) | **0.268** (0.183) |
| | 0.2 | | | | | |
| | MV-MR | 0.730 (0.322) | 0.744 (0.298) | 0.755 (0.279) | 0.745 (0.276) | 0.737 (0.278) |
| | MR-BMA | 0.417 (0.220) | 0.433 (0.219) | 0.440 (0.218) | 0.422 (0.210) | 0.425 (0.217) |
| | MRCE | 0.530 (0.268) | 0.541 (0.252) | 0.540 (0.234) | 0.512 (0.216) | 0.483 (0.230) |
| | mSSL | 0.646 (0.966) | 0.616 (0.964) | 0.638 (0.964) | 0.594 (0.969) | 0.830 (1.773) |
| | $MR^2$ | **0.378** (0.199) | **0.378** (0.197) | **0.385** (0.197) | **0.355** (0.202) | **0.321** (0.201) |
| | 0.4 | | | | | |
| | MV-MR | 0.863 (0.385) | 0.851 (0.348) | 0.847 (0.340) | 0.867 (0.346) | 0.853 (0.343) |
| | MR-BMA | 0.478 (0.235) | 0.455 (0.221) | 0.462 (0.215) | 0.470 (0.224) | 0.484 (0.234) |
| | MRCE | 0.633 (0.321) | 0.594 (0.264) | 0.590 (0.236) | 0.604 (0.258) | 0.566 (0.227) |
| | mSSL | 0.589 (0.340) | 0.574 (0.405) | 0.674 (0.868) | 0.722 (1.014) | 0.810 (1.406) |
| | $MR^2$ | **0.430** (0.216) | **0.401** (0.201) | **0.388** (0.191) | **0.361** (0.161) | **0.334** (0.152) |
| | 0.6 | | | | | |
| | MV-MR | 1.246 (0.597) | 1.205 (0.519) | 1.212 (0.514) | 1.200 (0.464) | 1.195 (0.470) |
| | MR-BMA | 0.673 (0.373) | 0.651 (0.329) | 0.680 (0.369) | 0.700 (0.348) | 0.693 (0.337) |
| | MRCE | 0.904 (0.516) | 0.849 (0.413) | 0.867 (0.431) | 0.849 (0.423) | 0.805 (0.355) |
| | mSSL | 0.881 (1.117) | 0.851 (1.095) | 0.845 (1.168) | 0.891 (1.167) | 1.005 (1.762) |
| | $MR^2$ | **0.567** (0.340) | **0.534** (0.289) | **0.545** (0.329) | **0.521** (0.278) | **0.438** (0.225) |
| | 0.8 | | | | | |
| | MV-MR | 2.496 (1.192) | 2.391 (0.968) | 2.310 (0.942) | 2.324 (0.939) | 2.287 (0.979) |
| | MR-BMA | 1.441 (0.807) | 1.341 (0.587) | 1.278 (0.525) | 1.346 (0.570) | 1.342 (0.635) |
| | MRCE | 1.667 (0.806) | 1.598 (0.630) | 1.474 (0.545) | 1.511 (0.609) | 1.430 (0.642) |
| | mSSL | 1.475 (1.113) | 1.493 (1.566) | 1.415 (1.533) | 1.371 (1.517) | 1.626 (2.440) |
| | $MR^2$ | **1.057** (0.629) | **0.977** (0.475) | **0.867** (0.425) | **0.844** (0.450) | **0.754** (0.425) |

**Table S.11. Simulation V-Dependence: Sum of squared errors (SSE) of the simulated scenario**, averaged over 50 replicates, with standard deviation in brackets, illustrating the performance of different MR multivariable implementations and multi-response multivariable methods to estimate the true effect size. In simulated Scenario V-Dependence, 30% of all exposure-confounder combinations are generated to have a direct causal effect. Open parameters are the correlation between exposures $r_X = \{0, 0.2, 0.4, 0.6, 0.8\}$ and the correlation between individual-level responses' errors $r_Y = \{0, 0.2, 0.4, 0.6, 0.8\}$. Confounding effects on the exposures and outcomes are fixed at $\theta_X^U = 2$ and $\theta_Y^U = 1$, respectively. Best results are highlighted in bold.

##### S.3.3 Supplemental Figures

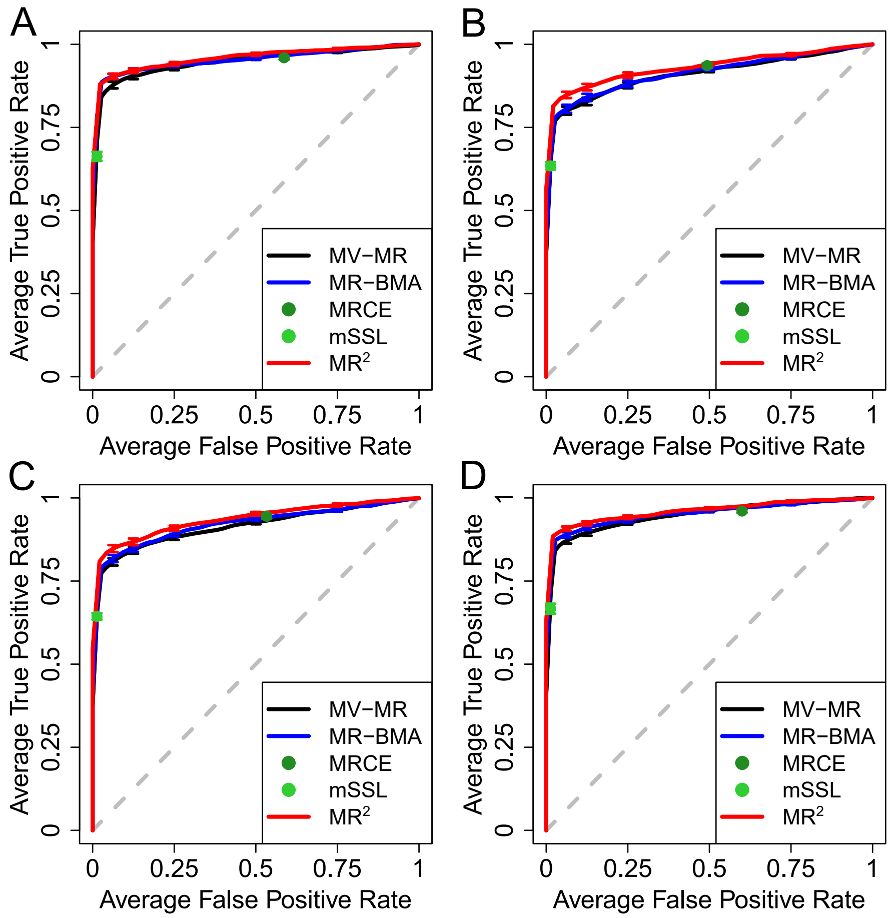


**Figure S.1. Receiver operating characteristic (ROC) curves in different simulated scenarios when there is no correlation between exposures**, averaged over 50 replicates, illustrating the performance of different MR multivariable implementations and multi-response multivariable methods to distinguish between true and false causal exposures for the simulated outcomes by plotting the true (TPR) against the false positive rate (FPR). (**A**) depicts baseline Scenario II-Confounding where only the confounding effects on the exposures and outcomes, $\theta_X^U = 2$ and $\theta_Y^U = 1$, respectively, are used to simulate the data. Residual correlation induced by shared undirected pleiotropy (Scenario III-Undirected pleiotropy) and shared directed pleiotropy (Scenario IV-Directed pleiotropy) with pleiotropic effect set at $\theta^A = 1$ are shown in (**B**) and (**C**), respectively. (**D**) displays the performance of the different methods in Scenario IV-Dependence, where the correlation between individual-level responses' errors is fixed at $r_Y = 0.6$. Vertical bars in each ROC curve, at specific FPR levels, indicate standard error across 50 replicates. For more details on the simulations setting, see Supplemental Table S.2.

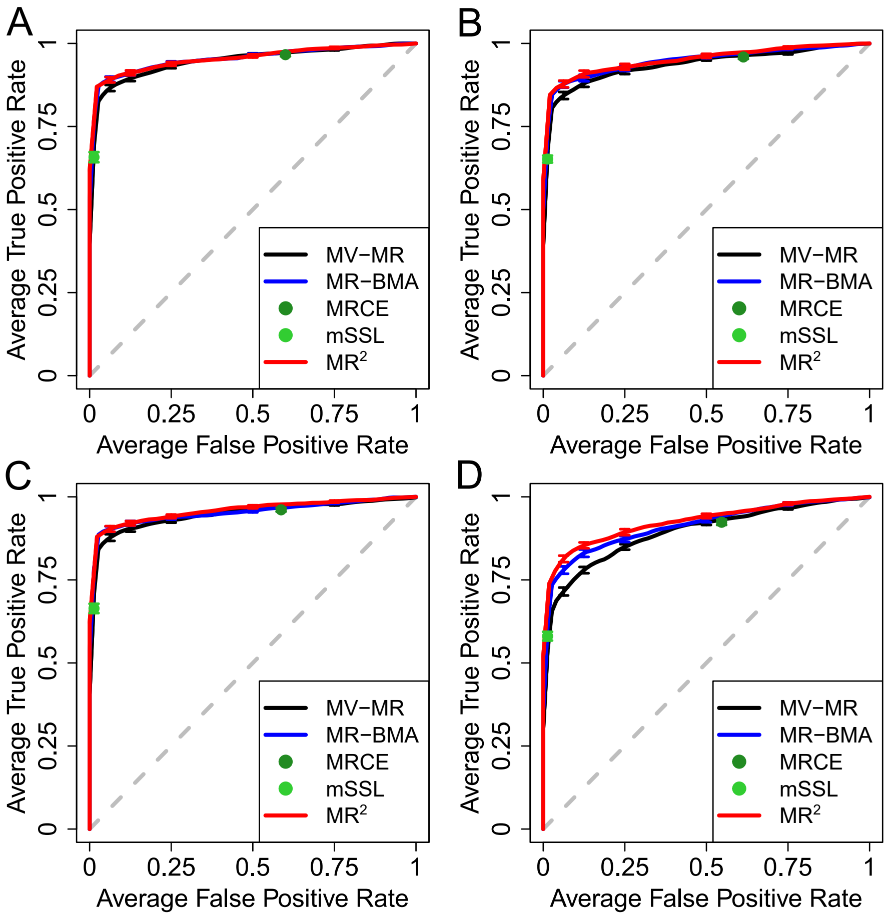


**Figure S.2. Simulation II-Confounding: Receiver operating characteristic (ROC) for different levels of correlation between the exposures**, averaged over 50 replicates, illustrating the performance of different MR multivariable implementations and multi-response multivariable methods to distinguish between true and false causal exposures for the simulated outcomes by plotting the true (TPR) against the false positive rate (FPR). Correlation between the exposures varies from light correlation $r_X = 0.2$ in (**A**) to strong correlation $r_X = 0.8$ in (**D**) with confounding effects on the exposures and outcomes fixed at $\theta_X^U = 2$ and $\theta_Y^U = 1$, respectively. Vertical bars in each ROC curve, at specific FPR levels, indicate standard error across 50 replicates.

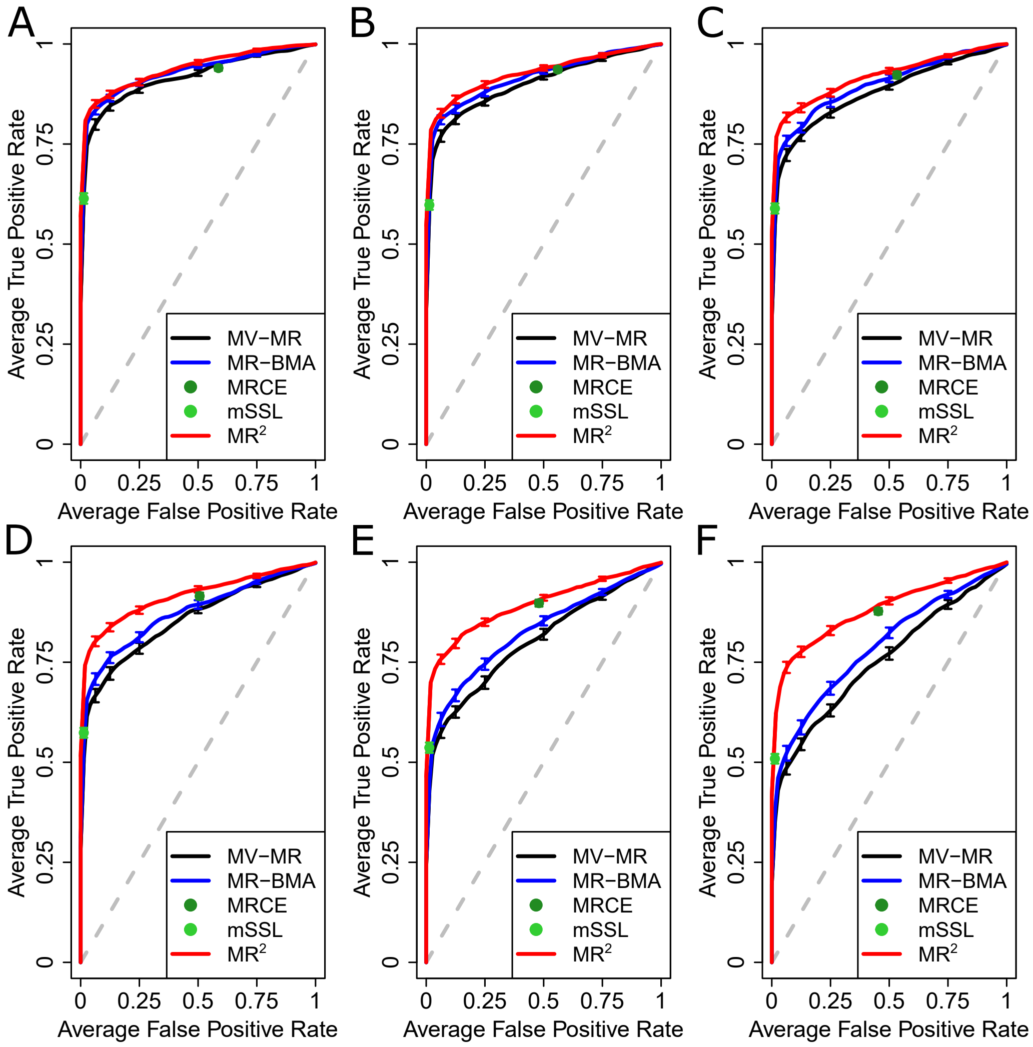


**Figure S.3. Simulation IV-Directed pleiotropy: Receiver operating characteristic (ROC) curves for different levels of the pleiotropic pathway effect $\boldsymbol{\theta^A}$ and when the correlation between exposures is set at $\boldsymbol{r_X = 0.6}$**, averaged over 50 replicates, illustrating the performance of different MR implementations and multi-response statistical methods to distinguish between true and false causal exposures for the simulated outcomes by plotting the true (TPR) against the false positive rate (FPR). Pleiotropic pathway effect varies from (**A**) to (**F**) with values $\theta^A = \{0.25, 0.5, 0.75, 1, 1.5, 2\}$. Confounding effects on the exposures and outcomes are fixed at $\theta_X^U = 2$ and $\theta_Y^U = 1$, respectively. Vertical bars in each ROC curve, at specific FPR levels, indicate standard error across 50 replicates.

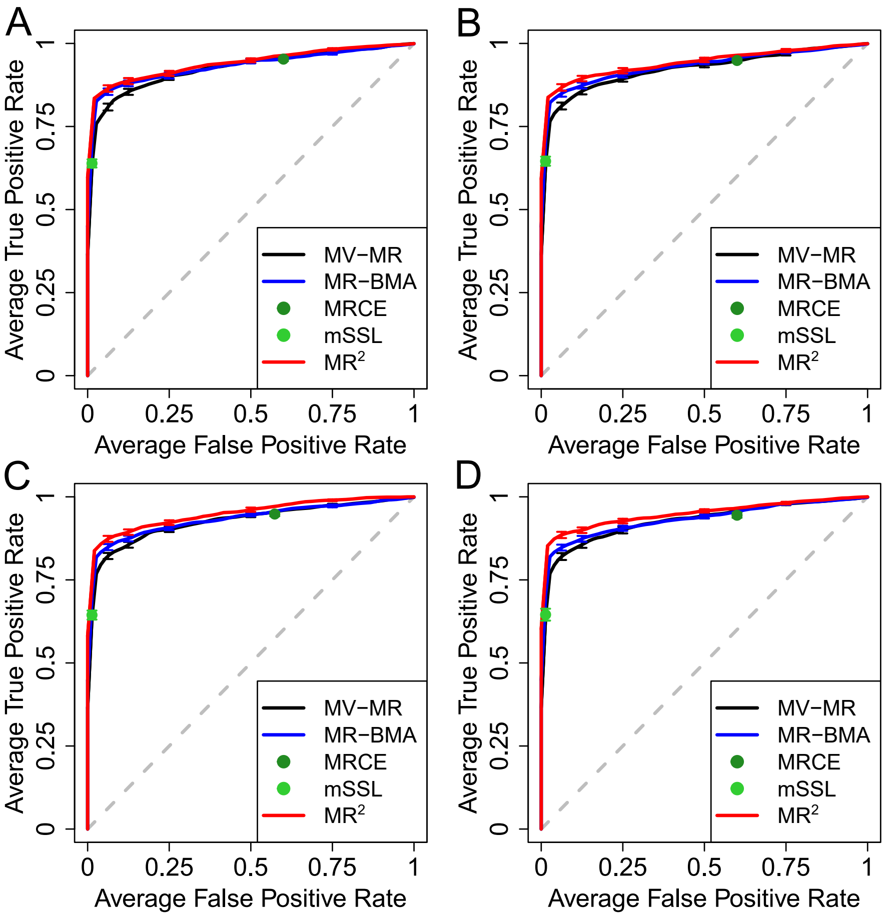


**Figure S.4. Simulation V-Dependence: Receiver operating characteristic (ROC) curves for different levels of the correlation between individual-level responses' errors**, averaged over 50 replicates, illustrating the performance of different MR multivariable implementations and multi-response multivariable methods to distinguish between true and false causal exposures for the simulated outcomes by plotting the true (TPR) against the false positive rate (FPR). Individual-level correlation between the responses' errors varies from (**A**) to (**D**) with values $r_Y = \{0.2, 0.4, 0.6, 0.8\}$. Correlation between exposures is set at $r_X = 0.6$ and confounding effects on the exposures and outcomes are fixed at $\theta_X^U = 2$ and $\theta_Y^U = 1$, respectively. Vertical bars in each ROC curve, at specific FPR levels, indicate standard error across 50 replicates.

### S.4 Application examples

#### S.4.1 Data pre-processing

Application examples are based on publicly available summary-level data. An overview of the data sets, with sample sizes and references, is provided in Supplemental Table S.12.

The first application example considers ten common risk factors for five cardiovascular disease outcomes (CVDs). The first step of the data-processing merges the summary-level data (beta regression coefficients, their standard errors and associated $p$-values) of all exposures by their unique "rs" identifier and aligns the effect direction of the genetic associations with each exposure according to the same effect allele. As IVs, we select any genetic variant which is associated with any of the exposures at genome-wide significance ($p$-value $< 5 \times 10^{-8}$). Next, we merge the genetic variants selected as IVs with the outcome data of the five CVDs by their unique "rs" identifier and aligned the effect direction of the genetic associations with each outcome according to the same effect allele. Finally, we clump the genetic variants to be independent at $r^2 < 0.001$ using a European reference panel [12]. This results in $n = 1,540$ independent genetic variants selected as IVs.

The second application example focuses on ten ApoB-containing lipoprotein subfractions derived from metabolic GWAS using the Nightingale nuclear magnetic resonance spectroscopy platform derived from fasting samples of $24,925$ European descent participants [13]. Instruments selection is based on an external data set. Specifically, given the prior hypothesis of ApoB as the leading exposure for CVDs, we select genetic variants associated with ApoB in UK Biobank at genome-wide significance ($p$-value $< 5 \times 10^{-8}$). Exposures and outcomes summary-level data are merged by their unique "rs" identifier and effect alleles are aligned. Finally, we clump the genetic variants to be independent at $r^2 < 0.001$ using a European reference panel [14]. This resulted in $n = 148$ IVs.

In both application examples, IVW is performed before the analysis for all outcomes and risk factors using weights derived jointly from all responses as described in Appendix. Moreover, the summary-level genetic associations with the risk factors are standardised before the analysis. This is usually required in (multi-response) multivariable regression models since the summary-level genetic associations with the risk factors should be on the same scale in order to interpret and compare the estimated effect sizes.

##### S.4.2 Analysis plan and comparisons with alternative methods

###### $MR^2$ parameters setting

In the first application example, the $MR^2$ algorithm is run for $75,000$ sweeps of which $25,000$ as burn-in and results saved every 50 sweeps, resulting in $1,000$ posterior samples for all the unknowns. In the second application example, we run the algorithm for $150,000$ sweeps of which $50,000$ as burn-in with $1,000$ posterior samples saved.

In both application examples, the hyper-parameters $a_k$ and $b_k$ in eq. (15) are chosen by specifying $\mathbb{E}(\gamma_k) = 2$ and $\mathbb{V}(\gamma_k) = 2$ which imply *a priori* a range between zero and eight for the number of significant direct causal effects for each outcome. We set $v = 1$ in eq. (13) after standardising the summary-level genetic associations with the IVW transformed risk factors in order to place appropriate prior mass on reasonable values of the non-zero direct causal effects [8].

We use Congdon [15] definition of outliers or high-leverage and influential observations and report the number of standardised CPOs less than 0.01 to check model adequacy. After excluding genetic variants with standardised CPOs below this threshold, we refit the model without repeating the procedure even if new outliers or high-leverage and influential observations are detected, see Supplemental Figures S.6 and S.9.

Finally, the FDR level is set at 5% to select important mPPIs and ePPIs, see Supplemental Figures S.7A and B and S.10A and B.

###### MR alternative methods and parameters setting

MR-BMA is run for each outcome separately with a prior probability of inclusion of $2/10 = 0.2$ in application example 1, using ten common risk factors for CVDs, as well as in application example 2, using ten molecular risk factors for the same disease outcomes, in order to match the sparsity prior used in $MR^2$. An exhaustive search evaluating all combinations of exposures in both applications is conducted and no stochastic search is necessary. Because summary-level genetic associations with the risk factors are standardised prior to the analysis, the prior variance on the direct causal effects is set at 1.

All results presented are generated after removing any influential or outlying genetic variant used as IV. Influential genetic variants are detected using the Cook's distance [12] and a cut-off is chosen according to the median of the $F$-distribution with $d$ and $n - d$ degrees of freedom, where $n$ is the number of IVs and $d$ is the size of the best visited model in the exhaustive search.

Outlying genetic variants are identified using the $q$-heterogeneity statistic [12] and threshold using a $\chi^2$ distribution with one degree of freedom and a Bonferroni-adjusted multiple testing correction. Empirical $p$-values for the marginal inclusion probabilities are calculated using a permutation procedure with $100,000$ permutations [14]. Multiple testing adjustment is performed using Benjamini-Hochberg [16] FDR and Bonferroni correction.

MV-MR [9] is applied on the summary-level data after removing outliers or influential variants and multiple testing adjustment is performed using Benjamini-Hochberg FDR and Bonferroni correction.

##### S.4.3 Supplemental Tables

| | Response or risk factor | Acronym | Reference | Number cases & controls |
|---|---|---|---|---|
| Cardiovascular disease outcomes: | Atrial fibrillation | AF | Nielsen *et al.* 2018 [17] | 60,620 cases & 970,216 controls |
| | Cardioembolic stroke | CES | Mishra *et al.* 2022 [18] | 10,804 cases & 1,234,808 controls |
| | Coronary artery disease | CAD | Nelson *et al.* 2017 [19] | 113,937 cases & 339,658 controls |
| | Peripheral artery disease | PAD | Klarin *et al.* 2019 [20] | 31,307 cases & 211,753 controls |
| | Heart failure | HF | Shah *et al.* 2020 [21] | 47,309 cases & 930,014 controls |
| Exposures in application example 1: Common risk factors | Apolipoprotein A | ApoA | Nealelab 2018 [22] | |
| | Apolipoprotein B | ApoB | | |
| | Body mass index | BMI | | |
| | High-density lipoprotein | HDL | | |
| | Low-density lipoprotein | LDL | | |
| | Systolic blood pressure | SBP | | |
| | Triglycerides | TG | | |
| | Physical activity | PA | Wang *et al.* 2022 [23] | 636,924 cases |
| | Smoking | Smoking | Wootton *et al.* 2020 [24] | 462,690 cases |
| | Type 2 diabetes | T2D | Mahajan *et al.* 2018 [25] | 74,124 cases & 824,006 controls |
| Exposures in application example 2: Molecular risk factors | Medium large-density lipoprotein particles | M.LDL.P | Kettunen *et al.* 2016 [13] | 24,925 controls |
| | Large large-density lipoprotein particles | L.LDL.P | | |
| | Intermediate-density lipoprotein particles | IDL.P | | |
| | Extra small very-large density lipoprotein particles | XS.VLDL.P | | |
| | Small very-large density lipoprotein particles | S.VLDL.P | | |
| | Medium very-large density lipoprotein particles | M.VLDL.P | | |
| | Large very-large density lipoprotein particles | L.VLDL.P | | |
| | Extra large very-large density lipoprotein particles | XL.VLDL.P | | |
| | Extra-extra large very-large density lipoprotein particles | XXL.VLDL.P | | |

**Table S.12. Overview of summary-level data in the two application examples.** Common risk factors (application example 1) and molecular risk factors (application example 2) for cardiovascular disease outcomes.

| Outcome | Ranking | Exposure | Effect estimate | Standard error | $p$-value | FDR | Bonferroni |
|---|---|---|---|---|---|---|---|
| CAD | 1 | **SBP** | 0.621 | 0.041 | $1.45\times10^{-47}$ | $1.45\times10^{-46}$ | $1.45\times10^{-46}$ |
| | 2 | **T2D** | 0.125 | 0.013 | $1.69\times10^{-20}$ | $8.43\times10^{-20}$ | $1.69\times10^{-19}$ |
| | 3 | **SMOKING** | 0.507 | 0.093 | $5.98\times10^{-8}$ | $1.99\times10^{-7}$ | $5.98\times10^{-7}$ |
| | 4 | **BMI** | 0.094 | 0.034 | 0.005 | 0.013 | 0.053 |
| | 5 | TG | 0.083 | 0.038 | 0.029 | 0.058 | 0.292 |
| | 6 | LDL | 0.251 | 0.136 | 0.065 | 0.105 | 0.650 |
| | 7 | ApoA | -0.159 | 0.089 | 0.074 | 0.105 | 0.737 |
| | 8 | ApoB | 0.178 | 0.121 | 0.141 | 0.176 | 1.407 |
| | 9 | HDL | 0.084 | 0.094 | 0.372 | 0.414 | 3.723 |
| | 10 | PA | 0.001 | 0.050 | 0.978 | 0.978 | 9.779 |
| PAD | 1 | **T2D** | 0.161 | 0.014 | $7.91\times10^{-30}$ | $7.91\times10^{-29}$ | $7.91\times10^{-29}$ |
| | 2 | **SMOKING** | 0.789 | 0.098 | $1.30\times10^{-15}$ | $6.48\times10^{-15}$ | $1.30\times10^{-14}$ |
| | 3 | **SBP** | 0.326 | 0.043 | $7.81\times10^{-14}$ | $2.60\times10^{-13}$ | $7.81\times10^{-13}$ |
| | 4 | **BMI** | 0.155 | 0.035 | $1.16\times10^{-5}$ | $2.91\times10^{-5}$ | $1.16\times10^{-4}$ |
| | 5 | TG | 0.074 | 0.040 | 0.063 | 0.126 | 0.632 |
| | 6 | ApoA | -0.130 | 0.093 | 0.163 | 0.271 | 1.626 |
| | 7 | ApoB | 0.123 | 0.127 | 0.334 | 0.477 | 3.340 |
| | 8 | HDL | 0.082 | 0.099 | 0.411 | 0.513 | 4.106 |
| | 9 | LDL | 0.035 | 0.143 | 0.807 | 0.818 | 8.067 |
| | 10 | PA | 0.012 | 0.053 | 0.818 | 0.818 | 8.182 |
| HF | 1 | **BMI** | 0.318 | 0.030 | $1.54\times10^{-25}$ | $1.54\times10^{-24}$ | $1.54\times10^{-24}$ |
| | 2 | **SBP** | 0.322 | 0.036 | $2.80\times10^{-18}$ | $1.40\times10^{-17}$ | $2.80\times10^{-17}$ |
| | 3 | **T2D** | 0.047 | 0.012 | $5.63\times10^{-5}$ | $1.88\times10^{-4}$ | $5.63\times10^{-4}$ |
| | 4 | **SMOKING** | 0.323 | 0.083 | $9.77\times10^{-5}$ | $2.44\times10^{-4}$ | $9.77\times10^{-4}$ |
| | 5 | ApoB | 0.113 | 0.108 | 0.296 | 0.584 | 2.958 |
| | 6 | HDL | -0.072 | 0.084 | 0.390 | 0.584 | 3.904 |
| | 7 | PA | 0.037 | 0.045 | 0.409 | 0.584 | 4.088 |
| | 8 | ApoA | 0.048 | 0.079 | 0.540 | 0.674 | 5.396 |
| | 9 | LDL | 0.015 | 0.121 | 0.903 | 0.983 | 9.026 |
| | 10 | TG | -0.001 | 0.034 | 0.983 | 0.983 | 9.829 |
| AF | 1 | **SBP** | 0.286 | 0.036 | $7.71\times10^{-15}$ | $7.71\times10^{-14}$ | $7.71\times10^{-14}$ |
| | 2 | **BMI** | 0.225 | 0.030 | $5.83\times10^{-14}$ | $2.92\times10^{-13}$ | $5.83\times10^{-13}$ |
| | 3 | **TG** | -0.129 | 0.034 | $1.39\times10^{-4}$ | $4.62\times10^{-4}$ | $1.39\times10^{-03}$ |
| | 4 | SMOKING | 0.140 | 0.082 | 0.090 | 0.187 | 0.899 |
| | 5 | ApoB | 0.180 | 0.107 | 0.094 | 0.187 | 0.938 |
| | 6 | HDL | -0.133 | 0.084 | 0.112 | 0.187 | 1.123 |
| | 7 | ApoA | 0.108 | 0.079 | 0.172 | 0.245 | 1.717 |
| | 8 | LDL | -0.151 | 0.120 | 0.210 | 0.263 | 2.103 |
| | 9 | T2D | -0.009 | 0.012 | 0.427 | 0.439 | 4.271 |
| | 10 | PA | 0.035 | 0.045 | 0.439 | 0.439 | 4.392 |
| CES | 1 | **SBP** | 0.266 | 0.063 | $2.76\times10^{-5}$ | $2.76\times10^{-4}$ | $2.76\times10^{-4}$ |
| | 2 | **T2D** | 0.064 | 0.020 | 0.001 | 0.007 | 0.015 |
| | 3 | **SMOKING** | 0.345 | 0.141 | 0.014 | 0.048 | 0.143 |
| | 4 | TG | -0.117 | 0.058 | 0.044 | 0.111 | 0.444 |
| | 5 | PA | 0.088 | 0.077 | 0.252 | 0.467 | 2.517 |
| | 6 | ApoB | 0.199 | 0.184 | 0.280 | 0.467 | 2.803 |
| | 7 | LDL | -0.177 | 0.208 | 0.395 | 0.564 | 3.948 |
| | 8 | HDL | -0.056 | 0.144 | 0.698 | 0.867 | 6.976 |
| | 9 | ApoA | 0.038 | 0.135 | 0.780 | 0.867 | 7.804 |
| | 10 | BMI | -0.001 | 0.051 | 0.979 | 0.979 | 9.792 |

**Table S.13. Application example 1 on common risk factors for cardiovascular disease outcomes using standard multivariable MR (MV-MR)** which is run on each outcome separately. Results include the causal effect estimates, the standard error and the corresponding $p$-value. We adjust for multiple testing using Benjamini-Hochberg False Discovery Rate (FDR) and Bonferroni correction. For each outcome, we sort the exposures by their $p$-value in increasing order. Exposures selected at 5% FDR highlighted in bold.

| Outcome | Ranking | Exposure | Effect estimate | Standard error | $p$-value | FDR | Bonferroni |
|---|---|---|---|---|---|---|---|
| CAD | | (Intercept) | -0.024 | 0.029 | 0.401 | 0.401 | - |
| | 1 | **SBP** | 0.621 | 0.041 | $1.53\times10^{-47}$ | $1.53\times10^{-46}$ | $1.53\times10^{-46}$ |
| | 2 | **T2D** | 0.125 | 0.013 | $1.46\times10^{-20}$ | $7.31\times10^{-20}$ | $1.46\times10^{-19}$ |
| | 3 | **SMOKING** | 0.506 | 0.093 | $6.37\times10^{-8}$ | $2.12\times10^{-7}$ | $6.37\times10^{-7}$ |
| | 4 | **BMI** | 0.094 | 0.034 | 0.005 | 0.013 | 0.051 |
| | 5 | TG | 0.083 | 0.038 | 0.030 | 0.059 | 0.296 |
| | 6 | LDL | 0.253 | 0.136 | 0.064 | 0.105 | 0.636 |
| | 7 | ApoA | -0.159 | 0.089 | 0.073 | 0.105 | 0.732 |
| | 8 | ApoB | 0.177 | 0.121 | 0.144 | 0.180 | 1.000 |
| | 9 | HDL | 0.085 | 0.094 | 0.371 | 0.412 | 1.000 |
| | 10 | PA | -0.001 | 0.050 | 0.985 | 0.985 | 1.000 |
| PAD | | (Intercept) | 0.004 | 0.030 | 0.883 | 0.883 | - |
| | 1 | **T2D** | 0.161 | 0.014 | $8.79\times10^{-30}$ | $8.79\times10^{-29}$ | $8.79\times10^{-29}$ |
| | 2 | **SMOKING** | 0.790 | 0.098 | $1.31\times10^{-15}$ | $6.55\times10^{-15}$ | $1.31\times10^{-14}$ |
| | 3 | **SBP** | 0.326 | 0.043 | $7.92\times10^{-14}$ | $2.64\times10^{-13}$ | $7.92\times10^{-13}$ |
| | 4 | **BMI** | 0.155 | 0.035 | $1.19\times10^{-5}$ | $2.97\times10^{-5}$ | $1.19\times10^{-4}$ |
| | 5 | TG | 0.074 | 0.040 | 0.063 | 0.126 | 0.631 |
| | 6 | ApoA | -0.130 | 0.093 | 0.163 | 0.271 | 1.000 |
| | 7 | ApoB | 0.123 | 0.127 | 0.333 | 0.476 | 1.000 |
| | 8 | HDL | 0.082 | 0.099 | 0.411 | 0.513 | 1.000 |
| | 9 | LDL | 0.035 | 0.143 | 0.808 | 0.812 | 1.000 |
| | 10 | PA | 0.013 | 0.053 | 0.812 | 0.812 | 1.000 |
| HF | | (Intercept) | 0.000 | 0.025 | 0.990 | 0.990 | - |
| | 1 | **BMI** | 0.318 | 0.030 | $1.63\times10^{-25}$ | $1.63\times10^{-24}$ | $1.63\times10^{-24}$ |
| | 2 | **SBP** | 0.322 | 0.036 | $2.87\times10^{-18}$ | $1.44\times10^{-17}$ | $2.87\times10^{-17}$ |
| | 3 | **T2D** | 0.047 | 0.012 | $5.68\times10^{-5}$ | $1.89\times10^{-4}$ | $5.68\times10^{-4}$ |
| | 4 | **SMOKING** | 0.323 | 0.083 | $9.85\times10^{-5}$ | $2.46\times10^{-4}$ | 0.001 |
| | 5 | ApoB | 0.113 | 0.108 | 0.296 | 0.586 | 1.000 |
| | 6 | HDL | -0.072 | 0.084 | 0.391 | 0.586 | 1.000 |
| | 7 | PA | 0.037 | 0.045 | 0.410 | 0.586 | 1.000 |
| | 8 | ApoA | 0.048 | 0.079 | 0.540 | 0.675 | 1.000 |
| | 9 | LDL | 0.015 | 0.121 | 0.903 | 0.983 | 1.000 |
| | 10 | TG | -0.001 | 0.034 | 0.983 | 0.983 | 1.000 |
| AF | | (Intercept) | 0.020 | 0.025 | 0.436 | 0.436 | - |
| | 1 | **SBP** | 0.286 | 0.036 | $7.62\times10^{-15}$ | $7.62\times10^{-14}$ | $7.62\times10^{-14}$ |
| | 2 | **BMI** | 0.225 | 0.030 | $6.65\times10^{-14}$ | $3.33\times10^{-13}$ | $6.65\times10^{-13}$ |
| | 3 | **TG** | -0.129 | 0.034 | $1.43\times10^{-4}$ | $4.76\times10^{-4}$ | 0.001 |
| | 4 | SMOKING | 0.141 | 0.082 | 0.088 | 0.184 | 0.877 |
| | 5 | ApoB | 0.181 | 0.107 | 0.092 | 0.184 | 0.920 |
| | 6 | HDL | -0.133 | 0.084 | 0.112 | 0.187 | 1.000 |
| | 7 | ApoA | 0.108 | 0.079 | 0.171 | 0.245 | 1.000 |
| | 8 | LDL | -0.152 | 0.120 | 0.207 | 0.259 | 1.000 |
| | 9 | T2D | -0.009 | 0.012 | 0.417 | 0.417 | 1.000 |
| | 10 | PA | 0.036 | 0.045 | 0.417 | 0.417 | 1.000 |
| CES | | **(Intercept)** | 0.169 | 0.043 | $9.58\times10^{-5}$ | $9.58\times10^{-5}$ | - |
| | 1 | **SBP** | 0.265 | 0.063 | $2.56\times10^{-5}$ | $2.56\times10^{-4}$ | $2.56\times10^{-4}$ |
| | 2 | **T2D** | 0.062 | 0.020 | 0.002 | 0.009 | 0.019 |
| | 3 | **SMOKING** | 0.357 | 0.140 | 0.011 | 0.037 | 0.110 |
| | 4 | TG | -0.116 | 0.058 | 0.044 | 0.109 | 0.438 |
| | 5 | PA | 0.106 | 0.077 | 0.168 | 0.336 | 1.000 |
| | 6 | ApoB | 0.204 | 0.183 | 0.265 | 0.441 | 1.000 |
| | 7 | LDL | -0.181 | 0.207 | 0.384 | 0.548 | 1.000 |
| | 8 | HDL | -0.060 | 0.143 | 0.675 | 0.844 | 1.000 |
| | 9 | ApoA | 0.041 | 0.134 | 0.760 | 0.844 | 1.000 |
| | 10 | BMI | -0.004 | 0.051 | 0.936 | 0.936 | 1.000 |

**Table S.14. Application example 1 on common risk factors for cardiovascular disease outcomes using multivariable MR-Egger** which is run on each outcome separately. Results include the causal effect estimates as well as the intercept, the standard error and the corresponding $p$-value. We adjust for multiple testing using Benjamini-Hochberg False Discovery Rate (FDR) and Bonferroni correction. For each outcome, we sort the exposures by their $p$-value in increasing order. Exposures selected at 5% FDR highlighted in bold.

| Outcome | Ranking | Exposure | MACE | mPPI | Empirical $p$-value | FDR | Bonferroni |
|---|---|---|---|---|---|---|---|
| CAD | 1 | **SBP** | 0.626 | 1.000 | $1.00\times10^{-5}$ | $3.33\times10^{-5}$ | $1.00\times10^{-4}$ |
| | 2 | **SMOKING** | 0.560 | 1.000 | $1.00\times10^{-5}$ | $3.33\times10^{-5}$ | $1.00\times10^{-4}$ |
| | 3 | **T2D** | 0.137 | 1.000 | $1.00\times10^{-5}$ | $3.33\times10^{-5}$ | $1.00\times10^{-4}$ |
| | 4 | **ApoB** | 0.285 | 0.659 | 0.001 | 0.002 | 0.007 |
| | 5 | **HDL** | -0.034 | 0.295 | 0.017 | 0.032 | 0.171 |
| | 6 | **LDL** | 0.161 | 0.344 | 0.022 | 0.032 | 0.219 |
| | 7 | **BMI** | 0.047 | 0.409 | 0.022 | 0.032 | 0.221 |
| | 8 | ApoA | -0.020 | 0.185 | 0.062 | 0.078 | 0.624 |
| | 9 | TG | 0.002 | 0.033 | 0.902 | 1.000 | 1.000 |
| | 10 | PA | 0.000 | 0.002 | 1.000 | 1.000 | 1.000 |
| PAD | 1 | **SMOKING** | 0.787 | 1.000 | $1.00\times10^{-5}$ | $2.00\times10^{-5}$ | $1.00\times10^{-4}$ |
| | 2 | **SBP** | 0.325 | 1.000 | $1.00\times10^{-5}$ | $2.00\times10^{-5}$ | $1.00\times10^{-4}$ |
| | 3 | **ApoB** | 0.184 | 0.975 | $1.00\times10^{-5}$ | $2.00\times10^{-5}$ | $1.00\times10^{-4}$ |
| | 4 | **BMI** | 0.174 | 0.998 | $1.00\times10^{-5}$ | $2.00\times10^{-5}$ | $1.00\times10^{-4}$ |
| | 5 | **T2D** | 0.171 | 1.000 | $1.00\times10^{-5}$ | $2.00\times10^{-5}$ | $1.00\times10^{-4}$ |
| | 6 | LDL | 0.005 | 0.025 | 0.978 | 1.000 | 1.000 |
| | 7 | TG | 0.000 | 0.000 | 1.000 | 1.000 | 1.000 |
| | 8 | PA | 0.000 | 0.000 | 1.000 | 1.000 | 1.000 |
| | 9 | ApoA | 0.000 | 0.001 | 1.000 | 1.000 | 1.000 |
| | 10 | HDL | 0.000 | 0.001 | 1.000 | 1.000 | 1.000 |
| HF | 1 | **SBP** | 0.324 | 1.000 | $1.00\times10^{-5}$ | $5.00\times10^{-5}$ | $1.00\times10^{-4}$ |
| | 2 | **BMI** | 0.324 | 1.000 | $1.00\times10^{-5}$ | $5.00\times10^{-5}$ | $1.00\times10^{-4}$ |
| | 3 | **ApoB** | 0.117 | 0.897 | $2.00\times10^{-5}$ | $6.67\times10^{-5}$ | $2.00\times10^{-4}$ |
| | 4 | **T2D** | 0.052 | 0.993 | $4.00\times10^{-5}$ | $1.00\times10^{-4}$ | $4.00\times10^{-4}$ |
| | 5 | **SMOKING** | 0.323 | 0.991 | $2.90\times10^{-4}$ | 0.001 | 0.003 |
| | 6 | LDL | 0.014 | 0.104 | 0.267 | 0.445 | 1.000 |
| | 7 | TG | 0.000 | 0.001 | 1.000 | 1.000 | 1.000 |
| | 8 | PA | 0.000 | 0.000 | 1.000 | 1.000 | 1.000 |
| | 9 | ApoA | 0.000 | 0.000 | 1.000 | 1.000 | 1.000 |
| | 10 | HDL | 0.000 | 0.001 | 1.000 | 1.000 | 1.000 |
| AF | 1 | **SBP** | 0.281 | 1.000 | $1.00\times10^{-5}$ | $5.00\times10^{-5}$ | $1.00\times10^{-4}$ |
| | 2 | **BMI** | 0.233 | 1.000 | $1.00\times10^{-5}$ | $5.00\times10^{-5}$ | $1.00\times10^{-4}$ |
| | 3 | **TG** | -0.035 | 0.520 | 0.009 | 0.031 | 0.094 |
| | 4 | HDL | -0.002 | 0.048 | 0.700 | 0.995 | 1.000 |
| | 5 | SMOKING | 0.018 | 0.132 | 0.870 | 0.995 | 1.000 |
| | 6 | ApoB | 0.003 | 0.040 | 0.878 | 0.995 | 1.000 |
| | 7 | ApoA | -0.001 | 0.030 | 0.923 | 0.995 | 1.000 |
| | 8 | LDL | -0.002 | 0.027 | 0.968 | 0.995 | 1.000 |
| | 9 | T2D | 0.000 | 0.008 | 0.988 | 0.995 | 1.000 |
| | 10 | PA | 0.001 | 0.028 | 0.995 | 0.995 | 1.000 |
| CES | 1 | **SBP** | 0.295 | 1.000 | $1.00\times10^{-5}$ | $5.00\times10^{-5}$ | $1.00\times10^{-4}$ |
| | 2 | **BMI** | 0.217 | 1.000 | $1.00\times10^{-5}$ | $5.00\times10^{-5}$ | $1.00\times10^{-4}$ |
| | 3 | TG | -0.013 | 0.232 | 0.048 | 0.160 | 0.481 |
| | 4 | SMOKING | 0.041 | 0.244 | 0.381 | 0.953 | 1.000 |
| | 5 | HDL | -0.001 | 0.025 | 0.947 | 0.990 | 1.000 |
| | 6 | ApoA | -0.001 | 0.024 | 0.960 | 0.990 | 1.000 |
| | 7 | ApoB | 0.002 | 0.024 | 0.970 | 0.990 | 1.000 |
| | 8 | PA | 0.002 | 0.040 | 0.978 | 0.990 | 1.000 |
| | 9 | T2D | 0.000 | 0.010 | 0.979 | 0.990 | 1.000 |
| | 10 | LDL | -0.001 | 0.020 | 0.990 | 0.990 | 1.000 |

**Table S.15. Application example 1 on common risk factors for cardiovascular disease outcomes using the Mendelian randomisation Bayesian model averaging (MR-BMA) algorithm** which is run on each outcome separately. Results include the model-averaged causal effect estimate (MACE), the marginal posterior probability of inclusion (mPPI) and the corresponding empirical $p$-value [12]. We adjust for multiple testing using Benjamini-Hochberg False Discovery Rate (FDR) and Bonferroni correction. For each outcome, we sort the exposures by their empirical $p$-value in increasing order and the absolute value of MACE in decreasing order. Exposures selected at 5% FDR highlighted in bold.

| Outcome | Ranking | Exposure | Effect estimate | Standard error | $p$-value | FDR | Bonferroni |
|---|---|---|---|---|---|---|---|
| CAD | 1 | M.VLDL.P | 0.354 | 0.289 | 0.224 | 0.853 | 1.000 |
| | 2 | S.LDL.P | -0.519 | 0.549 | 0.347 | 0.853 | 1.000 |
| | 3 | M.LDL.P | 1.550 | 1.727 | 0.371 | 0.853 | 1.000 |
| | 4 | S.VLDL.P | -0.239 | 0.340 | 0.483 | 0.853 | 1.000 |
| | 5 | L.LDL.P | -1.096 | 2.207 | 0.620 | 0.853 | 1.000 |
| | 6 | IDL.P | 0.541 | 1.329 | 0.685 | 0.853 | 1.000 |
| | 7 | L.VLDL.P | -0.120 | 0.322 | 0.709 | 0.853 | 1.000 |
| | 8 | XL.VLDL.P | -0.068 | 0.212 | 0.749 | 0.853 | 1.000 |
| | 9 | XXL.VLDL.P | -0.048 | 0.184 | 0.795 | 0.853 | 1.000 |
| | 10 | XS.VLDL.P | 0.105 | 0.566 | 0.853 | 0.853 | 1.000 |
| PAD | 1 | XS.VLDL.P | 0.727 | 0.469 | 0.124 | 0.632 | 1.000 |
| | 2 | S.LDL.P | -0.690 | 0.533 | 0.198 | 0.632 | 1.000 |
| | 3 | S.VLDL.P | -0.384 | 0.305 | 0.210 | 0.632 | 1.000 |
| | 4 | XL.VLDL.P | -0.240 | 0.209 | 0.253 | 0.632 | 1.000 |
| | 5 | XXL.VLDL.P | 0.172 | 0.184 | 0.352 | 0.703 | 1.000 |
| | 6 | M.LDL.P | 1.164 | 1.666 | 0.486 | 0.810 | 1.000 |
| | 7 | IDL.P | -0.636 | 1.156 | 0.583 | 0.812 | 1.000 |
| | 8 | M.VLDL.P | 0.126 | 0.276 | 0.649 | 0.812 | 1.000 |
| | 9 | L.VLDL.P | 0.078 | 0.309 | 0.801 | 0.890 | 1.000 |
| | 10 | L.LDL.P | -0.076 | 2.065 | 0.971 | 0.971 | 1.000 |
| HF | 1 | M.LDL.P | 3.620 | 1.478 | 0.016 | 0.156 | 0.156 |
| | 2 | L.LDL.P | -3.783 | 1.856 | 0.044 | 0.158 | 0.435 |
| | 3 | S.VLDL.P | -0.550 | 0.275 | 0.047 | 0.158 | 0.474 |
| | 4 | S.LDL.P | -0.867 | 0.468 | 0.066 | 0.166 | 0.664 |
| | 5 | M.VLDL.P | 0.390 | 0.249 | 0.120 | 0.240 | 1.000 |
| | 6 | XXL.VLDL.P | 0.218 | 0.165 | 0.189 | 0.287 | 1.000 |
| | 7 | IDL.P | 1.344 | 1.046 | 0.201 | 0.287 | 1.000 |
| | 8 | L.VLDL.P | -0.209 | 0.279 | 0.456 | 0.562 | 1.000 |
| | 9 | XL.VLDL.P | -0.125 | 0.188 | 0.506 | 0.562 | 1.000 |
| | 10 | XS.VLDL.P | 0.238 | 0.423 | 0.574 | 0.574 | 1.000 |
| AF | 1 | L.VLDL.P | -0.417 | 0.250 | 0.097 | 0.945 | 0.968 |
| | 2 | XXL.VLDL.P | 0.157 | 0.140 | 0.263 | 0.945 | 1.000 |
| | 3 | M.VLDL.P | 0.172 | 0.220 | 0.438 | 0.945 | 1.000 |
| | 4 | XL.VLDL.P | -0.093 | 0.157 | 0.556 | 0.945 | 1.000 |
| | 5 | XS.VLDL.P | 0.209 | 0.374 | 0.578 | 0.945 | 1.000 |
| | 6 | IDL.P | -0.448 | 0.902 | 0.620 | 0.945 | 1.000 |
| | 7 | S.VLDL.P | 0.086 | 0.247 | 0.730 | 0.945 | 1.000 |
| | 8 | L.LDL.P | 0.401 | 1.572 | 0.799 | 0.945 | 1.000 |
| | 9 | M.LDL.P | -0.131 | 1.262 | 0.917 | 0.945 | 1.000 |
| | 10 | S.LDL.P | 0.029 | 0.414 | 0.945 | 0.945 | 1.000 |
| CES | 1 | IDL.P | -3.923 | 1.654 | 0.019 | 0.142 | 0.191 |
| | 2 | L.LDL.P | 6.416 | 2.894 | 0.028 | 0.142 | 0.283 |
| | 3 | M.LDL.P | -4.059 | 2.311 | 0.081 | 0.271 | 0.813 |
| | 4 | XS.VLDL.P | 0.928 | 0.673 | 0.170 | 0.426 | 1.000 |
| | 5 | S.VLDL.P | 0.541 | 0.440 | 0.220 | 0.440 | 1.000 |
| | 6 | XXL.VLDL.P | 0.223 | 0.249 | 0.372 | 0.541 | 1.000 |
| | 7 | L.VLDL.P | -0.392 | 0.443 | 0.378 | 0.541 | 1.000 |
| | 8 | M.VLDL.P | -0.264 | 0.391 | 0.500 | 0.625 | 1.000 |
| | 9 | S.LDL.P | 0.406 | 0.742 | 0.585 | 0.650 | 1.000 |
| | 10 | XL.VLDL.P | 0.046 | 0.286 | 0.874 | 0.874 | 1.000 |

**Table S.16. Application example 2 on molecular risk factors for cardiovascular disease outcomes using standard multivariable MR (MV-MR)** which is run on each outcome separately. Results presented include the causal effect estimates, the standard error and the corresponding $p$-value. We adjusted for multiple testing using Benjamini-Hochberg False Discovery Rate (FDR) and Bonferroni correction. For each outcome, we sort the exposures by their $p$-value in increasing order. No exposures are selected at 5% FDR.

| Outcome | Ranking | Exposure | Effect estimate | Standard error | $p$-value | FDR | Bonferroni |
|---|---|---|---|---|---|---|---|
| CAD | | (Intercept) | -0.031 | 0.112 | 0.781 | 0.781 | - |
| | 1 | M.VLDL.P | 0.348 | 0.291 | 0.235 | 0.860 | 1.000 |
| | 2 | S.LDL.P | -0.511 | 0.552 | 0.356 | 0.860 | 1.000 |
| | 3 | M.LDL.P | 1.558 | 1.733 | 0.371 | 0.860 | 1.000 |
| | 4 | S.VLDL.P | -0.235 | 0.341 | 0.493 | 0.860 | 1.000 |
| | 5 | L.LDL.P | -1.121 | 2.217 | 0.614 | 0.860 | 1.000 |
| | 6 | IDL.P | 0.555 | 1.335 | 0.678 | 0.860 | 1.000 |
| | 7 | L.VLDL.P | -0.119 | 0.324 | 0.715 | 0.860 | 1.000 |
| | 8 | XL.VLDL.P | -0.071 | 0.213 | 0.739 | 0.860 | 1.000 |
| | 9 | XXL.VLDL.P | -0.047 | 0.185 | 0.799 | 0.860 | 1.000 |
| | 10 | XS.VLDL.P | 0.100 | 0.568 | 0.860 | 0.860 | 1.000 |
| PAD | | (Intercept) | 0.085 | 0.107 | 0.426 | 0.426 | - |
| | 1 | XS.VLDL.P | 0.767 | 0.473 | 0.107 | 0.615 | 1.000 |
| | 2 | S.LDL.P | -0.724 | 0.535 | 0.179 | 0.615 | 1.000 |
| | 3 | S.VLDL.P | -0.409 | 0.307 | 0.185 | 0.615 | 1.000 |
| | 4 | XL.VLDL.P | -0.232 | 0.210 | 0.270 | 0.675 | 1.000 |
| | 5 | XXL.VLDL.P | 0.174 | 0.185 | 0.349 | 0.698 | 1.000 |
| | 6 | M.LDL.P | 1.162 | 1.668 | 0.487 | 0.769 | 1.000 |
| | 7 | IDL.P | -0.717 | 1.162 | 0.539 | 0.769 | 1.000 |
| | 8 | M.VLDL.P | 0.137 | 0.277 | 0.623 | 0.778 | 1.000 |
| | 9 | L.VLDL.P | 0.078 | 0.309 | 0.801 | 0.890 | 1.000 |
| | 10 | L.LDL.P | 0.015 | 2.071 | 0.994 | 0.994 | 1.000 |
| HF | | (Intercept) | -0.126 | 0.095 | 0.186 | 0.186 | - |
| | 1 | M.LDL.P | 3.673 | 1.474 | 0.014 | 0.139 | 0.139 |
| | 2 | L.LDL.P | -3.961 | 1.855 | 0.035 | 0.173 | 0.346 |
| | 3 | S.VLDL.P | -0.515 | 0.275 | 0.064 | 0.196 | 0.637 |
| | 4 | S.LDL.P | -0.830 | 0.468 | 0.078 | 0.196 | 0.784 |
| | 5 | M.VLDL.P | 0.371 | 0.249 | 0.138 | 0.271 | 1.000 |
| | 6 | IDL.P | 1.471 | 1.047 | 0.163 | 0.271 | 1.000 |
| | 7 | XXL.VLDL.P | 0.215 | 0.165 | 0.195 | 0.279 | 1.000 |
| | 8 | L.VLDL.P | -0.208 | 0.279 | 0.458 | 0.531 | 1.000 |
| | 9 | XL.VLDL.P | -0.133 | 0.188 | 0.478 | 0.531 | 1.000 |
| | 10 | XS.VLDL.P | 0.179 | 0.424 | 0.674 | 0.674 | 1.000 |
| AF | | (Intercept) | 0.070 | 0.085 | 0.417 | 0.417 | - |
| | 1 | L.VLDL.P | -0.415 | 0.250 | 0.099 | 0.987 | 0.994 |
| | 2 | XXL.VLDL.P | 0.156 | 0.140 | 0.266 | 0.987 | 1.000 |
| | 3 | M.VLDL.P | 0.180 | 0.221 | 0.418 | 0.987 | 1.000 |
| | 4 | XS.VLDL.P | 0.232 | 0.376 | 0.538 | 0.987 | 1.000 |
| | 5 | IDL.P | -0.491 | 0.905 | 0.588 | 0.987 | 1.000 |
| | 6 | XL.VLDL.P | -0.085 | 0.157 | 0.592 | 0.987 | 1.000 |
| | 7 | L.LDL.P | 0.442 | 1.575 | 0.780 | 0.989 | 1.000 |
| | 8 | S.VLDL.P | 0.066 | 0.249 | 0.791 | 0.989 | 1.000 |
| | 9 | M.LDL.P | -0.112 | 1.264 | 0.929 | 0.997 | 1.000 |
| | 10 | S.LDL.P | -0.002 | 0.416 | 0.997 | 0.997 | 1.000 |
| CES | | (Intercept) | 0.286 | 0.151 | 0.060 | 0.060 | - |
| | 1 | IDL.P | -4.151 | 1.643 | 0.013 | 0.109 | 0.127 |
| | 2 | L.LDL.P | 6.657 | 2.870 | 0.022 | 0.109 | 0.219 |
| | 3 | M.LDL.P | -4.028 | 2.289 | 0.081 | 0.269 | 0.807 |
| | 4 | XS.VLDL.P | 1.044 | 0.669 | 0.121 | 0.303 | 1.000 |
| | 5 | S.VLDL.P | 0.462 | 0.437 | 0.293 | 0.554 | 1.000 |
| | 6 | XXL.VLDL.P | 0.220 | 0.246 | 0.373 | 0.554 | 1.000 |
| | 7 | L.VLDL.P | -0.381 | 0.439 | 0.388 | 0.554 | 1.000 |
| | 8 | M.VLDL.P | -0.234 | 0.387 | 0.547 | 0.684 | 1.000 |
| | 9 | S.LDL.P | 0.292 | 0.737 | 0.692 | 0.769 | 1.000 |
| | 10 | XL.VLDL.P | 0.072 | 0.284 | 0.801 | 0.801 | 1.000 |

**Table S.17. Application example 2 on molecular risk factors for cardiovascular disease outcomes using multivariable MR-Egger** which is run on each outcome separately. Results include the causal effect estimates as well as the intercept, the standard error and the corresponding $p$-value. We adjust for multiple testing using Benjamini-Hochberg False Discovery Rate (FDR) and Bonferroni correction. For each outcome, we sort the exposures by their $p$-value in increasing order. No molecular exposures are selected at 5% FDR.

| Outcome | Ranking | Exposure | MACE | mPPI | Empirical $p$-value | FDR | Bonferroni |
|---|---|---|---|---|---|---|---|
| CAD | 1 | **L.LDL.P** | 0.300 | 0.689 | $3.00\times10^{-5}$ | $3.00\times10^{-4}$ | $3.00\times10^{-4}$ |
| | 2 | IDL.P | 0.094 | 0.291 | 0.013 | 0.065 | 0.130 |
| | 3 | M.LDL.P | 0.063 | 0.212 | 0.055 | 0.184 | 0.553 |
| | 4 | XS.VLDL.P | 0.010 | 0.093 | 0.491 | 0.999 | 1.000 |
| | 5 | S.LDL.P | -0.011 | 0.091 | 0.846 | 0.999 | 1.000 |
| | 6 | M.VLDL.P | 0.008 | 0.055 | 0.985 | 0.999 | 1.000 |
| | 7 | L.VLDL.P | -0.003 | 0.040 | 0.997 | 0.999 | 1.000 |
| | 8 | S.VLDL.P | 0.004 | 0.049 | 0.997 | 0.999 | 1.000 |
| | 9 | XL.VLDL.P | -0.004 | 0.041 | 0.997 | 0.999 | 1.000 |
| | 10 | XXL.VLDL.P | -0.002 | 0.035 | 0.999 | 0.999 | 1.000 |
| PAD | 1 | **XS.VLDL.P** | 0.192 | 0.840 | $9.00\times10^{-5}$ | 0.001 | 0.001 |
| | 2 | IDL.P | 0.026 | 0.149 | 0.154 | 0.772 | 1.000 |
| | 3 | XL.VLDL.P | -0.015 | 0.106 | 0.888 | 0.995 | 1.000 |
| | 4 | L.LDL.P | 0.004 | 0.063 | 0.938 | 0.995 | 1.000 |
| | 5 | M.LDL.P | 0.004 | 0.054 | 0.972 | 0.995 | 1.000 |
| | 6 | S.VLDL.P | 0.004 | 0.065 | 0.986 | 0.995 | 1.000 |
| | 7 | L.VLDL.P | -0.003 | 0.056 | 0.988 | 0.995 | 1.000 |
| | 8 | S.LDL.P | -0.002 | 0.044 | 0.992 | 0.995 | 1.000 |
| | 9 | M.VLDL.P | 0.003 | 0.049 | 0.994 | 0.995 | 1.000 |
| | 10 | XXL.VLDL.P | -0.001 | 0.049 | 0.995 | 0.995 | 1.000 |
| HF | 1 | **XS.VLDL.P** | 0.190 | 0.881 | $1.00\times10^{-5}$ | $1.00\times10^{-4}$ | $1.00\times10^{-4}$ |
| | 2 | IDL.P | 0.018 | 0.118 | 0.336 | 0.997 | 1.000 |
| | 3 | L.LDL.P | 0.001 | 0.054 | 0.964 | 0.997 | 1.000 |
| | 4 | M.LDL.P | 0.004 | 0.051 | 0.977 | 0.997 | 1.000 |
| | 5 | S.LDL.P | 0.002 | 0.044 | 0.991 | 0.997 | 1.000 |
| | 6 | M.VLDL.P | 0.005 | 0.047 | 0.993 | 0.997 | 1.000 |
| | 7 | XXL.VLDL.P | 0.005 | 0.049 | 0.993 | 0.997 | 1.000 |
| | 8 | XL.VLDL.P | -0.005 | 0.050 | 0.993 | 0.997 | 1.000 |
| | 9 | L.VLDL.P | -0.003 | 0.041 | 0.997 | 0.997 | 1.000 |
| | 10 | S.VLDL.P | 0.001 | 0.050 | 0.997 | 0.997 | 1.000 |
| AF | 1 | **S.VLDL.P** | 0.071 | 0.433 | 0.002 | 0.016 | 0.023 |
| | 2 | **L.VLDL.P** | -0.122 | 0.473 | 0.003 | 0.016 | 0.031 |
| | 3 | XL.VLDL.P | -0.041 | 0.274 | 0.044 | 0.135 | 0.442 |
| | 4 | M.VLDL.P | 0.045 | 0.220 | 0.054 | 0.135 | 0.539 |
| | 5 | XS.VLDL.P | 0.009 | 0.132 | 0.147 | 0.293 | 1.000 |
| | 6 | IDL.P | 0.005 | 0.108 | 0.368 | 0.614 | 1.000 |
| | 7 | L.LDL.P | 0.007 | 0.108 | 0.476 | 0.681 | 1.000 |
| | 8 | M.LDL.P | 0.008 | 0.103 | 0.646 | 0.807 | 1.000 |
| | 9 | S.LDL.P | -0.005 | 0.093 | 0.785 | 0.872 | 1.000 |
| | 10 | XXL.VLDL.P | 0.001 | 0.089 | 0.918 | 0.918 | 1.000 |
| CES | 1 | XXL.VLDL.P | -0.010 | 0.170 | 0.273 | 0.639 | 1.000 |
| | 2 | L.VLDL.P | -0.011 | 0.149 | 0.336 | 0.639 | 1.000 |
| | 3 | S.VLDL.P | 0.006 | 0.109 | 0.369 | 0.639 | 1.000 |
| | 4 | XS.VLDL.P | -0.003 | 0.095 | 0.419 | 0.639 | 1.000 |
| | 5 | IDL.P | -0.005 | 0.100 | 0.461 | 0.639 | 1.000 |
| | 6 | M.VLDL.P | 0.001 | 0.124 | 0.478 | 0.639 | 1.000 |
| | 7 | XL.VLDL.P | -0.003 | 0.139 | 0.509 | 0.639 | 1.000 |
| | 8 | M.LDL.P | 0.006 | 0.104 | 0.615 | 0.639 | 1.000 |
| | 9 | L.LDL.P | 0.000 | 0.099 | 0.618 | 0.639 | 1.000 |
| | 10 | S.LDL.P | -0.004 | 0.102 | 0.639 | 0.639 | 1.000 |

**Table S.18. Application example 2 on molecular risk factors for cardiovascular disease outcomes using the Mendelian randomisation Bayesian model averaging (MR-BMA) algorithm** wich is run on each outcome separately. Results include the model-averaged causal effect estimate (MACE), the marginal posterior probability of inclusion (mPPI) and the corresponding empirical $p$-value [12]. We adjust for multiple testing using Benjamini-Hochberg False Discovery Rate (FDR) and Bonferroni correction. For each outcome, we sort the exposures by their empirical $p$-value in increasing order and the absolute value of MACE in decreasing order. Exposures selected at 5% FDR highlighted in bold.

##### S.4.4 Supplemental Figures

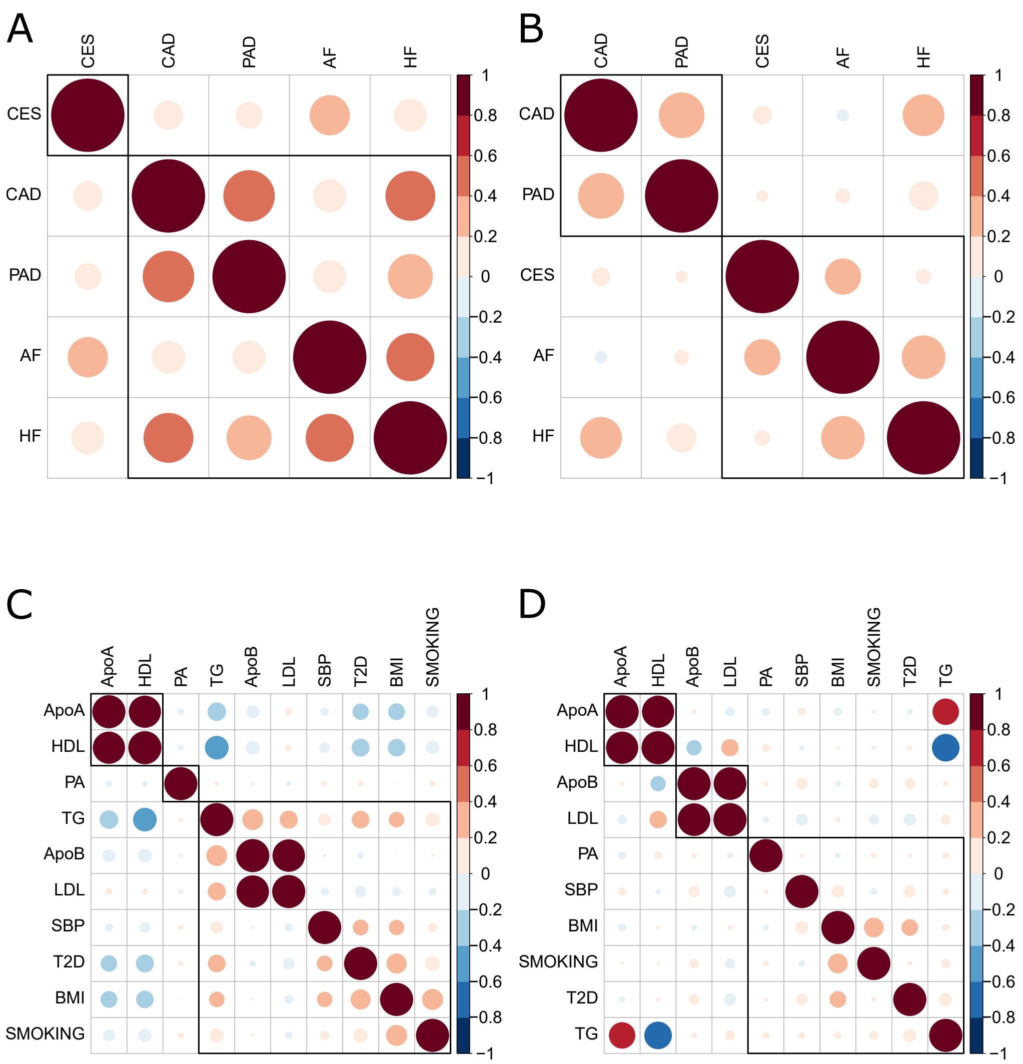


**Figure S.5. Correlation and partial correlation between summary-level outcomes and between summary-level exposures in application example 1 on common risk factors for cardiovascular disease outcomes**. Black boxes indicate clusters obtained by using hierarchical clustering with average agglomeration method, where the optimal number of clusters is estimated using `NbClust` R package. (**A**) Correlation between responses based on 1, 533 independent genetic variants associated with any of the ten exposures after removing seven IVs identified as outliers or high-leverage and influential observations (see Supplemental Figure S.6A). (**B**) Partial correlation between responses based on the same IVs. (**C**) Correlation between summary-level risk factors based on 1, 533 independent genetic variants associated with any of the ten exposures. (**D**) Partial correlation between summary-level risk factors based on the same IVs.

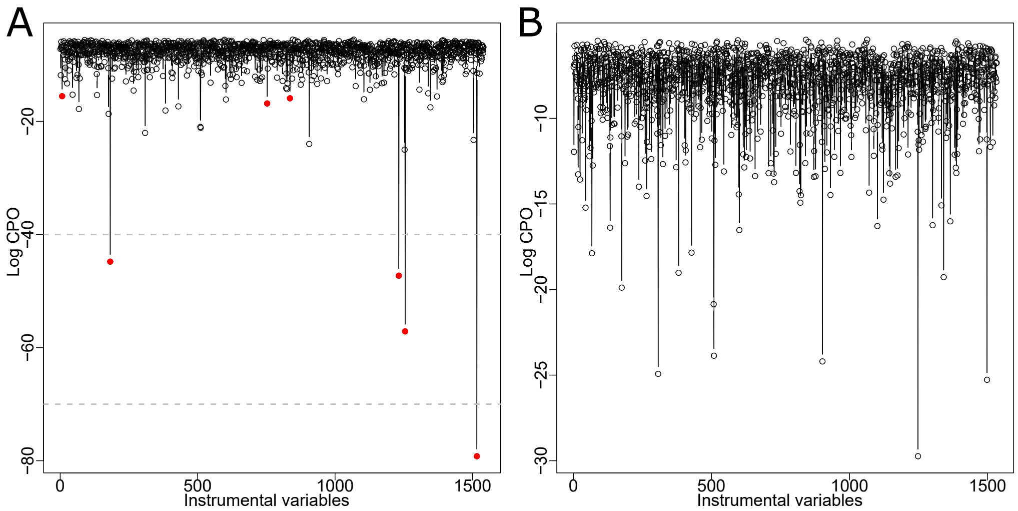


**Figure S.6. Outliers, high-leverage and influential observations detection using the multi-response MR$^2$ model in application example 1 on common risk factors for cardiovascular disease outcomes (CVDs)**, plotting the log-conditional predictive ordinate (CPO) ($y$-axis) against 1.540 instrumental variables ($x$-axis). Red dots indicate observations with scaled CPOs below 0.01 to be regarded as outliers, high-leverage and influential observations [15]. We also show log-inverse-CPOs larger than 40 (possible outliers) and higher than 70 (extreme values) [26]. (**A**) Log-CPO for each instrumental variable and outliers, high-leverage and influential observations detected in the original summary-level data set of common risk factors for CVDs. Seven IVs (0.5%) have scaled CPOs below 0.01, so the MR$^2$ model is considered to fit adequately. (**B**) Log-CPOs after removing outliers, high-leverage and influential observations from the original data set. No outliers or high-leverage and influential observations are detected.

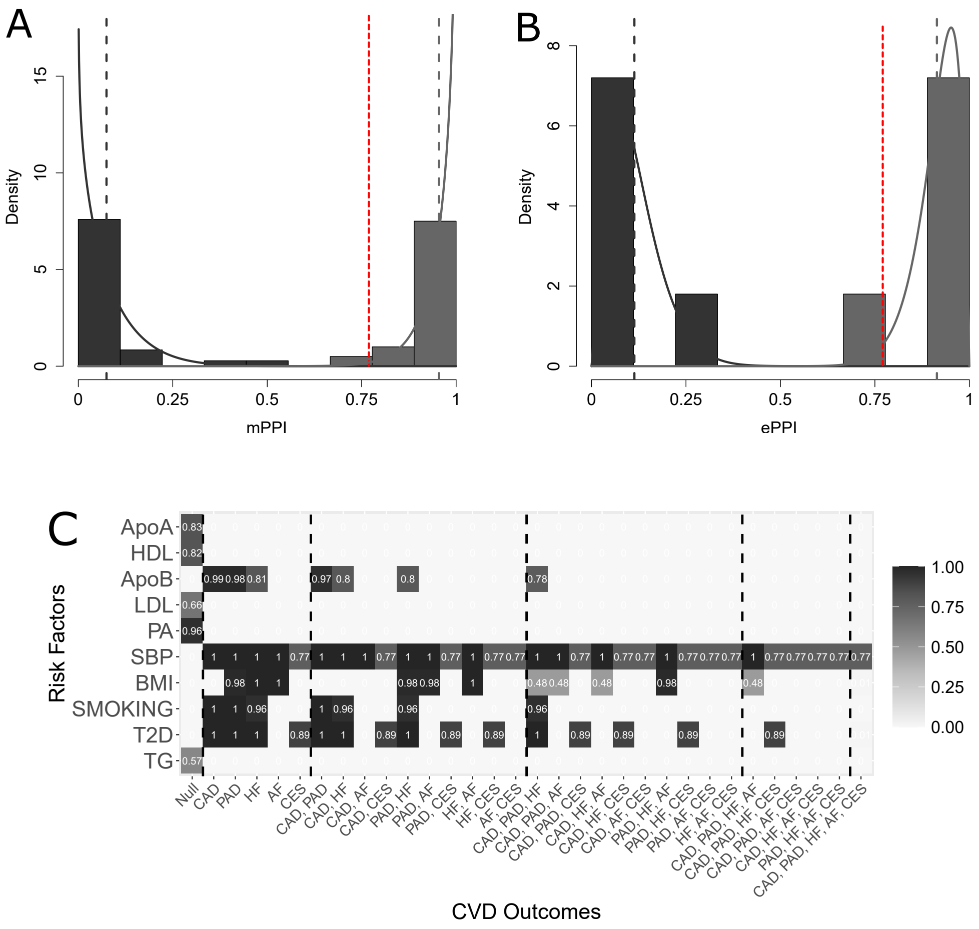


**Figure S.7. Post-processing results of the multi-response MR$^2$ model in application example 1 on common risk factors for cardiovascular disease outcomes.** (**A**) Non-parametric False Discovery Rate (FDR) strategy based on two-component mixture model which clusters low and high levels of the marginal posterior probabilities of inclusion (mPPIs). Dark grey colour code indicates mPPIs and the estimated density of the null ($H_0$) component, while light grey is used for the alternative ($H_1$) component. Dark grey and light grey dotted vertical lines shows the means of the beta components with a good separation between them, while red dotted vertical line specifies the threshold on the mPPI scale (mPPI $\geq 0.77$) that corresponds to 5% FDR nominal level. (**B**) Non-parametric FDR strategy based on two-component mixture model which clusters low and high levels of the edge posterior probabilities of inclusion (ePPIs). Red dotted vertical line specifies the threshold on the ePPI scale (ePPI $\geq 0.78$) that corresponds to 5% FDR nominal level. (**C**) Marginal and joint posterior probability of inclusion (jPPI) of each exposure ($y$-axis) against a single response or combinations of them ($x$-axis). Black dotted vertical lines separate combinations of outcomes with the same group size, *i.e.*, singletons, all possible pairs, all possible triplets, etc. For clarity of presentation, mPPIs for non-selected outcome-exposure pairs (mPPI $< 0.77$ at 5% FDR) are plotted as zero as well as jPPIs for combination of risk factors marginally not selected. The "Null" vertical bar on the left indicates the posterior model probability of the null model.

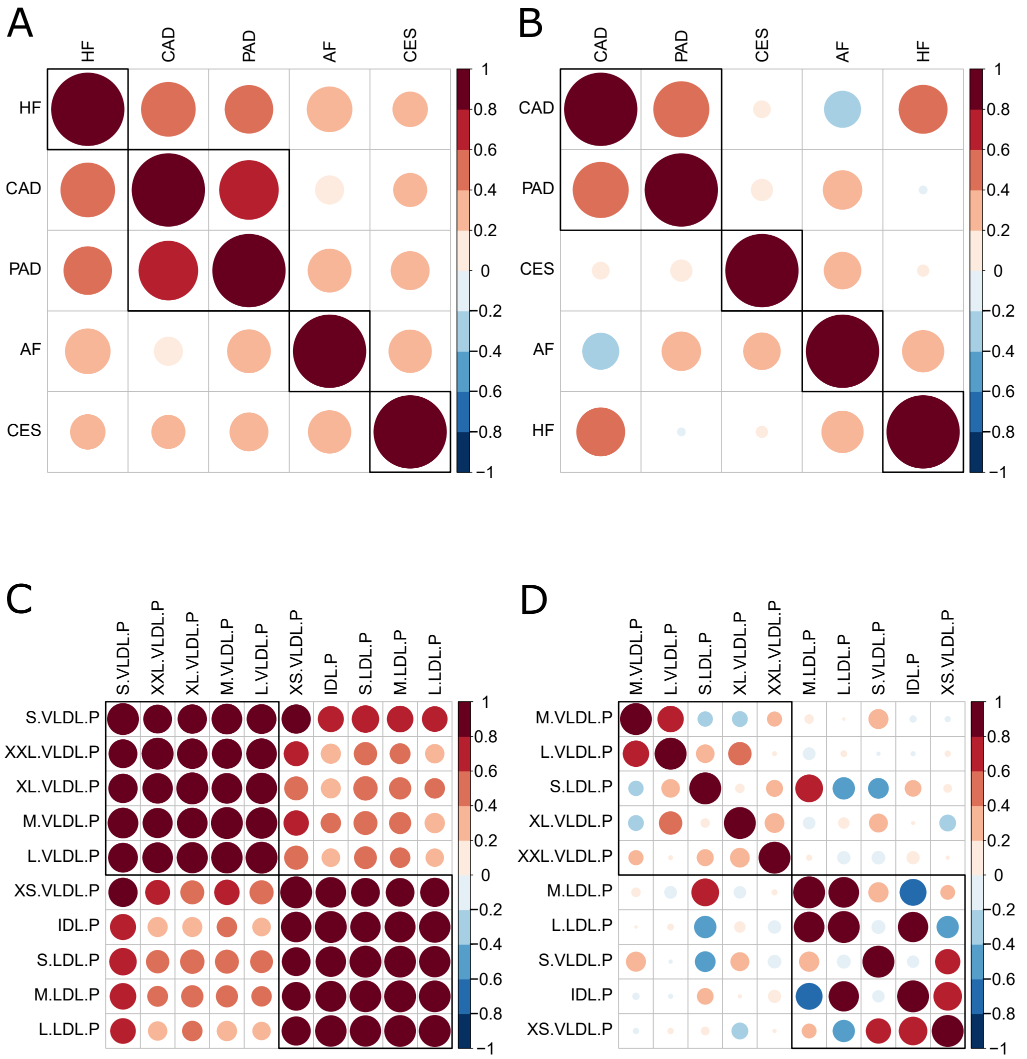


**Figure S.8. Correlation and partial correlation between summary-level outcomes and between summary-level exposures in application example 2 on molecular risk factors for cardiovascular disease outcomes.** Black boxes indicate clusters obtained by using hierarchical clustering with average agglomeration method where the optimal number of clusters was estimated using `NbClust` R package. (**A**) Correlation between responses based on 140 independent genetic variants associated with any of the ten molecular exposures after removing eight IVS identified as outliers or high-leverage and influential observations (see Supplemental Figure S.9A). (**B**) Partial correlation between summary-level responses based on the same IVs. (**C**) Correlation between risk factors based on 140 independent genetic variants associated with any of the ten molecular exposures. (**D**) Partial correlation between summary-level risk factors based on the same IVs.

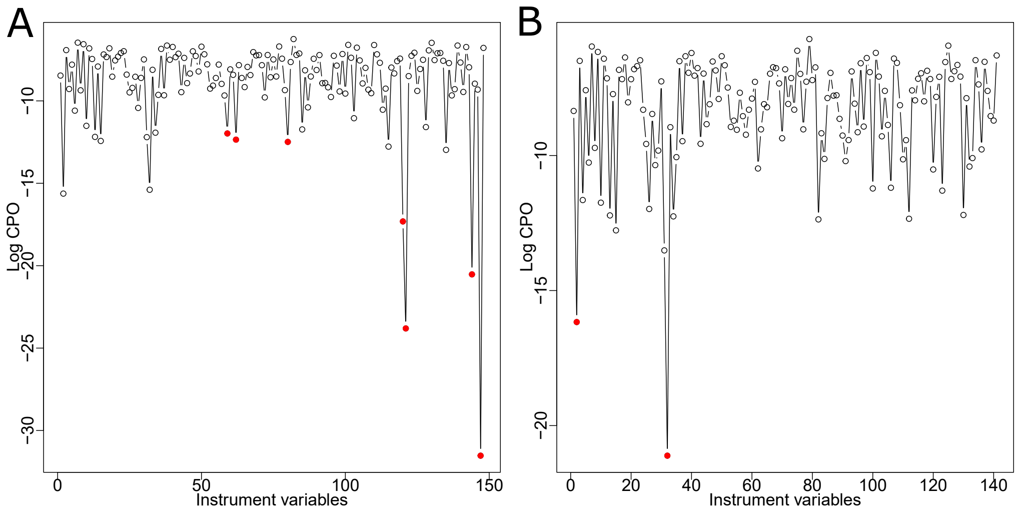


**Figure S.9. Outliers, high-leverage and influential observations detection using the multi-response MR$^2$ model in application example 2 on molecular risk factors for cardiovascular disease outcomes (CVDs)**, plotting the log-conditional predictive ordinate (CPO) ($y$-axis) against 148 instrumental variables ($x$-axis). Red dots indicate observations with scaled CPOs below 0.01 to be regarded as outliers, high-leverage and influential observations [15]. (**A**) Log-CPO for each instrumental variable and outliers, high-leverage and influential observations detected in the original summary-level data set of molecular risk factors for CVDs. Seven IVs (5%) have scaled CPOs below 0.01. (**B**) Log-CPOs after removing outliers, high-leverage and influential observations from the original data set. The MR$^2$ model is now considered to fit adequately since only two outliers or high-leverage and influential observations (2%) are still detected.

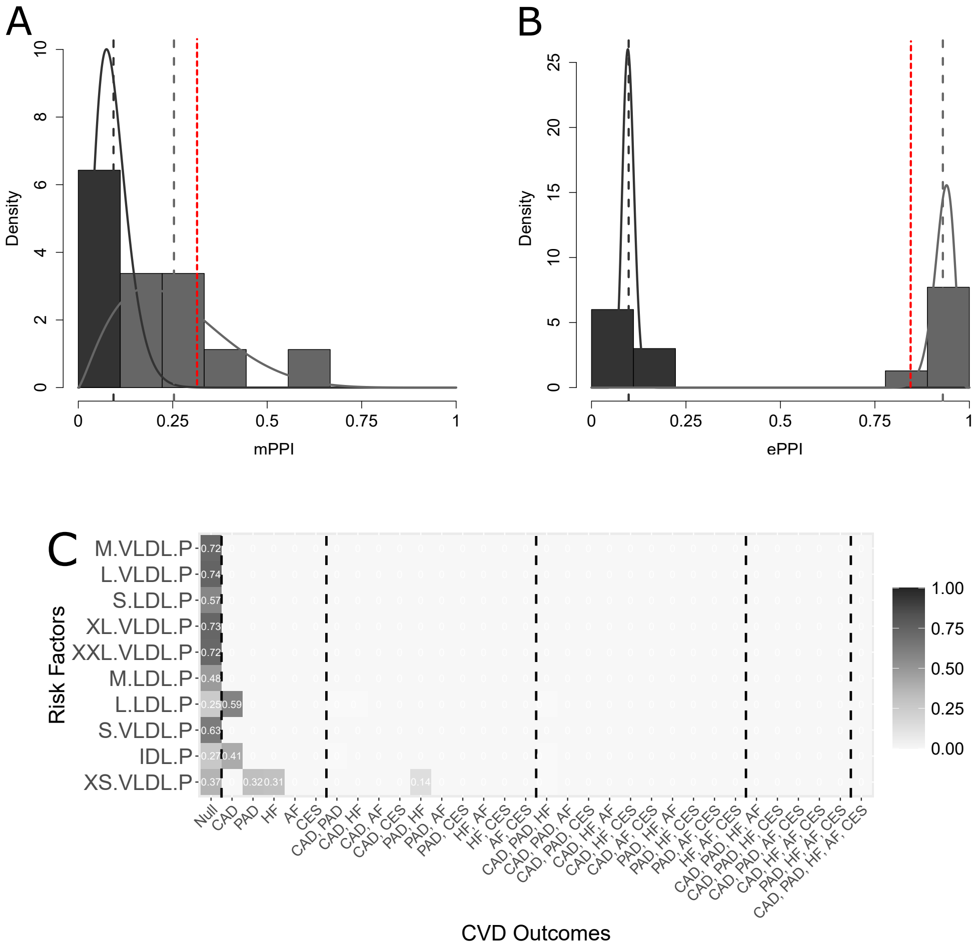


**Figure S.10. Post-processing results of the multi-response MR$^2$ model in application example 2 on molecular risk factors for cardiovascular disease outcomes.** (**A**) Non-parametric False Discovery Rate (FDR) strategy based on two-component mixture model which clusters low and high levels of the marginal posterior probabilities of inclusion (mPPIs). Dark grey colour code indicates mPPIs and the estimated density of the null ($H_0$) component, while light grey is used for the alternative ($H_1$) component. Dark grey and light grey dotted vertical lines shows the means of the beta components, while red dotted vertical line specifies the threshold on the mPPI scale (mPPI $\geq$ 0.29) that corresponds to 5% FDR nominal level. (**B**) Non-parametric FDR strategy based on two-component mixture model which clusters low and high levels of the edge posterior probabilities of inclusion (ePPIs). Red dotted vertical line specifies the threshold on the ePPI scale (ePPI $\geq$ 0.85) that corresponds to 5% FDR nominal level.(**C**) Marginal and joint posterior probability of inclusion (jPPI) of each exposure ($y$-axis) against a single response or combinations of them ($x$-axis). Black dotted vertical lines separate combinations of outcomes with the same group size, *i.e.*, singletons, all possible pairs, all possible triplets, etc. For clarity of presentation, mPPIs for non-selected outcome-exposure pairs (mPPI $<$ 0.29 at 5% FDR) are plotted as zero as well as jPPIs for combination of risk factors marginally not selected. The "Null" vertical bar on the left indicates the posterior model probability of the null model.
